# A standardized, surgically relevant map for emergence of organ-specific branches from the human vagus nerve

**DOI:** 10.64898/2026.05.08.723047

**Authors:** Siyar Bahadir, Frank L Chen, Istvan P Tamas, Elizabeth R McGonagle, Zeinab Nassrallah, Isabelle Pelcher, Joshua Sun, Tiaosi Xing, Michelle Titunick, Shannon Knutson, Todd Levy, Eric H Chang, Robert V Hill, Theodoros P. Zanos, Mary F Barbe, Stavros Zanos

## Abstract

**Introduction:** Vagus nerve stimulation modulates airway, cardiac, pulmonary, and gastrointestinal functions through respective organ-specific branches. Knowledge of where along the nerve vagal branches emerge may support alternative surgical placement of stimulation devices to activate targeted functions while avoiding off-target effects. However, no quantified map of the emergence of vagal branches and their relationship to surgically relevant anatomical landmarks exists in humans.

**Methods:** Fifty-eight vagus nerves (29 left, 29 right) from 29 embalmed donor bodies (15 females) were dissected from the jugular foramen through the thoracic cavity. Branches were traced to end organs and allocated to one of seven groups (sympathetic, muscular, vascular, cardiac, pulmonary, esophageal, or multiple targets) and several sub-groups. Distances between branch emergence and jugular foramen (JF) were normalized to three anatomical landmarks: carotid bifurcation, laryngeal prominence, and superior border of clavicle.

**Results:** Emergence of branches follows a proximal-to-distal order (median [IQR] distance from JF): sympathetic (3.9 cm [3.0–6.6]), muscular (6.2 [3.4–17.0]), vascular (9.6 [5.5–15.1]), cardiac (20.1 [16.9–22.5]), pulmonary (25.0 [22.8–27.6]), and esophageal (26.5 [24.3–29.1]). Branches emerge into two embryological domains separated at the clavicle: pharyngeal arch branches cluster proximally (7.3 cm [3.9–14.0]) and primitive mediastinum branches cluster distally (24.2 cm [20.9–27.1]), with sympathetic, muscular, and vascular sub-groups occupying distinct zones within the proximal domain. The longest branch-free intervals are all left-sided, occurring around the laryngeal prominence (3.0 cm [1.8–5.7]), the superior border of the clavicle (2.7 cm [1.3– 5.2]), and the carotid bifurcation (2.2 cm [0.9–5.7]). Several “target” branches are maximally separated from “off-target” branches at specific distances: sympathetic vs. cervical/ muscular/ vascular at 6/8 cm (L/R), cardiac vs. carotid sinus at 14/10 cm, and recurrent laryngeal vs. other muscular at 18/13 cm. Minimal differences are found between male and female donors.

**Conclusion:** This study describes a quantified, landmark-registered map of cervical and thoracic vagal branch emergence and provides an anatomical reference that may help guide more selective device design and placement.

## Introduction

The vagus nerve innervates most internal organs, including the larynx and pharynx, major vessels, heart, lungs, esophagus and abdominal viscera. Inside the vagus nerve, fibers convey sensory and motor signals that regulate homeostatic functions ranging from airway protection, cardiovascular control, respiration, and gastrointestinal functions (Prescott and Liberles 2022; Kupari et al. 2019; Berthoud and Neuhuber 2000). This functional diversity is mediated anatomically by discrete branches that route organ-specific fibers to their respective target organs or plexuses (Prescott and Liberles 2022; Kupari et al. 2019; Berthoud and Neuhuber 2000; N. Thompson et al. 2019; Kronsteiner B et al. 2024; Thompson N et al. 2026; Na et al. 2025; Jayaprakash N et al. 2023).

Vagus nerve stimulation (VNS) devices are typically implanted in the mid-cervical region, where most vagal fibers converge (Reid 1990; Giordano F et al. 2017). With such placement, VNS activates vagal fibers relatively non-selectively, sometimes leading to off-target effects which may limit therapeutic efficacy. For that reason, strategies for more organ-selective vagal neuromodulation are explored (U. Ahmed et al. 2022). When a stimulation device is implanted at a certain level on the nerve trunk, responses mediated by branches emerging distally to device placement are activated, whereas those mediated by branches emerging proximally are spared (McGee and Grill 2014; Schiefer et al. 2010). Knowledge of the pattern of emergence of organ-specific branches would provide an anatomical template for which organ-specific responses would be expected after placing VNS devices at different levels along the vagal trunk. Additionally, defining patterns of branch emergence relative to surgically identifiable anatomical landmarks may assist in surgical approaches and alternative device placements for more selective vagal neuromodulation.

Several detailed anatomical studies have described patterns of emergence of vagal branches in humans, including laryngeal (Cakir BO et al. 2006; A. Patra 2023), carotid and other vascular (Toorop 2009; Kikuta 2019; Hammer N et al. 2018), sympathetic chain-communicating (Sato et al. 1997; Kawagishi et al. 2008; Seki et al. 2014), cardiac (Kawashima 2005; Kronsteiner B et al. 2024), pulmonary (Weijs et al. 2015), and distal esophagogastric/foregut plexus branches (Toorop 2009; Weijs et al. 2015, 2017; Parpex et al. 2020). However, there are no studies covering the full range of vagal branches, identified in the same nerves and subjects; likewise, no anatomical studies have linked branch emergence with surgically identifiable landmarks such as the clavicle, carotid bifurcation, or laryngeal prominence.

In this study, we mapped emergence of branches from the human vagal trunk on a standardized template and performed a statistical analysis of branch emergence patterns for left and right nerves, in males and females. Our objectives were to: 1) identify all vagal branches from the upper cervical through the thoracic region and group them into organ-specific groups and subgroups; 2) measure distances between branch emergence and the jugular foramen, and normalize these distances with respect to surgically identifiable anatomical landmarks; 3) quantify prevalence of different branch groups, as well as branch-free regions on the vagal trunk, and assess differences between left and right nerves, and between male and female donors; 4) compare level of emergence of branches mediating some targeted vs. off-target effects and explore potential implications for alternative placement of devices for more selective vagal neuromodulation.

## Methods

### Donors and nerves

All post-mortem study materials used in this work came from direct donations to the Anatomical Gift Programs of the Zucker School of Medicine, SUNY Upstate Medical University, and Lewis Katz School of Medicine of Temple University. The study was conducted under the relevant policies of these institutions. Donor demographic data were recorded from death certificate information available when each donor body was received by the institution. The use of human donor bodies in this study falls under IRB Exempt Category 4.ii for research, as defined by the US Department of Health and Human Services and NIH regulations, 45 CFR 46.104.

Twenty-nine human donor bodies were prepared by intravascular embalming through the femoral artery using an anatomical solution from Embalmers’ Supply Company, East Lyme, CT. This preparation was selected to preserve the cervical vagus and minimize procedural disturbance of the nerve. Donor bodies with major medical histories involving cancer or prior procedures in the neck or thorax were not included. Bilateral dissections were performed to identify and characterize vagal anatomy in 16 female and 13 male donor bodies, yielding 58 nerves total, including 29 left and 29 right sides. None of the donor bodies had undergone previous anatomical dissection. In accordance with HIPAA protected health information Safe Harbor guidelines, 164.514(b)(2), ages greater than 90 years were reported as 90+.

One donor was excluded from further analysis due to unusually highly branched vagus nerves and our inability to reliably trace most branches to their target endpoints and confidently assign them to groups and subgroups. The analyzed dataset comprised 28 donors (15 female, 13 male; median age 88 years, range 57–90+) and 56 vagus nerves (28 left, 28 right).

### Anatomical dissection

Anatomical dissection started with the cervical region. The vagus nerve and its targets were isolated, bilaterally, in the cervical region. After disarticulation of craniovertebral junction, and hyperflexion of the head, the vagus nerve within carotid sheath at the parapharyngeal space was exposed. Neck and appropriate thorax dissections were performed along with the disarticulation to expose the vagus from the jugular foramen through the thoracic cavity.

Due to the potential convergence of the vagus nerve with the accessory, glossopharyngeal and hypoglossal cranial nerves in the most superior cervical region (which we termed the cranial portion of the vagus nerve), during the dissections, we focused our examinations on the cervical portion of the vagus nerve, defined as the portion that is distal to the emergence of the superior laryngeal nerve and proximal to superior border of clavicle. Detailed dissection information is provided in the Supplementary Methods of the Supplementary files.

After the vagus nerve was exposed, sutures were placed bilaterally on the epineurium of the vagus nerve at the level of the jugular foramen, superior laryngeal prominence, carotid bifurcation, and superior border of the clavicle as landmark indicators. The location of landmarks, point of emergence of each branch from the vagus nerve relative to the jugular foramen, and number of branches per vagus nerve was documented. We systematically traced each branch of the vagus to their target endpoint(s) and documented if a branch innervated more than one target (termed as multiple targets). Sutures of various colors and sizes, and tissue marking dyes (22-050-455, FisherScientific, Epredia™ Mark-It™ Tissue Marking Dye kit, Richard-Allan Scientific LLC (a subsidiary of Epredia), Kalamazoo, MI, USA) were used as fiduciary markers for the various subsets of vagal nerve branches. After marking, the vagus nerve and its branches were video recorded and photo-documented, both prior to and after extraction and sectioning into 1cm partitions.

A detailed protocol on anatomical dissection can be found under: https://dx.doi.org/10.17504/protocols.io.14egn683ml5d/v1

### Measurement of branch emergence distance

After dissection and extraction, each vagus nerve was laid out in a standardized orientation and segmented using the same set of anatomical landmarks. Branch emergence distances were measured manually with a metric ruler along the course of the main trunk (centerline/ along-nerve path), from the nearest proximal landmark reference point to the branch takeoff (defined as the first visible divergence of the branch from the parent trunk). Measurements were recorded and linked to the branch ID and target endpoint classification. These ruler-based distances were cross-checked against micro-CT derived measurements from scans of the same nerves. When discrepancies were identified, branch or landmark positions were re-evaluated and the recorded values were updated to reflect the micro-CT based measurement (i.e., micro-CT served as the adjudicating reference for select corrected entries).

### Partitioning and standardization of branches according to anatomical landmarks

For landmark-based partitioning, the vagus was divided into four segments defined by fixed anatomical boundaries (landmarks): upper cervical (jugular foramen to carotid bifurcation), mid-cervical (carotid bifurcation to laryngeal prominence), lower cervical (laryngeal prominence to superior border of the clavicle), and thoracic (distal to the clavicle). All analyses and interpretations refer to these landmark-defined segments. Prevalence of branches by segment and branch density (branches per centimeter, and per nerve) were determined and reported (Table 1).

**Table 1.** Overview of donors, nerves, branches and anatomical reference landmarks.

|  | **Total** | **Left (L)** | **Right (R)** |
| --- | --- | --- | --- |
| **Donors** | 28 (15F /13M),<br>median age 88<br>(range: 57–90+) | — | — |
| **Nerves** | 56 | 28 | 28 |
| **Branches** | 2,177 | 997 | 1,180 |
| **Total nerve length, mean**<br>**(range), in cm** | 32.6 (24.5–44.7) | 32.6 (24.5–44.0) | 32.6 (26.0–44.7) |
| **Anatomical landmarks** |  |  |  |
| **Carotid bifurcation, native**<br>**distance from JF ($D_N$)** | 6.2 (2.7–10.1) | 6.3 (2.7–10.1) | 6.1 (2.9–10.0) |
| **Laryngeal prominence, $D_N$** | 8.6 (5.4–12.1) | 8.7 (6.0–11.4) | 8.5 (5.4–12.1) |
| **Superior border of clavicle,**<br>**$D_N$** | 15.6 (9.6–22.5) | 15.7 (12.0–19.8) | 15.5 (9.6–22.5) |
| **Average branch emergence,**<br>**$D_N$** | 17.5 (0.2–43.5) | 17.3 (0.5–43.5) | 17.6 (0.2–41.8) |
| **Average branch emergence,**<br>**scaled distance from JF** | 17.4 (0.3–32.6) | 17.2 (0.5–32.6) | 17.5 (0.3–32.6) |
| **Branches by segment (%)** |  |  |  |
| **Upper cervical** | 18.1 | 20.1 | 16.5 |
| **Mid cervical** | 6.8 | 6.7 | 6.9 |
| **Lower cervical** | 14.0 | 16.0 | 12.2 |
| Thoracic | 61.1 | 57.2 | 64.4 |
| **Branch density (branches/cm/nerve)** |  |  |  |
| Upper cervical | 1.07 | 1.05 | 1.09 |
| Mid cervical | 0.92 | 0.91 | 0.93 |
| Lower cervical | 0.76 | 0.80 | 0.71 |
| Thoracic | 1.44 | 1.23 | 1.65 |

F = Female, M = male, JF = jugular foramen.

Each nerve was mapped to a standardized longitudinal axis to compare branch emergence locations across donors with different trunk lengths and inter-landmark distances. Registration was performed separately for left and right vagus nerves. The distal endpoint for each nerve was defined as the true trunk length, computed as the sum of sequential segment lengths along the main vagal trunk, including branch-free sections. Occasional inconsistencies were observed in the relative ordering of the carotid bifurcation and laryngeal prominence (change in order and being on the same position) (shown in Supplementary Table S1). The registration transform itself is implemented in an order-based manner for robustness: the three landmark coordinates on each nerve side were sorted and treated as uppermost, mid, and lowermost fiducials, and a linear scaling is applied to each individual nerve over each of the segments defined by the mean positions of landmarks above. This way, all segments were scaled to the mean distance of that segment, for that nerve. For rare ties (carotid and laryngeal prominence at the same coordinate), the mid fiducial is offset by ε = 0.10 cm to avoid a zero-length interval. The donors with higher laryngeal prominence than carotid bifurcation and the ones with those two structures at the same exact level are reported in Supplementary Table S1.

In practical terms, each nerve is normalized to a common template by scaling (stretching or compressing) its inter-landmark segments to a shared, cohort-average length. Because every donor’s landmarks are mapped to the same registered positions, a branch is characterized not by its actual positions on that nerve but by its scaled position within the segment it occupies, relative to the average length of that segment (Supplementary Figure 10). A branch emerging midway between the laryngeal prominence and the clavicle receives the same registered distance whether the nerve is long or short, even though the native distances differ. This removes between-donor differences in total trunk length and in landmark spacing, which allows pooling of branch positions and comparisons between donors. The effect of the registration procedure is illustrated in Supplementary Figure 10, which contrasts emergence positions of branches from two short (B825) and two long (B923) vagus nerves in raw and registered coordinates. The transformation preserves the order of branches along the nerve, and each branch’s position relative to the landmarks. Registration expresses branch position in landmark-relative terms that are comparable across individuals, rather than in raw distances that can conflate nerves of different size.

### Partitioning of branches according to innervated organ

Each branch emergence was defined as a discrete emergence point from the main vagal trunk (first visible divergence of a branch) and was categorized based on the innervated organ documented during dissection. Branches were assigned to one of seven main groups: Sympathetic, Muscular, Vascular, Cardiac, Pulmonary, Esophageal, or Multiple Targets. The Multiple Targets group was not included in the figures and analyses due to low prevalence in both group and subgroup level. Branches within each group were further divided in one of several subgroups, depending on target or plexus. In the Sympathetic group, subgroups include: superior cervical ganglion, sympathetic trunk, and branches that target both the sympathetic trunk and superior cervical ganglion; examples are shown in Figure 1, A and Supplementary Figure 1. In the Vascular group, subgroups include branches that directly innervate the carotid sinus, internal carotid artery, external carotid artery, carotid bifurcation, branches that innervate multiple vascular targets (termed “Multiple branches to the carotid artery”), internal jugular vein, common carotid, aorta, and general vascular, used when a branch is confidently vascular but no more specific target could be assigned; examples are shown in Figure 1 B1, B2, B3 and B4 and Supplementary Figure 4. In the Cardiac group, subgroups include: cardiac plexus, superficial cardiac plexus, general cardiac (branches directed toward the heart for which the precise endpoint could not be reliably identified), deep cardiac plexus, and cardiopulmonary; examples are shown in Figure 1 D1, D2, D3 and D4 and D5 and Supplementary Figure 5 and 6. In the Muscular group, subgroups include pharyngeal, superior laryngeal nerve (SLN), ansa cervicalis, general laryngeal (branches from the vagus that directly innervated laryngeal muscles), non-recurrent laryngeal nerve, and recurrent laryngeal nerve (RLN); examples are shown in Figure1 C1, C2, C3, C4, and C5, and Supplementary Figures 2 and 3. The Pulmonary group has no subgroup, as medial and lateral pulmonary branches were combined for this study. In the Esophageal group, subgroups include general esophageal branches (defined as a branch to esophagus, but not to the esophageal plexus), organ-specific branches that arose from the esophageal plexus (pulmonary, cardiac, cardiopulmonary), as well as branches to the esophageal plexus. Pulmonary and esophageal branches are shown in with examples in Figure 1 D5 and Supplementary Figure 6. When a single branch bifurcated to supply more than one distinct target group or subgroup, the branches were assigned to the Multiple Targets group and labeled by its mixed-endpoint subgroup (e.g., vascular and superficial cardiac plexus; vascular and sympathetic chain; vascular and laryngeal, vascular and pharyngeal; vascular and muscular – laryngeal; sympathetic chain, cardiovascular, and pulmonary; sympathetic chain, cardiovascular, and laryngeal muscles). This group was not included in statistical analysis however are used in visualization. Group- and subgroup-level summaries are reported as totals and stratified by side (Left vs Right) and sex (Female vs Male) where indicated; side comparisons are treated as within-donor (paired) observations.

**Figure 1.**
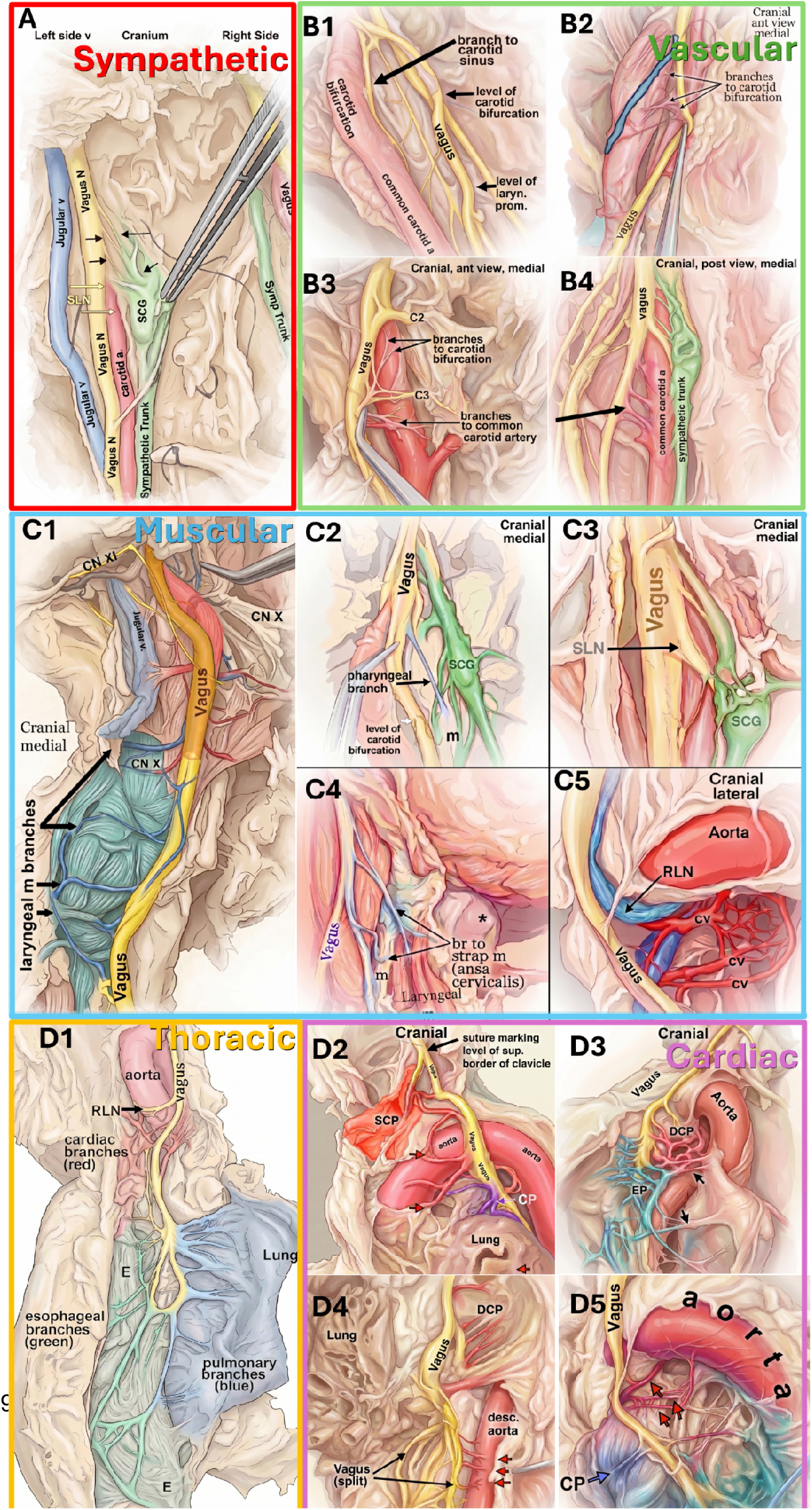
Identification of vagal branches belonging to specific groups during anatomical dissection. Individual pictures are grouped together and framed colored borders and panel titles indicating the branch group: red, sympathetic (A); green, vascular (B1–B4); blue, muscular (C1– C5); purple, cardiac (D2, D3); orange, thoracic overview and pulmonary/esophageal (D1, D4, D5). The vagus is dyed yellow throughout; other labelled structures are shown for orientation. (A) Sympathetic: vagus-to-superior cervical ganglion (SCG) branches and the sympathetic trunk, the superior laryngeal nerve (SLN), jugular vein, and carotid artery are labelled for orientation. (B) Vascular: (B1) branch to the carotid sinus at the carotid bifurcation; (B2) branches to the carotid bifurcation; (B3) branches to the carotid bifurcation and common carotid artery (C2/C3 roots for orientation); (B4) branches to the common carotid artery (sympathetic trunk for orientation). (C) Muscular: (C1) laryngeal-muscle branches (spinal accessory nerve, CN XI, and adjacent cranial nerve labelled for orientation); (C2) pharyngeal-muscle branch (SCG for orientation); (C3) origin of the superior laryngeal nerve (SLN); (C4) branches to the strap muscles via the ansa cervicalis; (C5) origin of the recurrent laryngeal nerve (RLN) at the aortic arch (cardiovascular branches(cv), and aorta for orientation). (D) Thoracic: (D1) overview of the thoracic vagus with cardiac, esophageal, and pulmonary branches; (D2) superficial cardiac plexus (SCP) and cardiopulmonary (CP) branches at the level of the clavicle; (D3) deep cardiac plexus (DCP) branches (esophageal plexus, EP, for orientation); (D4) deep cardiac plexus branches near the descending aorta, where the vagus splits into trunks; (D5) cardiopulmonary branches near the aorta. Abbreviations: CN XI, spinal accessory nerve; CP, cardiopulmonary; cv, cardiovascular branch; DCP, deep cardiac plexus; EP, esophageal plexus; RLN, recurrent laryngeal nerve; SCG, superior cervical ganglion; SCP, superficial cardiac plexus; SLN, superior laryngeal nerve.

A secondary topographic partitioning aggregated related branch groups into two broader longitudinal domains. The Pharyngeal Arch Target-related domain (cervical domain) comprised the sympathetic, muscular, and vascular groups — including the recurrent laryngeal and aortic branches — and the Primitive Mediastinum Target-related domain (thoracic domain) comprised the cardiac, pulmonary, and esophageal groups. The two domains correspond to the pharyngeal-arch–derived cervical territory and the mediastinal/visceral thoracic territory, respectively, separated at the superior border of the clavicle. The cervical domain was further divided into three longitudinal zones superior cervical, mid/low cervical, and upper thoracic reflecting the separable proximal, intermediate, and distal emergence regions of its muscular and vascular targets. These domains and zones did not replace the original group/subgroup taxonomy; they were used only as an additional analysis of regional topography along the vagus. The developmental and evolutionary rationale for this grouping is detailed in Supplementary Methods.

## Statistical Analysis

### Descriptive Statistics

All statistical analyses are performed in Python using pandas, numpy, scipy, matplotlib, and seaborn libraries. Descriptive statistics include the number of branches, the number of contributing donors, median distance of emergence and interquartile range, standardized effect sizes (difference in least square means), linear discriminant analysis separation thresholds with 95% bootstrap confidence intervals, mean proximal and distal branch-free boundaries with corresponding standard deviations around anatomical landmarks, and anatomical separation summaries including the mean i(x) curve, standard deviation across comparison curves, and the peak distance x*. Results are reported for the pooled dataset and stratified by side. Sex stratification is used only when explicitly noted.

### Emergence distance comparisons

Distributions of emergence distance from the jugular foramen are visualized as violin plots with overlaid branch emergence. Two types of comparisons are performed. The first compares emergence distance in left vs. right nerves, for each subgroup; for each subgroup, paired Wilcoxon signed-rank tests are applied across donors, requiring at least six donors. The second compares emergence distance between selected subgroups, to test whether one subgroup emerges proximally or distally compared to another; for each pair of subgroups, paired Wilcoxon signed-rank tests were performed, requiring at least six donors. For male-female comparisons, Mann-Whitney-u test was used. In each case, Bonferroni correction for multiple comparison was applied. Only significant results after correction are annotated in figures. Standardized effect sizes are summarized using difference between least square means.

### Linear mixed-effects models

To use all branch-level observations while correcting for possible correlations between branches from the same donors, we compiled restricted maximum likelihood, linear mixed-effects models. For each group of branches, registered emergence distance was modeled as dependent variable, with subgroup, side, and sex as fixed effects, their two-way interactions (sex × side, sex × subgroup, side × subgroup), and an intercept for donor as random effect, to account for within-donor correlations. Model assumptions and convergence were checked. Least-squares (adjusted) subgroup means, averaged over side and sex, are reported with 95% confidence intervals (Table 3); fixed-effect beta estimates with 95% confidence limits are in Supplementary Tables S2–S6; and the between-donor variance share is summarized by the intraclass correlation coefficient (ICC). Between-subgroup differences were evaluated as linear contrasts of the fixed-effect estimates, with Bonferroni correction for multiple comparisons. Analyses were performed using Python (statsmodels).

### Cervical-thoracic transition estimation

To estimate the transition from cervical-domain targets to thoracic-domain targets along the trunk (Figure 7A), we used one-dimensional linear discriminant analysis with distance from the jugular foramen as the only input, separately for left and right nerves, and male and female donors. This method treats the two branch groups as overlapping distance distributions along a single axis, estimates their central tendency and spread, and identifies a single cut point where a branch becomes more likely to belong to one class than the other, based on the observed class frequencies. This cut point is reported as the transition threshold. Uncertainty is quantified by bootstrap resampling of branch events within each class, producing a distribution of thresholds from which a ninety five percent confidence interval is derived. We additionally quantified how well this boundary classifies branches — accuracy, sensitivity, specificity, and the area under the ROC curve — treating thoracic (primitive-mediastinum) branches as the positive class, so that sensitivity is the fraction of thoracic branches and specificity the fraction of cervical branches correctly classified. These metrics were computed separately for the left and right nerves and are reported in Supplementary Table S.

### Branch-free intervals

For the branch-free interval analysis (Figure 7), we quantified the uninterrupted vagus trunk surrounding three fixed anatomical landmarks—the carotid bifurcation, laryngeal prominence, and superior border of the clavicle—by identifying, for each donor and side, the nearest branch proximal and the nearest branch distal to each landmark along the landmark-registered axis. These donor-level intervals are plotted separately for the left and right vagus, with each horizontal line representing the branch-free interval centered on a given landmark. Donors are ordered according to the largest branch-free interval observed on the left side to facilitate side-by-side visual comparison, and study-population-level summaries are overlaid as mean proximal and distal boundaries with corresponding standard deviations for each landmark (Figure 7, Supplementary Table 6).

### Anatomical separation between target vs. off-target branches

We performed a distance-based anatomical separation analysis along the vagal trunk. We computed an anatomical separation index i(x) that captures the probability that a selected set of target (T) vs. another set of off-target (OT) branches, emerge distally to a level of the vagal trunk at distance x from JF.

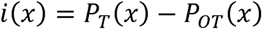

Positive values of i(x) indicate that at that level, targeted branches are more likely to arise distally compared to off-target branches. The greater value the value of i(x), the bigger the difference between probabilities of targeted vs. off-target branch emergence; for example, i(x) of 70% indicates that there is a 70% higher probability that targeted branches will emerge distally to the given level of the vagal trunk vs. off-target branches. The exact value of i(x) depends on the level of the vagal trunk (x) and the level at which i(x) becomes maximal is called peak anatomical separation i(x*) for that specific target vs. off-target contrast. Importantly, this index reflects relative branch separations only and does not indicate fiber activation or functional selectivity.

Throughout, ’selectivity’ refers to spatially selective surgical placement based on separation of branch emergence, not validated functional or electrophysiological selectivity. Also, the map is a gross anatomical proxy and does not imply fiber-level separation.

### Interactive atlas explorer

We developed a browser-based interactive atlas explorer using Python and JavaScript to allow readers to examine the vagus nerve branching atlas beyond the static manuscript figures. The application displays branch emergence along the landmark-registered vagus axis and allows users to filter the atlas by side, sex, donor, branch group, and branch subgroup. It includes interactive views of individual branch events, donor-level subgroup prevalence, group/subgroup distributions, and distance-based anatomical separation. Users can also define custom target and comparator branch populations to explore where along the vagus trunk a candidate stimulation level may enrich desired organ- or system-related branches while reducing expected off-target branch populations. In this way, the tool supports anatomical inspection, translation of the atlas into operative and device-placement questions, and exploratory hypothesis generation using the same branch classification and landmark-registered framework described in the manuscript.

The explorer is a self-contained, client-side browser application (HTML and JavaScript) that operates on a single structured dataset, with no server or database backend. The dataset is organized as one observation per branch, with fields for donor identifier, side, sex, functional group, subgroup, traced target, native distance from the jugular foramen, landmark-registered distance, and the donor’s landmark positions; landmark observations are stored in the same schema. Upon loading, the application reads the dataset and builds four linked views: individual-branch, donor-level prevalence (“subway map”), group and subgroup distributions, and distance-based anatomical separation. User can select filters like side, sex, donor, group, and subgroup, which are applied to the dataset, and all views are updated accordingly. The anatomical separation view is updated after the user defines a “target” set of branches and an “off-target” set of branches; then, for different positions along the registered axis, the anatomical separation index i(x) is calculated, as the difference between the probability of target branches and the probability of off-target branches emerging distal to x; the peak of i(x) function marks the level of maximal anatomical separation.

All computation runs in the browser on the flat data file, and the complete dataset, the figure-generating code, and a reproducible notebook that regenerates every figure and table in this manuscript from that same dataset are provided in the project repository (https://github.com/drsiyarb/vagus_nerve_explorer). To ensure long-term accessibility, an archived snapshot of the dataset and code has been deposited on Zenodo with a persistent DOI (https://doi.org/10.5281/zenodo.21268191) (Bahadır 2026). Because the tool is a static client-side application operating on a single flat data file, it can be downloaded and run locally or re-hosted without reliance on GitHub or any backend service. The tool is also currently available through SPARC portal (https://sparc.science/tools-and-resources/4dhLyABA6FBJo1nFcJdRKm)

## Results

### 1. Interactive online tool for visualization and analysis of emergence of vagal branches

One of the outputs of our work is the development of an interactive, online tool for visualizing and statistically analyzing emergence of different groups of vagal branches, in single donors or in groups of donors (Figure 2A). Users can interrogate the dataset through four linked visualization modes. The individual-branch view allows selected donor nerves, sides, sexes, groups, or subgroups to be displayed along the landmark-registered vagus axis (Figure 2B). The prevalence, or subway-map, view summarizes branch subgroups by left-only, right-only, bilateral, and total donor-level prevalence (Figure 2C). The distribution view allows selected branch populations to be compared across left and right nerves, male and female donors, individual donors, or custom-defined groups and subgroups (Figure 2D). The relative emergence view allows the comparison of relative emergence of selected targeted and off-target branches to estimate how placement of a device at different levels along the vagal trunk could potentially change the relative engagement of branches (Figure 2E). Throughout the views, the tool references branch emergence to surgically recognizable landmarks such as the carotid bifurcation, laryngeal prominence, and superior border of the clavicle.

**Figure 2.**
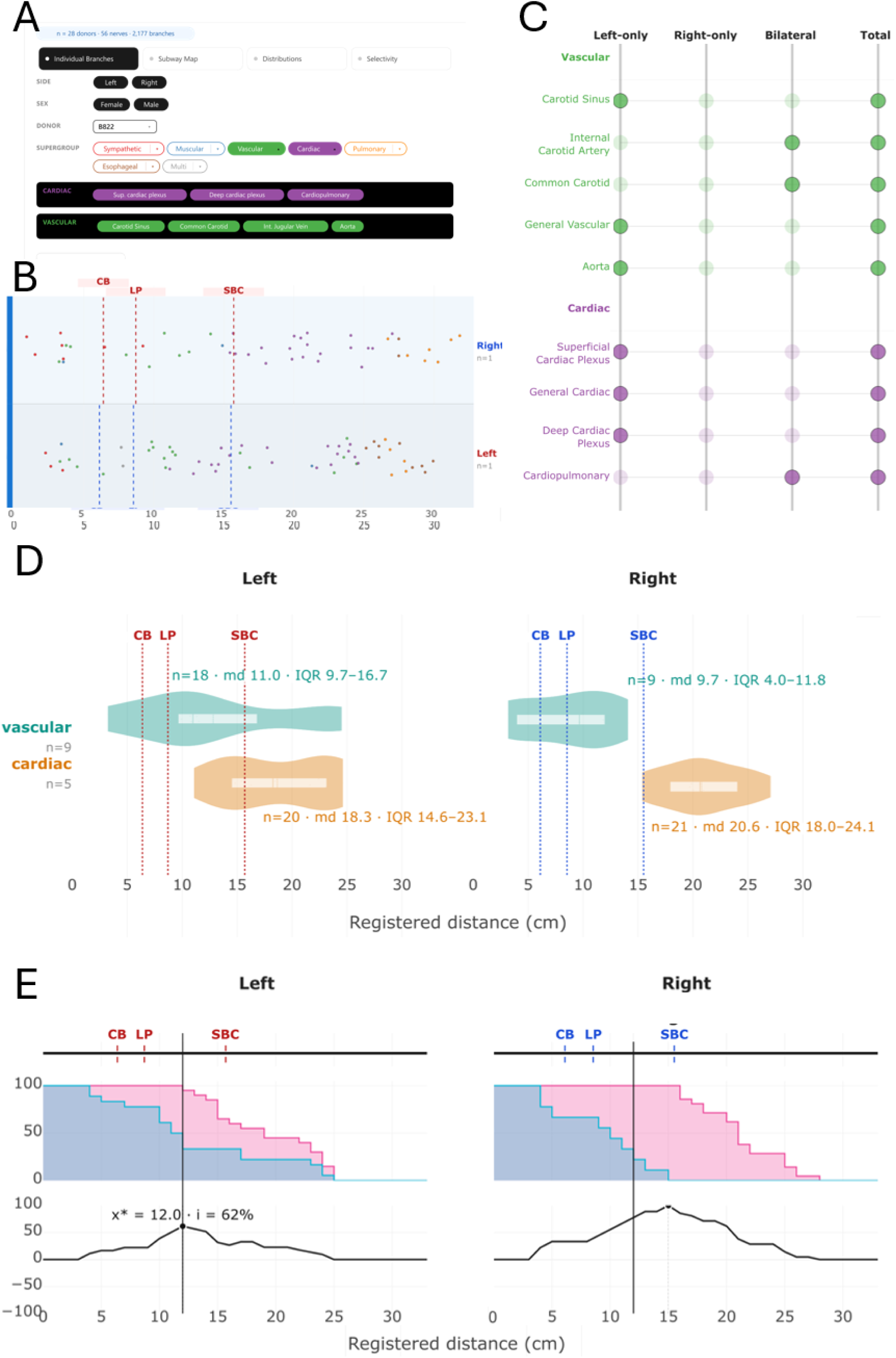
An interactive, online tool for exploring patterns of emergence of vagus nerve branches, in single or multiple donors. (A) Query-building interface. User can select side, sex, donor, branch group and subgroup to define branch populations for visualization or comparison. (B) Individual branch view. Branch emergence is shown along the landmark-registered axis for the selected nerves, with each branch represented by a point colored according to its group, allowing inspection of single-subject and side-specific anatomy; vertical lines mark the cohort-mean positions of the carotid bifurcation (CB), laryngeal prominence (LP), and superior border of the clavicle (SBC). Raw distances for each nerve is also available on explorer. (C) Branch prevalence view (“subway map”). Presence of groups or subgroups of branches in selected donors, shown as left-only, right-only or bilateral, and overall presence in the donor cohort. (D) Distance distribution view. Branch-emergence distances for selected branch groups, compared across left and right nerves, individual or pooled donors, sex-stratified groups, or custom-defined sets; annotations report number of branches (b), number of nerves that have that branch (n) median (md), and interquartile range (IQR). (E) Relative branch emergence view. Users define target and off-target branch sets and estimate, for a given level along the vagal trunk, the cumulative fraction of target and off-target branches emerging distally to that level (top) and an anatomical separation index index (bottom); x* marks the level of maximal index (i). The abbreviations listed below are used consistently in all subsequent figures: JF, jugular foramen; CB, carotid bifurcation; LP, laryngeal prominence; SBC, superior border of the clavicle; md, median; IQR, interquartile range; b, number of branches; n, number of nerves containing the branch; i, relative-emergence (“anatomical separation”) index; x*, level of maximal index.

### 2. General patterns in vagal branch emergence

We identified 2,177 branches (997 left, 1,180 right) across 56 nerves from 28 donors. Most emerge in the thoracic region below the clavicle (61.1%), followed by the upper cervical (18.1%), lower cervical (14.0%), and mid-cervical (6.8%) segments; branching density is highest in the thorax and lowest in the lower cervical segment (Table 1). The overall arrangement of the branch groups and their subgroups along the left and right nerves is summarized in Figure 2.

Branch groups emerge in a consistent proximal-to-distal order (Figure 3, Figure 4, Table 2): sympathetic branches are the most proximal (median 3.9 cm [3.0–6.6]), followed by muscular (6.2 [3.4–17.0]), vascular (9.6 [5.5–15.1]), and multiple-target branches (11.8 [4.4–18.3]), and then by cardiac (20.1 [16.9–22.5]), pulmonary (25.0 [22.8–27.6]), and esophageal branches (26.5 [24.3–29.1]) most distally. This order follows the positions of the organs each group innervates, from the neck down into the thorax. The superior border of the clavicle marks a structural transition between two embryological domains: pharyngeal-arch (cervical) targets cluster proximally and primitive-mediastinum (thoracic) targets distally. Linear discriminant analysis places the cervical–thoracic boundary at 17.2 cm (left) and 15.1 cm (right), near the clavicle, and classifies branches by domain with accuracy 0.85 (left)/0.91 (right), sensitivity 0.89/0.95, specificity 0.81/0.83, and area under the ROC curve 0.93/0.98 (Supplementary Table S12). The high AUC confirms the domains are well separated, while the specificity of 0.81–0.83 indicates that roughly 15–20% of branches fall on the opposite side of the boundary, so the clavicle is a genuine transition rather than a sharp anatomical line.

**Figure 3.**
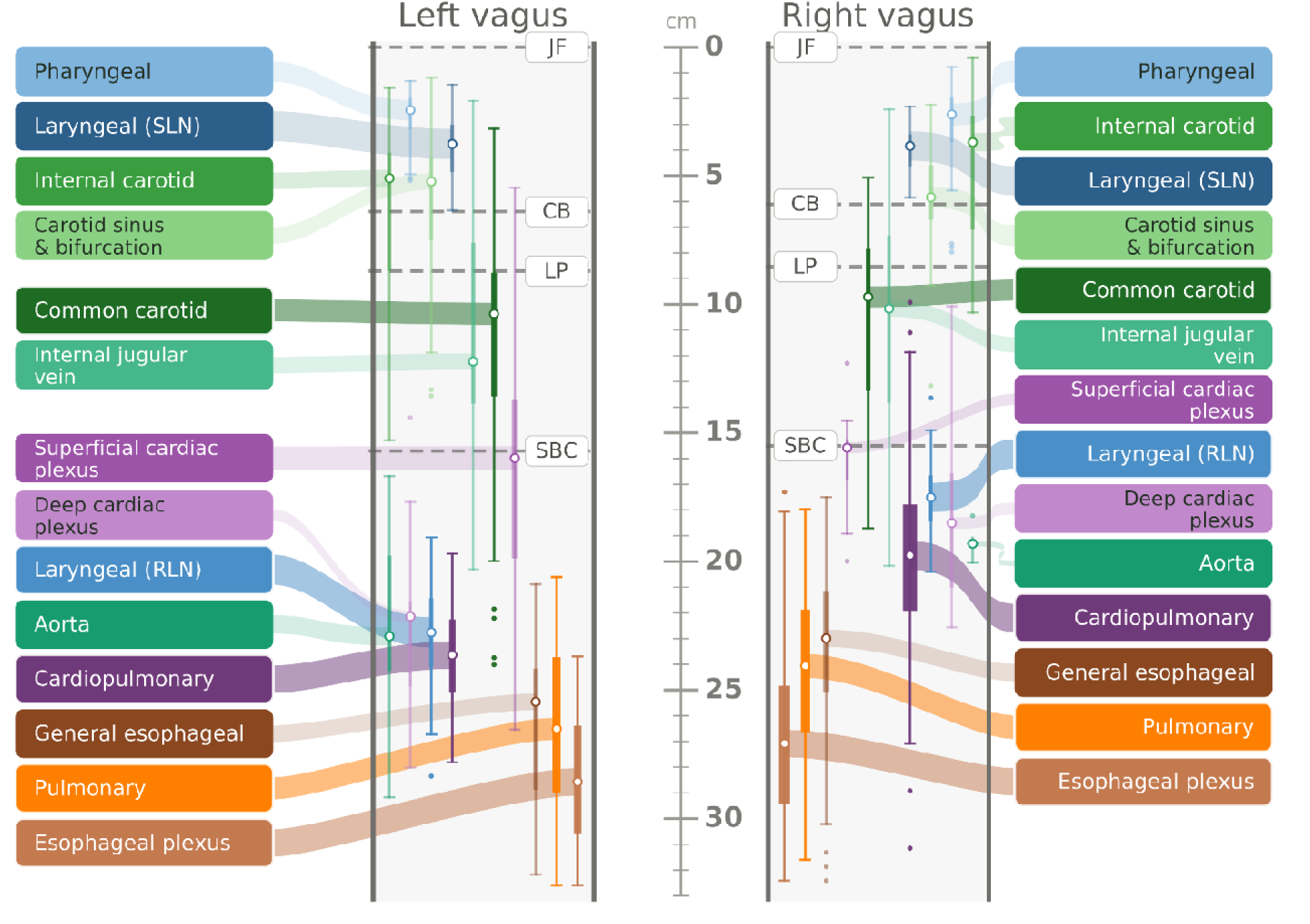
Summary schematic of the emergence of groups and subgroups of branches along the left and right vagus nerves. For each subgroup, the median distance of emergence from the jugular foramen (JF) and its interquartile range are shown along the vagal trunk, together with three anatomical landmarks (carotid bifurcation, CB; laryngeal prominence, LP; superior border of the clavicle, SBC). Each subgroup is plotted at its median registered distance (circle marker), with the interquartile range as a thick vertical bar and whiskers extending to the furthest branches within 1.5×IQR of the quartiles; branches beyond the whiskers are shown as dots. Values are pooled across donors and plotted separately for each side. The thickness of the interquartile bar is scaled to the number of branch events contributing to that subgroup, and the width of the curve-like projection (ribbon) linking each label to its bar is scaled to the subgroup’s prevalence (the number of donors in which it is present on that side). Horizontal dashed lines mark the cohort-mean positions of the three landmarks. Other abbreviations: SLN, superior laryngeal nerve; RLN, recurrent laryngeal nerve.

**Figure 4.**
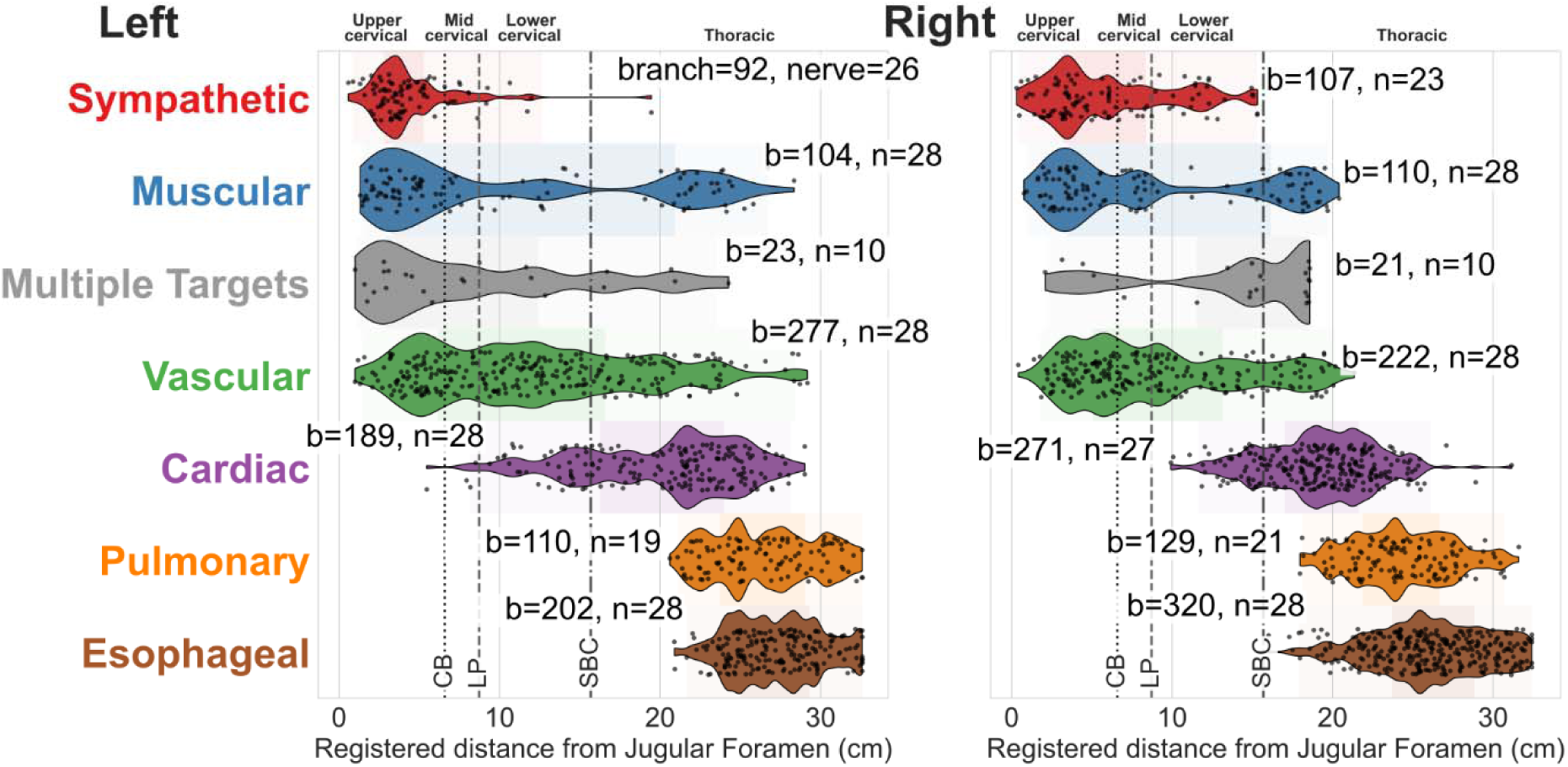
Emergence of vagal branches follows a proximodistal ordering according to the innervated organ or system. For each group of branches, the distribution of emergence distances is shown as a violin plot with a shaded spread band (darker central band, the interquartile range), for the left and right nerves; the upper cervical, mid cervical, lower cervical, and thoracic segments are labelled at the top, and labels report the number of branches (b) and nerves (n) containing the branches. Groups emerge in a proximal-to-distal order: sympathetic, muscular, multiple-targets, and vascular in the neck, then cardiac, pulmonary, and esophageal in the thorax. Sex- and side-stratified values are given in Supplementary Table 2.

**Table 2.**
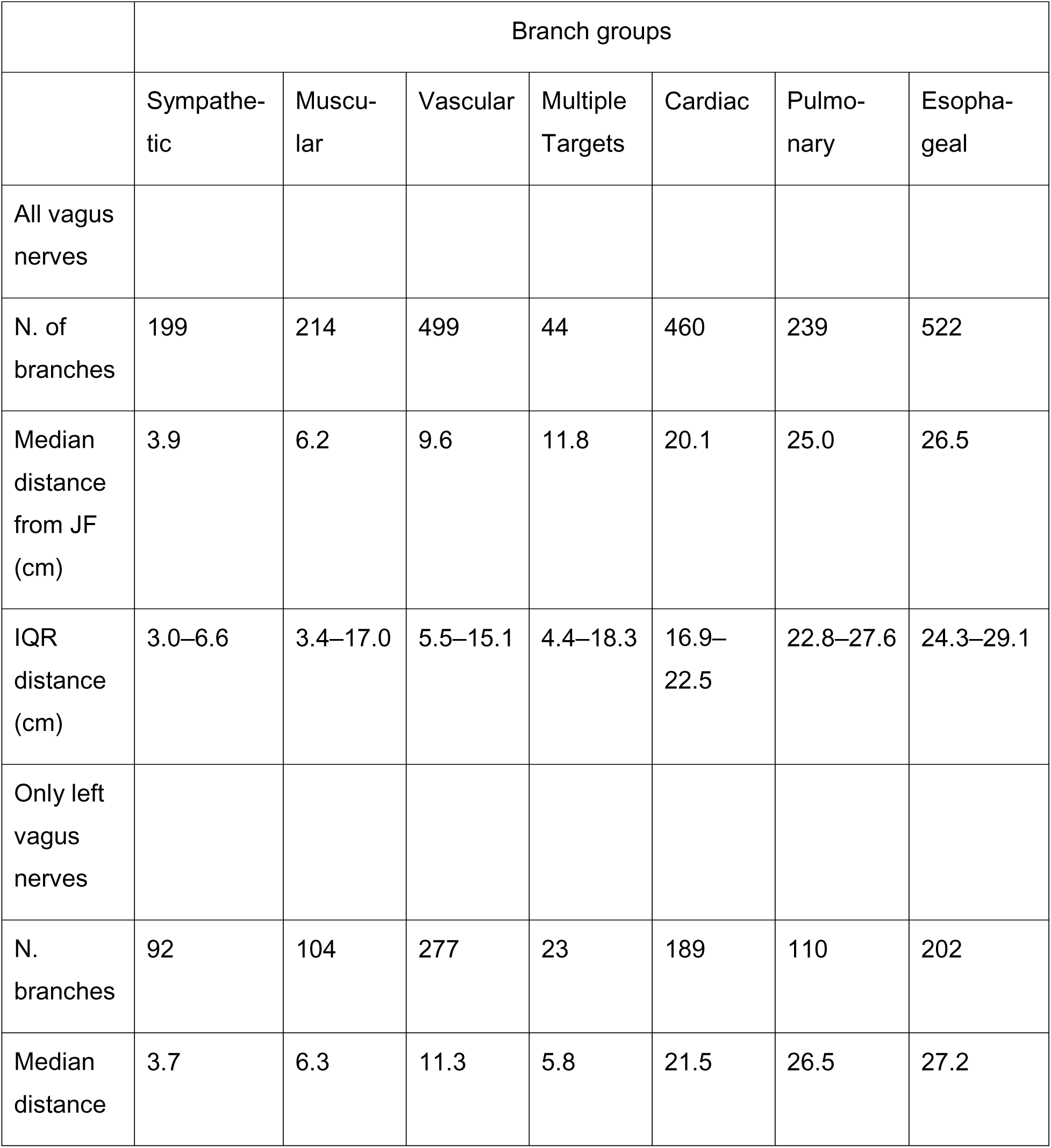
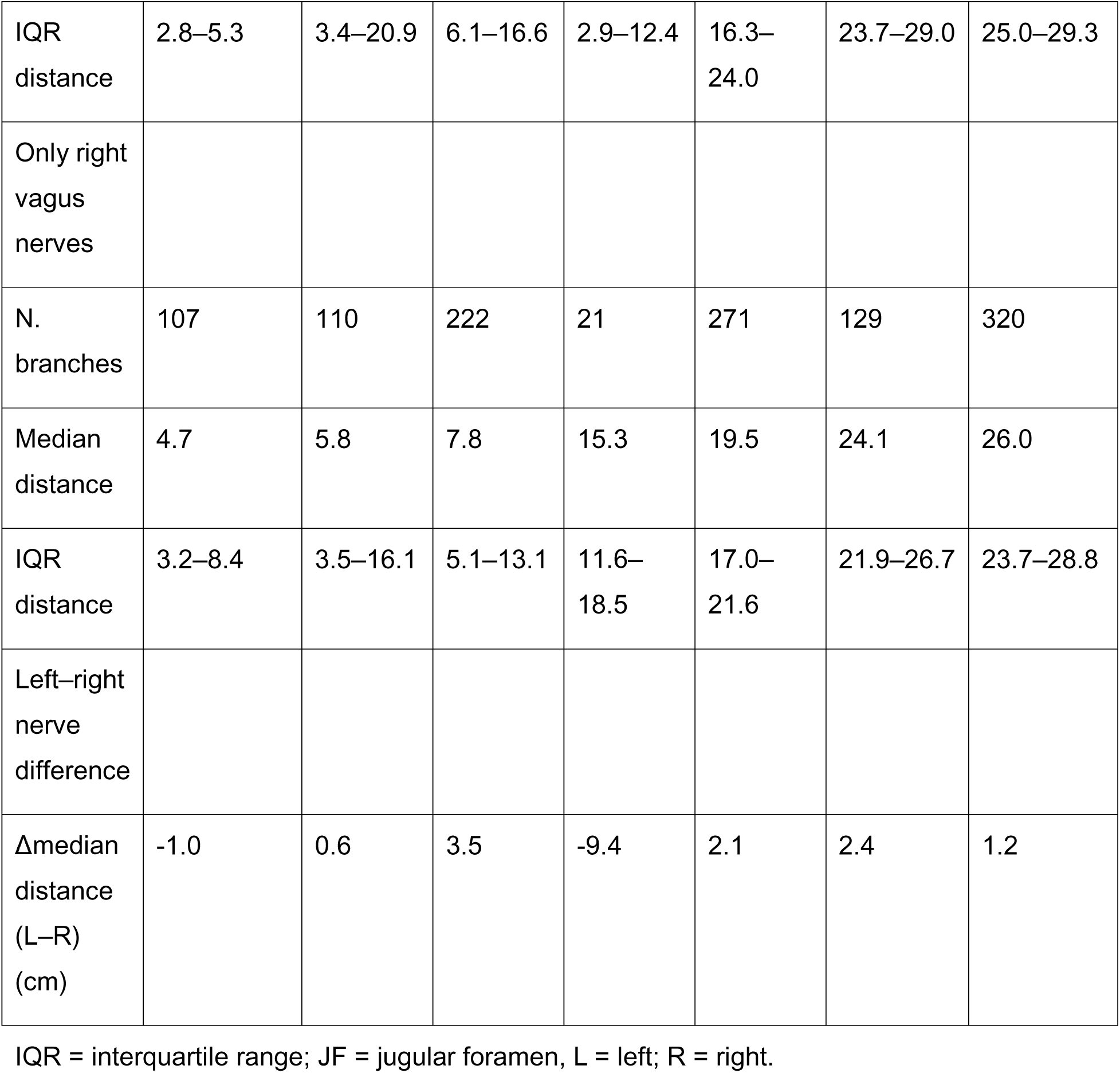
Emergence of different branch groups (distance from jugular foramen), for left and right vagus nerves. Values are shown for the total (pooled left and right) dataset and separately for the left (L) and right (R) vagus nerves as branch count (n), mean registered distance from the jugular foramen (cm), interquartile range (IQR), and side difference in mean emergence location (Δmean [L–R], cm). Positive Δmean values indicate more distal emergence on the left, whereas negative values indicate more distal emergence on the right.

Descriptive statistics for distances from JF of emergence of specific groups (median and interquartile range) are given in Table 2. Model-estimated positions are least-squares means [95% CI] and effect sizes are the least-squares-mean distance from each subgroup to the preceding (next-more-proximal) one, with the corresponding p-value, from per-group mixed-effects models with a donor random intercept are given in Table 3. Side and sex effects were assessed both by paired Wilcoxon signed-rank on per-donor medians and within the mixed-effects models (Sex-stratified data in Supplementary Figure 1, Supplementary Tables S2–S6 for non parametric tests; Supplementary tables S7–S11 for mixed effect model results). Non-parametric testing found no sex differences and side asymmetries only in the recurrent laryngeal, cardiopulmonary, pulmonary, and general esophageal subgroups (reported by group below). Mixed-effects models additionally showed that vascular and esophageal branches emerge more proximally in male donors (by 2.6 and 2.5 cm; p = 0.03 and 0.003) and on the right side (by 2.8 and 3.0 cm; p = 0.004 and < 0.001); the remaining significant terms were the subgroup-specific side asymmetries and within group ordering reported below.

**Table 3.**
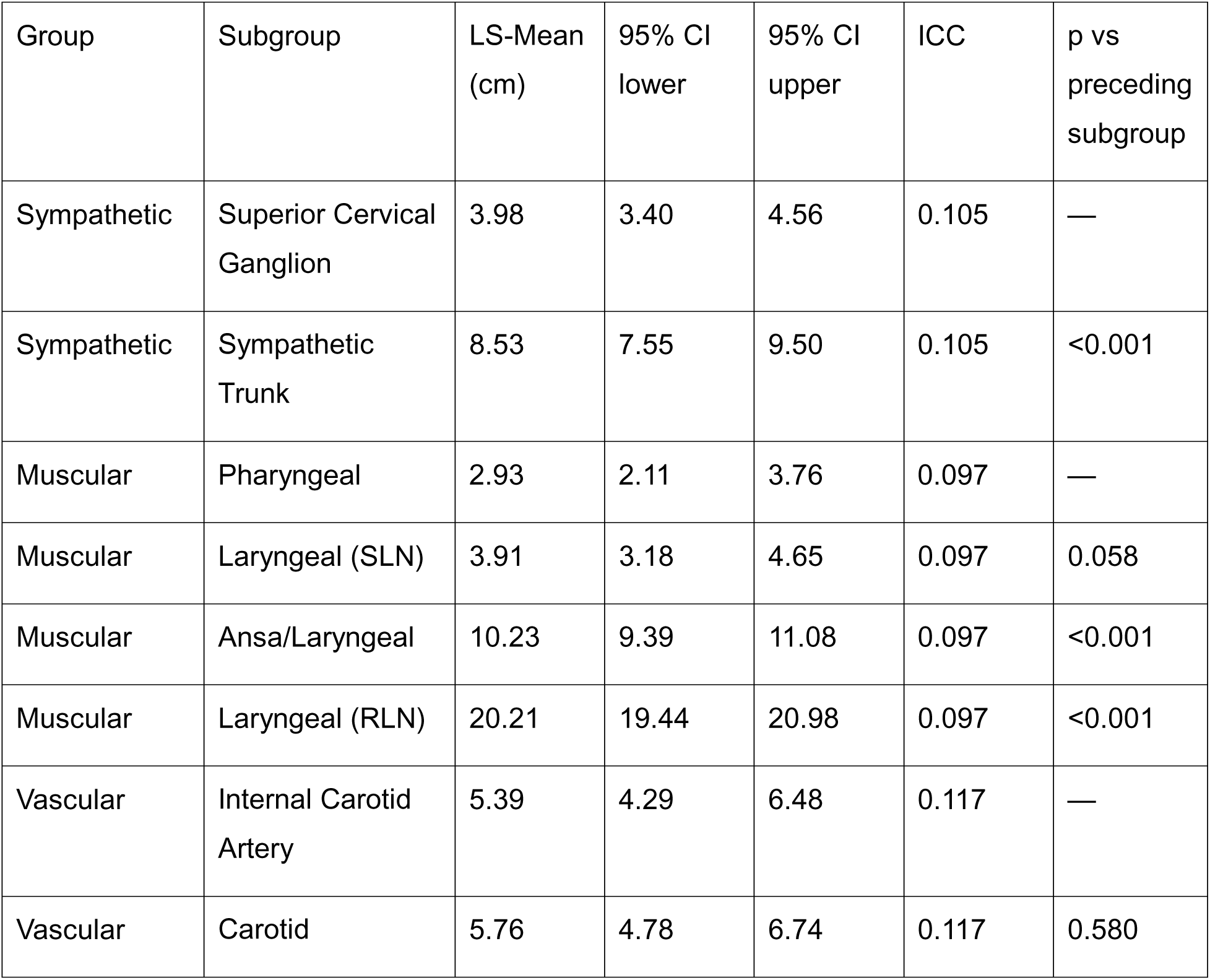
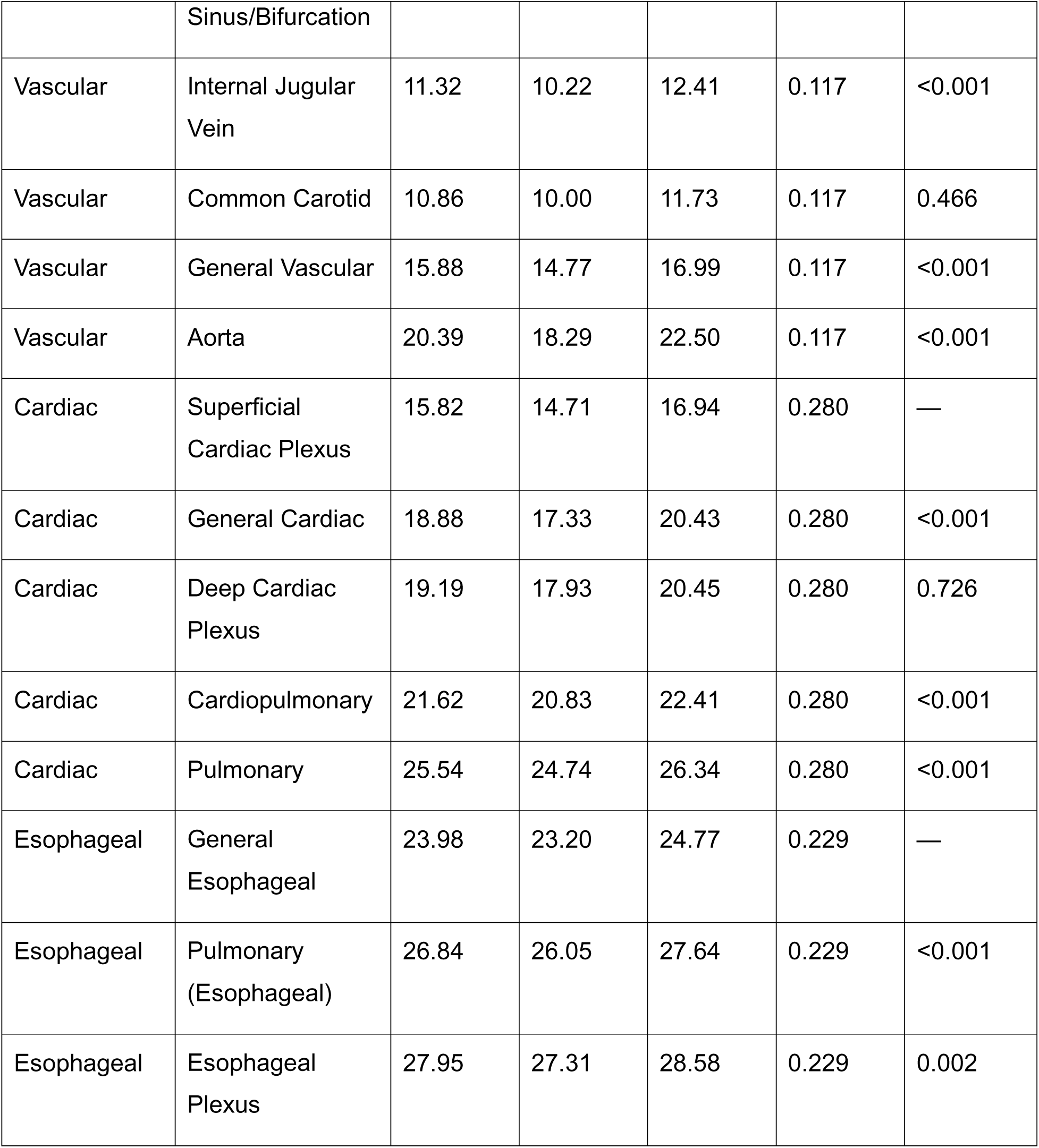
Model-estimated emergence position of each branch subgroup. The least-squares mean (and 95% confidence intervals, CI) represents the average emergence distance of a subgroup from the jugular foramen (cm), estimated from the respective mixed-effects model. The intraclass correlation (ICC) is the fraction of variance in branch emergence distance variation due to between-donor variability. The pulmonary group is included as a distinct subgroup within the cardiac group. The final column gives the raw p-value for the comparison of branch emergence distance between each subgroup versus the subgroup immediately proximal to it.

### 2. Emergence of specific groups of vagal branches

#### Sympathetic Group

The sympathetic group is the most proximal along the trunk and comprises two subgroups, the superior cervical ganglion and the sympathetic trunk; the superior cervical ganglion is one of the most prevalent subgroups overall, present in 96% of donors (Figure 5). The superior cervical ganglion emerges closest to the jugular foramen (median 3.6 cm [2.7–5.2]) and the sympathetic trunk more distally (9.1 [4.8–12.0]). In the mixed-effects model, which adjusts for side, sex, and the clustering of branches within a donor, the sympathetic trunk lies 4.6 cm distal to the ganglion as a least-squares-mean difference (p < 0.001; Table 3, Supplementary Table S7). Both subgroups sit within the cervical, pharyngeal-arch domain, above the clavicle, anchoring the proximal end of the whole-nerve order (Figure 6A).

**Figure 5.**
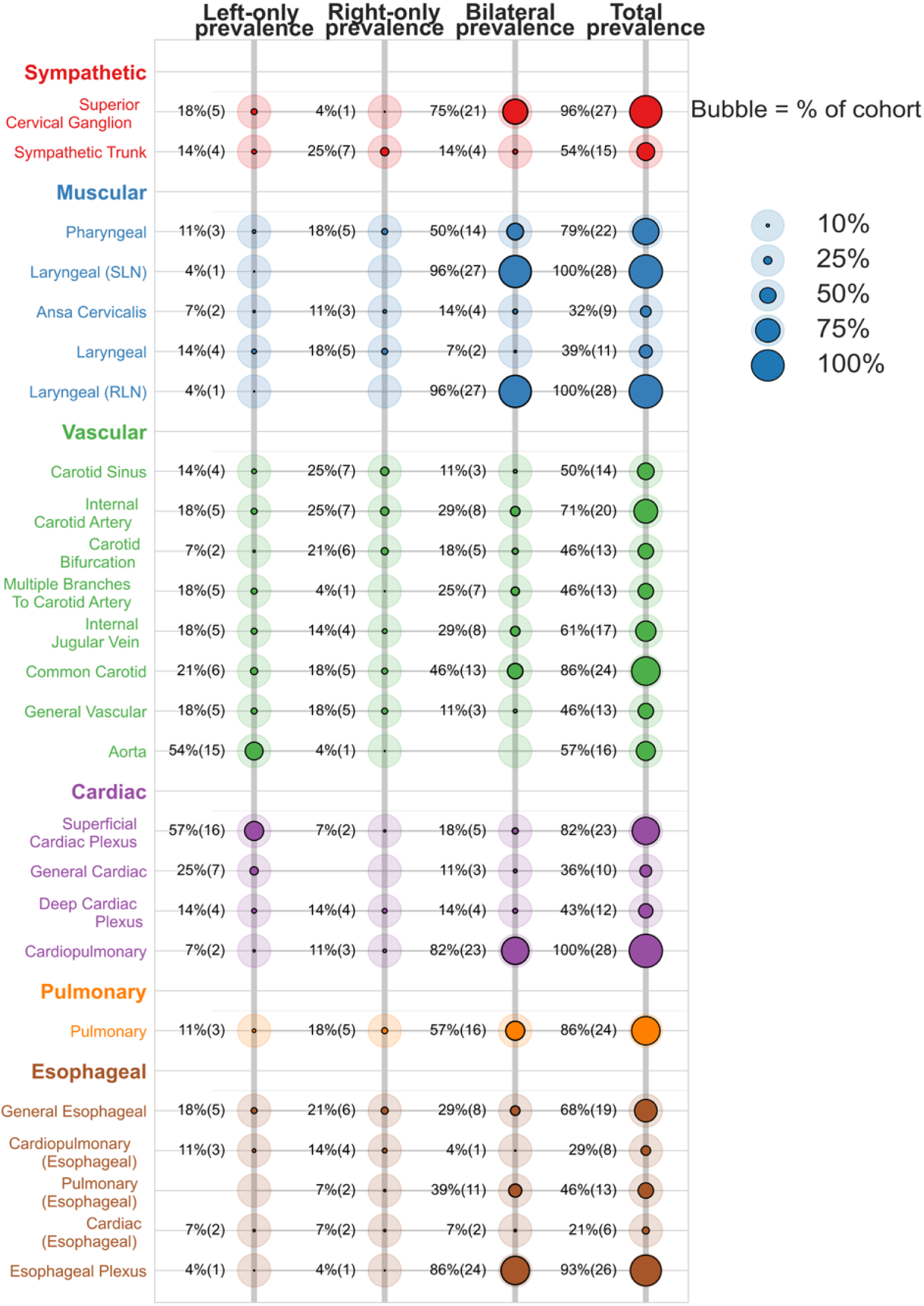
Prevalence and laterality of common vagal branch subgroups (“subway map”). Each row reports on one of the 25 most frequently observed branch subgroups, colored according to the group to which it belongs, and ordered proximal-to-distal within each group (mean positions in Table 3). The four columns provide prevalence of each branch subgroup in our cohort of 28 donors as left-only, right-only, bilateral, and total (either side). The faint, background bubble represents the size of a full cohort (100%), and the solid-colored bubble is scaled to the observed prevalence; labels give the percentage with the donor count in parentheses.

**Figure 6.**
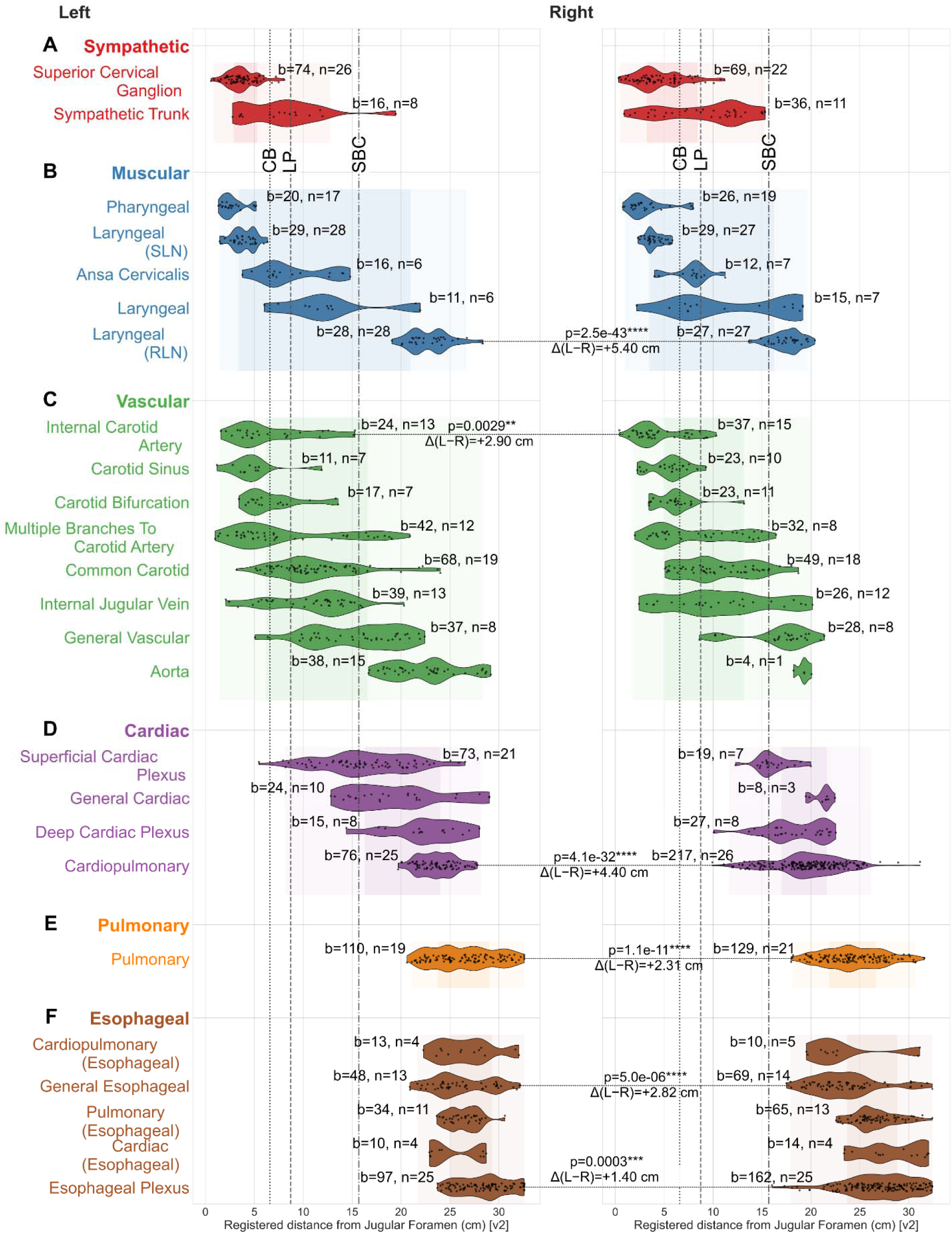
Emergence of subgroups of vagal branches follows a proximodistal order within larger groups. Distributions of branch-emergence distance for the most common subgroups within each group (violin plots; individual branches, landmark reference lines, background spread/interquartile bands, and b/s labels as in Figure 3), for the left and right nerves. For subgroups whose emergence level differs between sides, the left–right least square mean differences (Δ, L–R) (Suplementary Tables S7–11) and associated p-values are shown. Abbreviations are same as in Figures 2 and 3.

#### Muscular Group

The muscular group spans the widest proximodistal range of any group, its five subgroups reaching from the skull base to the upper thorax: pharyngeal, superior laryngeal (SLN), ansa cervicalis, general laryngeal, and recurrent laryngeal (RLN). The SLN and RLN are present in every donor and bilateral in 96%, and pharyngeal branches in 79% (Figure 5). Pharyngeal (median 2.6 cm [2.0–3.5]) and SLN branches (3.8 [3.3–4.8]) emerge together most proximally, ansa cervicalis and general laryngeal branches follow at the mid-cervical level (8.0 [6.9–9.2] and 11.9 [7.5–17.9]), and the recurrent laryngeal nerve emerges furthest of all, in the upper thorax (20.1 [17.7–22.8]). In the model, the SLN lies at essentially the same level as the pharyngeal subgroup (1.0 cm apart; p = 0.058), the ansa/laryngeal subgroup is 6.3 cm more distal (p < 0.001), and the RLN a further 10.0 cm distal again (p < 0.001; Table 3, Supplementary Table S8). Grouped by emergence level, these subgroups fall into three longitudinal zones — superior cervical, mid-to-low cervical, and upper thoracic (Figure 7B) — following the same significant ordering. The recurrent laryngeal nerve emerges markedly more distally on the left than the right (median Δ = +5.4 cm, p < 0.001)

**Figure 7.**
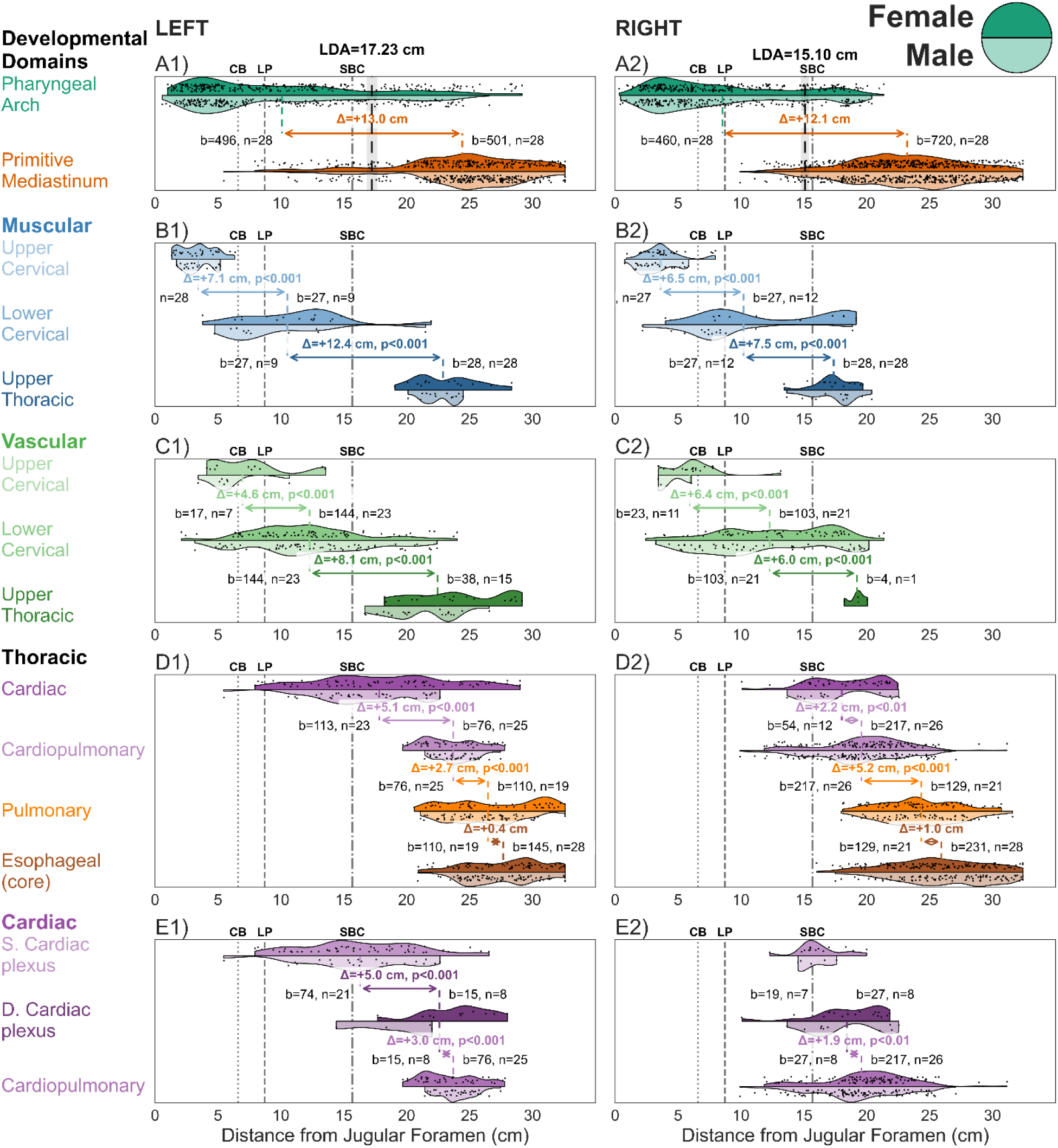
Groups and subgroups of vagal branches emerge within separate longitudinal zones. Each row represents a contrast between emergence zones of two groups or subgroups of vagal branches, with double-headed arrows providing the effect size (calculated as difference between least square means of branch emergence) between those groups or subgroups, computed separately for the left and right nerve (Supplementary Table S13). (A) The two developmental domains the pharyngeal-arch (sympathetic, muscular, vascular) and primitive-mediastinum (cardiac, pulmonary, esophageal) are separated by a level near the clavicle (shown as vertical thick dashed line), as determined by one-dimensional linear discriminant analysis. (B) Muscular branches emerge in three separate zones (upper cervical, lower cervical, upper thoracic). (C) Same for vascular branches. (D) Within the thorax, cardiac, cardiopulmonary, pulmonary, and esophageal branches emerge consecutively at separate levels. (E) Within the cardiac group, the superficial cardiac plexus emerges proximal to the deep cardiac plexus and cardiopulmonary branches. Abbreviations same as in Figure 2.

#### Vascular Group

The vascular group (n=499) is large and spans the neck and upper thorax, with subgroups for the carotid sinus, internal carotid, carotid bifurcation, common carotid (present in 86% of donors), internal jugular vein, general vascular, and aorta; the aorta is strongly left-biased (present on the left only in 54% of donors, the right only in 4%, and never bilaterally) (Figure 5). The three carotid subgroups emerge together most proximally (internal carotid median 4.3 cm [3.2–7.5], carotid sinus 5.3 [4.0–6.2], carotid bifurcation 5.9 [4.7–7.5]), the common carotid and internal jugular vein follow at the mid-cervical level (10.3 [8.3–13.4] and 11.8 [7.0–13.9]), and general vascular and aortic branches are the most distal (17.0 [12.4–19.1] and 21.9 [19.3– 23.7]). In the model, the internal carotid and carotid sinus/bifurcation sit together (0.4 cm apart; p = 0.58); the internal jugular vein is 5.6 cm more distal (p < 0.001) but level with the common carotid (p = 0.47); and general vascular and the aorta are 5.0 and 4.5 cm further distal than the preceding subgroup (p < 0.001; Table 3, Supplementary Table S9). As in the muscular group, these subgroups fall into superior cervical, mid-to-low cervical, and upper thoracic zones (Figure 7C).

#### Cardiac Group

The cardiac group **(n = 459)** comprises the superficial cardiac plexus (present in 82% of donors and left-biased), the deep cardiac plexus, general cardiac branches, and the cardiopulmonary subgroup, which is present in every donor and bilateral in 82% (Figure 5). Most cardiac branches emerge below the clavicle (81.0%); the surgically accessible portion above the clavicle (19.0%) lies almost entirely between the laryngeal prominence and the clavicle. The superficial cardiac plexus is the most proximal (median 15.9 cm [14.3–18.9]), followed by general cardiac (20.7 [16.8–21.7]) and deep cardiac plexus branches (20.8 [17.0–22.0]) at a similar level, with cardiopulmonary branches the most distal (21.0 [18.6–23.3]). In the model, general cardiac branches lie 3.1 cm distal to the superficial plexus (p < 0.001), the deep cardiac plexus is level with general cardiac (0.3 cm apart; p = 0.73), and cardiopulmonary branches are 2.4 cm distal again (p < 0.001; Table 3, Supplementary Table S10). This accessible component is asymmetric: on the left it is almost entirely superficial plexus (32 of 41 above-clavicle branches), whereas above-clavicle cardiopulmonary branching occurs only on the right (32 of 216 right branches versus none of 76 left). Cardiopulmonary branches also emerge markedly more distally on the left than the right (median Δ = +4.0 cm, p < 0.001). When pulmonary branches are added to this model, they fall furthest of all the cardiac subgroups, 3.9 cm beyond the cardiopulmonary branches (p < 0.001).

#### Pulmonary Group

Pulmonary branches are present in 86% of donors and emerge entirely below the clavicle (all 239), at a median of 25.0 cm [22.8–27.6], and more distally on the left than the right (median Δ = +1.7 cm, p = 0.016) (Figure 6E). Tested within the cardiac model, they lie furthest of all the cardiac subgroups, 3.9 cm beyond the cardiopulmonary branches (p < 0.001; Table 3, Supplementary Table S10).

#### Esophageal Group

The esophageal group is the most distal, and its esophageal plexus is present in 93% of donors and bilateral in 86% (Figure 5). All esophageal branches emerge below the clavicle. General esophageal branches are the most proximal of the group (median 24.2 cm [21.9–26.5]), the pulmonary-esophageal subgroup follows more distally (26.3 [25.3–27.7]), and the esophageal plexus is furthest of all (27.9 [25.4–29.7]). In the model, pulmonary-esophageal branches lie 2.9 cm distal to general esophageal (p < 0.001) and the esophageal plexus a further 1.1 cm distal (p = 0.002; Table 3, Supplementary table S11). General esophageal branches emerge more distally on the left than the right (median Δ = +3.2 cm, p = 0.047).

### 4. Branch-free intervals around anatomical landmarks

To estimate how much uninterrupted vagal trunk is typically available around surgically relevant anatomical landmarks, we measured, for each donor and side, the branch-free window straddling each landmark — the distance from the nearest branch proximal to the nearest branch distal — at the carotid bifurcation, laryngeal prominence, and superior border of the clavicle (Figure 8A; Supplementary Table S6). Window lengths are reported as the median [interquartile range] across nerves.

**Figure 8.**
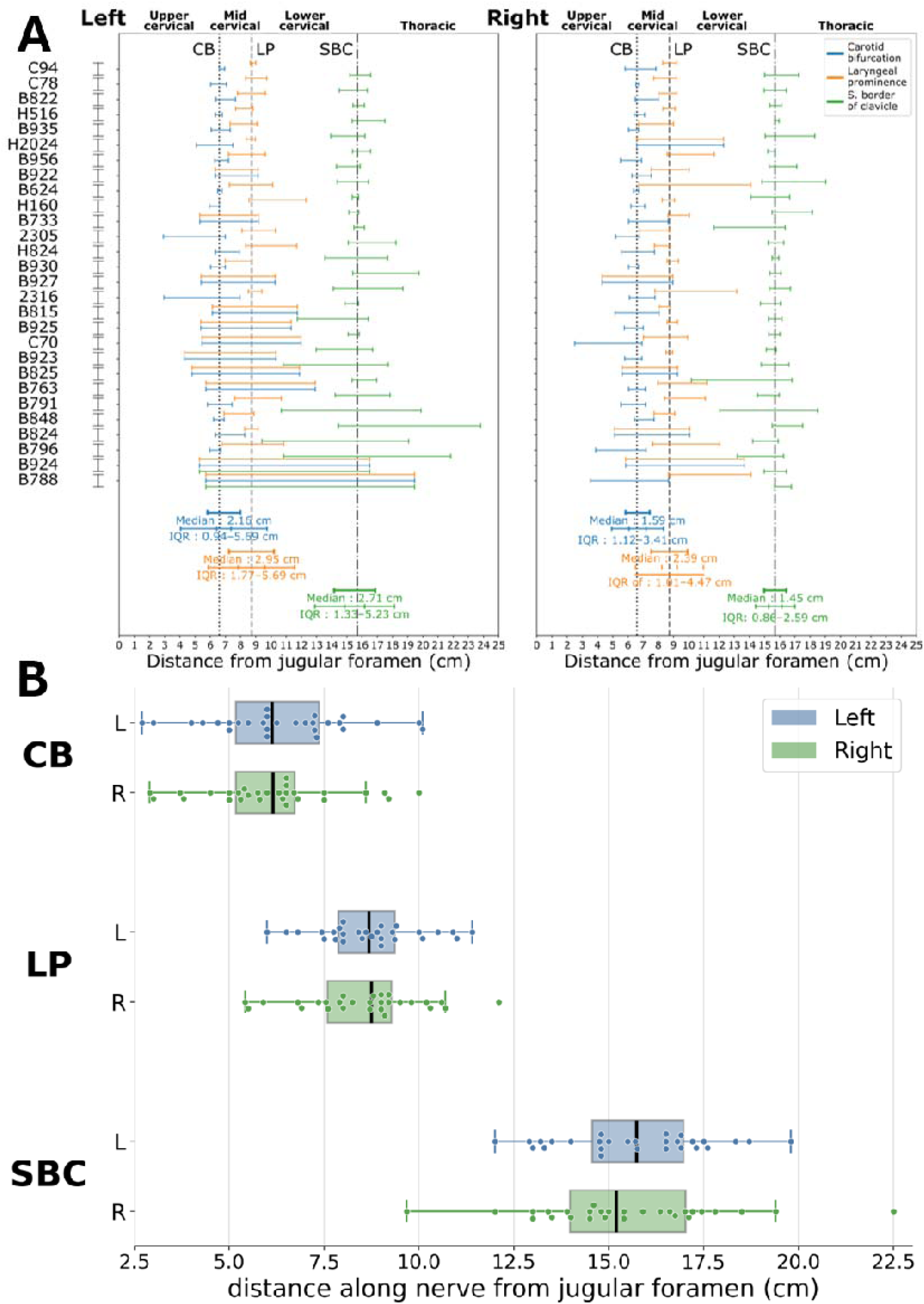
Branch-free intervals along the left and right vagal trunk. (A) Landmark-centered branch-free intervals for each donor and side, with cohort medians and IQRs shown at the bottom. (B) Between-donor spread of landmark positions, left versus right, demonstrating level of inter-individual variability. Abbreviations same as in Figure 2.

Branch-free windows are consistently larger on the left than the right. On the left, the widest windows are around the laryngeal prominence (median 3.0 cm [IQR 1.8–5.7]) and the superior border of the clavicle (2.7 cm [1.3–5.2]), with the carotid bifurcation slightly smaller (2.2 cm [0.9– 5.7]); the corresponding right-side windows are shorter (2.4 [1.0–4.5], 1.5 [0.9–2.6], and 1.6 [1.1–3.4] cm, respectively). Across donors these windows vary widely: the shortest are near zero — a branch emerging essentially at the landmark — while the longest uninterrupted intervals extend to ∼13.8 cm on the left and ∼7.8 cm on the right (Figure 8A, Supplementary Table S6). This wide donor-to-donor spread means the usable window at a given landmark is patient-specific rather than fixed.

Landmark locations themselves also vary considerably between donors (Figure 8B). The carotid bifurcation and laryngeal prominence sit at relatively consistent levels (median 6.1 cm [IQR 5.2– 7.3] and 8.7 cm [7.8–9.4]), whereas the superior border of the clavicle is more variable (15.5 cm [14.0–17.0]); full ranges span 2.7–10.1, 5.4–12.1, and 9.7–22.5 cm, respectively (Table 1). Left and right landmark-position distributions overlap substantially.

Landmark locations vary considerably between donors (Figure 8B). The carotid bifurcation and laryngeal prominence lie with relatively narrow ranges (6.25 cm, range 2.70–10.10; and 8.62 cm, range 5.41–12.10), whereas the superior border of the clavicle spans a wider range (15.60 cm, range 9.68–22.50; Table 1). Left and right side landmark location distributions overlap significantly.

### 5. Anatomical separation of branches mediating target vs. off-target effects

The level at which a device is implanted constrains which branches lie within reach of stimulation: branches emerging distally to that level are anatomically positioned to be engaged, whereas those emerging proximally will be spared. It is feasible that placement of a stimulation device at a given level of the vagal trunk is expected to engage a mix of targeted and off-target branches. Whether this translates into functional selectivity was not tested here. We sought to determine whether the relative emergence of vagal branches mediating some targeted and off-target responses anatomically supports such an approach in the human vagus nerve.

The separation between broad thorax–neck branch emergence peaks distally to the superior border of the clavicle : at that level, the probability of distal emergence of targeted (thorax) vs. off-target branches (neck) is maximal. When primitive mediastinum domain (cardiac, pulmonary, esophageal) branches are targeted against pharyngeal arc branches (cardiac, pulmonary, esophageal) their separation peaks at 17 cm on the left and 19 cm on the right, with high values of peak anatomical separation (mean i(x*) = 73% left; 79% right) (Figure 9A). This indicates a strong dual-domain organization in which thoracic targets become reliably enriched distal to a clavicle-adjacent window.

**Figure 9.**
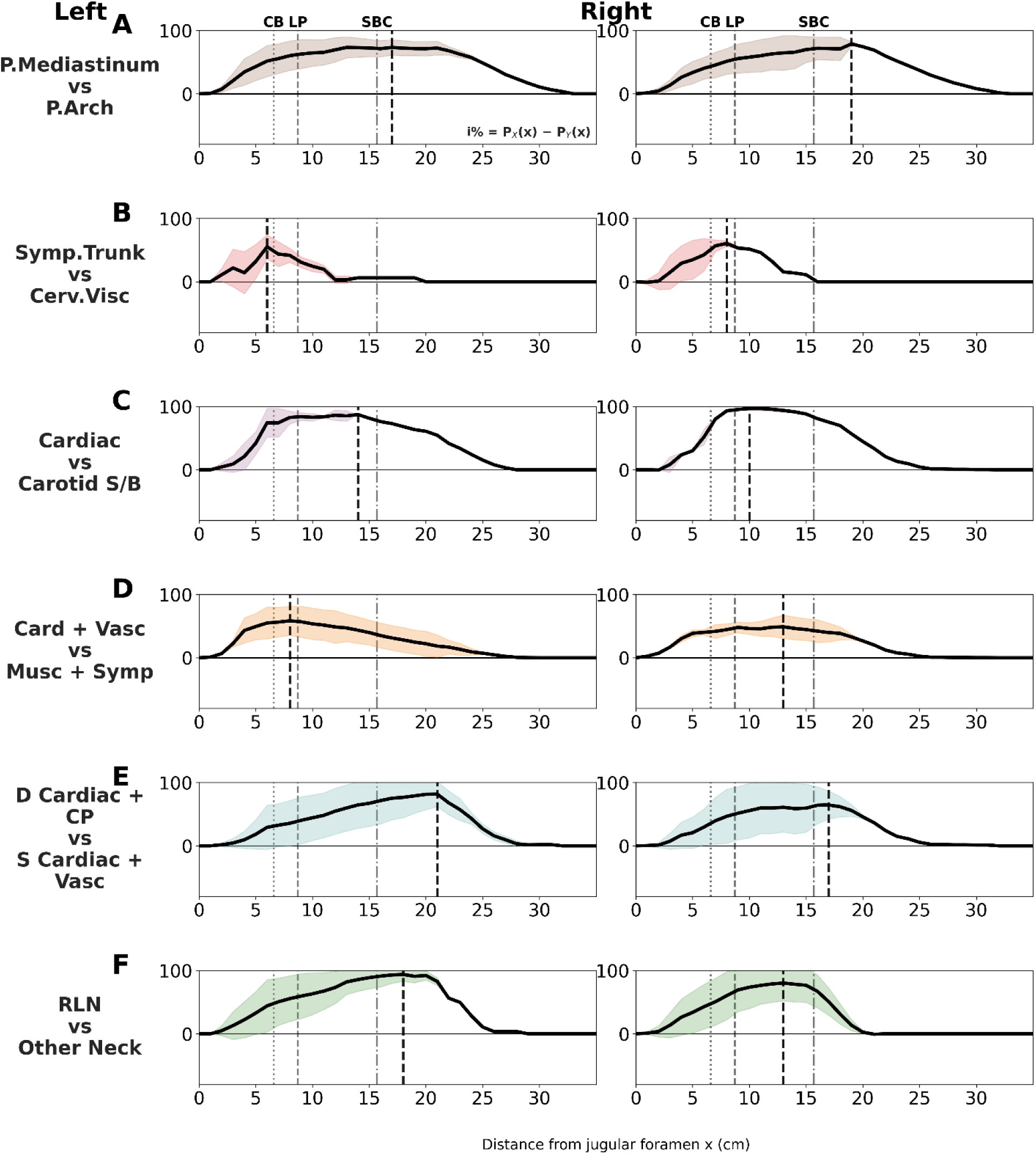
Maximal anatomical separation for different contrasts between “target” branches and “off-target” branches. For each contrast, the anatomical separation index i(x) is calculated as the probability of target minus probability of off-target branches having emerged rostrally relative to a level at distance x from the JF. The black dashed line marks x*, where the index is maximal and the target vs. off-target branch separation is greatest; in principle that is the level where engagement of the target branch would spare the most off-target branches (black trace is mean index; shared area is SD of index).(A) Primitive mediastinum (cardiac, pulmonary, esophageal) vs pharyngeal-arch derivatives (sympathetic, muscular, vascular neck branches). (B) Sympathetic trunk vs upper cervical visceral branches (pharyngeal, superior laryngeal [SLN], carotid sinus, carotid bifurcation) (C) Cardiac vs carotid sinus and carotid bifurcation. (D) Cardiac + vascular vs muscular + sympathetic. (E) Deep cardiac + cardiopulmonary vs superficial cardiac + vascular. Reference lines and abbreviations same as in Figure 2.

When cardiac branches are targeted, maximum separation from other branches shifts proximally into a narrower region around 13–15 cm from the jugular foramen (Figure 9B-D). Specifically, for cardiac against neck/muscular branches, separation peaks at 13 cm on the left and 15 cm on the right (Figure 9B). Targeting cardiac vs. vascular branches, separation peaks at 14 cm on the left and 13 cm on the right (Figure 9C). Targeting branches from the superficial cardiac plexus against neck and vascular subgroups, anatomical separation peaks at 13 cm on the left and 14 cm on the right (Figure 9D). Finally, targeting branches from the deep cardiac plexus against other cardiac or pulmonary branches, anatomical separation peaks at 21 cm on the left and 17 cm on the right, with high peak separation (Figure 9E).

Taken together, these results suggest that a window extending from 13 to 15 cm from the jugular foramen maximizes separation between targeted, cardiac and superficial cardiac branches relative to neck and muscular off-target branches, whereas deep cardiac targets have a more distal maximal separation from other cardiac or pulmonary off-target branches, approximately 17–21 cm from the jugular foramen.

## Discussion

### The need for a standardized map of vagal branch emergence

Most vagus nerve anatomy studies describe branching relative to local landmarks or within single surgical compartments. For example, studies on the upper cervical vagus include the jugular foramen and skull base (Tummala et al. 2005; Liang et al. 2019). Studies on mid-cervical vagus focus on anatomical relationships of the recurrent laryngeal nerve within the thyroid field (Cakir BO et al. 2006; A. Patra 2023)] . Reports on thoracic and abdominal regions study vagal branches specific to cardiac (Kawashima 2005), pulmonary, mediastinal (Weijs et al. 2015), or gastric (Parpex et al. 2020) targets. Each body of work is informative within its own compartment, although the absence of a shared longitudinal coordinate frame makes it difficult to compare branch positions across compartments, across individuals, or between studies. Our online tool and standardized map, registered to three surgically identifiable landmarks and spanning from the jugular foramen through the distal thorax, addresses this gap. By normalizing branch emergence distances to a common scale, our framework makes it possible to contextualize older compartment-specific observations onto a common coordinate system and to quantify inter-subject variability in a way that isolated regional descriptions cannot. The registration framework is compatible with emerging 3D thoracic reconstruction efforts (Wu et al. 2013) and could enrich computational atlases with probabilistic branch-emergence data as branch-level ground truth scales.

The variability in landmark locations across donors is one of the motivations for our registration approach. A branch emerging 15 cm from the JF could lie above the clavicle in one donor and well below it in another. Registering each nerve to these common, surgically identifiable landmarks removes some of this variability: normalized locations rather than absolute distances make levels of emergence comparable across donors and, by extension, potentially more practical to surgical practice.

The present study and the associated exploratory tool are intended to be an anatomical reference that different users may use in different ways. Surgeons may use our resource to plan the placement of a stimulation device at a level on the vagal trunk compatible with the desired combination of target and off-target effects, using the relative anatomical separation data (Figure 6, 7<u>, 9</u>), while considering surgical and anatomical limitations, using branch-free interval data (Figure 8A) and measurements from accessible anatomical landmarks (Figure 8B). Device developers and engineers may use our resource in decisions on device design vs. intended use, e.g. design of an electrode based on desired level of placement, or device placement that minimizes lead length among placements that target similar branches. To anatomists, our resource may offer a standardized, landmark-registered frame on which our findings and findings from other studies using different anatomical references can be compared (Fig 2). Importantly, given the documented anatomical variability of vagal branching, our resource is meant to inform and complement, but not replace, patient-specific confirmation of branch anatomy.

### Proximal-distal organization of vagal branch emergence

In our cohort of donors and nerves, vagal branches follow a relatively consistent proximal-to-distal sequence (Figure 3, 6, 7, Table 2), consistent with prior literature. For example, reports on branches emerging close to the jugular foramen (Tummala et al. 2005; Linn et al. 2009; Song et al. 2008; Liang et al. 2019), SLN and RLN branches within the cervical region (Cakir BO et al. 2006; A. Patra 2023), sympathetic trunk communications in the mid and lower neck (Sato et al. 1997; Seki et al. 2014) as well as sympathetic-marker-positive fibers within the vagus at cervical and thoracic levels (Kawagishi et al. 2008; Seki et al. 2014), all agree with our findings. In the thorax, cardiac autonomic branches (Kawashima 2005), pulmonary branch dissections, mediastinal anatomy (Weijs et al. 2015), and esophagogastric plexus descriptions interconnected (Osugi et al. 2017; Haenssgen et al. 2020; Sisu et al. 2012; Weijs et al. 2017) each describe region-specific patterns that fit the proximal-distal order we describe here.

We calculated that the boundary between pharyngeal-arch-domain branches (sympathetic, muscular and vascular) and primitive-mediastinum-domain branches (cardiac, pulmonary, esophageal) lies at 17.21 cm on the left and 15.14 cm on the right, close to the mean clavicular position (15.60 cm). Most muscular and vascular branches emerge within three distinct zones: a superior cervical group (pharyngeal, SLN, carotid sinus, internal carotid), a mid-to-lower cervical group (ansa cervicalis, general laryngeal, common carotid, internal jugular vein), and an upper thoracic group (RLN, aortic branches) (Figure 7B, C). These three zones parallel three pharyngeal arches: SLN is associated with fourth-arch derivatives and RLN with sixth-arch derivatives; the intermediate muscular branches (ansa cervicalis, general laryngeal), present in approximately one third of our cohort, may represent a transitional region related to the rudimentary fifth pharyngeal arch (Frisdal and Trainor 2014).

### Separation of cardiac, cardiopulmonary, pulmonary, and esophageal branches

Within the thorax, cardiac, cardiopulmonary, and pulmonary branches emerge at successively more distal, statistically significant levels (Figure 7D): cardiac branches emerge most proximally, cardiopulmonary branches at intermediate levels, and pulmonary branches in the mid-to-distal thorax, entirely below the clavicle. Esophageal branches form the most distal group, but their emergence overlaps substantially with pulmonary branches rather than separating from them, and they too are entirely thoracic.

We observed the expected left-right asymmetry of the cardiac plexuses: the deep cardiac plexus emerged more distally on the left than on the right, so that it separated from the superficial plexus on the left but not the right (Figure 7E). Both on the left and the right emergence of pulmonary and esophageal branches are overlapping, consistent with postmortem studies describing a shared peri-esophageal compartment that interfaces with pulmonary plexus structures (Weijs et al. 2015, 2017). These findings support treating cardiopulmonary branching as a separate category rather than merging it with the cardiac or pulmonary groups. They also confirm that cardiac branches are more dispersed along the thoracic course than pulmonary branches, which cluster more tightly. This dispersal is consistent with reports showing that vagal contributions to the cardiac plexus are anatomically interconnected (González et al. 2023), whereas pulmonary branches follow a more concentrated emergence pattern aligned with the posterior pulmonary plexus (Weijs et al. 2015).

Most prior studies on vagal anatomy use individually chosen surgical or radiologic landmarks (aortic arch, carina, lung hilum, diaphragmatic hiatus). Our registered map helps translate these observations into a common anatomical axis. Reports treating the carina and lung root as key landmarks (Osugi et al. 2017; Wang et al. 2012) map to the mid-to-lower thoracic portions in our nerve template where cardiac, cardiopulmonary, and pulmonary branches separate (Figure 3, 6, 7). Anatomical observations on pulmonary branches reported in lower mediastinal dissections (Weijs et al. 2015) describe what branches look like once dissection intersects the pulmonary emergence band; our data further clarify where these branches cluster and how that clustering differs between sides (Figure 6, Figure 7D, Supplementary Table S2).

### Implications for surgical placement of vagus nerve stimulation devices

We found that branch-free intervals around landmarks differ by side: on the left, the longest intervals are above the clavicle and below the carotid bifurcation (Figure 8, Supplementary Table S6); on the right, the same intervals are shorter (Figure 8A, Supplementary Table S6). Intervals around the laryngeal prominence and carotid bifurcation are short on both sides, though variability across individuals is large. These findings suggest that the and branch-free intervals around left clavicle and laryngeal prominence may offer more room for cuff placement with reduced risk of physical interference with an adjacent branch, with implications for branch preservation during cervicothoracic surgery, and operative planning when the vagus nerve is either the target or a structure at risk (Weijs et al. 2015; Ruurda J.P. TJ et al. 2016; Osugi et al. 2017).

The branch-free interval around a given landmark varies substantially between donors (Figure 8B), ranging from a branch lying essentially at the landmark to an uninterrupted interval of several centimeters. For device placement this means the usable window at a landmark is patient-specific rather than fixed: a level offering a comfortable branch-free margin in one patient may lie immediately adjacent to a branch in another. Population-level margins therefore indicate where a branch-free window is typically found and how large it tends to be, but cannot guarantee the margin in any individual. This reinforces the value of confirming the branch-free interval in the specific patient before implantation, for example with high-resolution ultrasound, so the cuff can be positioned within that individual’s available window rather than an average one.

It is important to distinguish anatomical opportunity from surgical accessibility. The more proximal windows identified here lie in the cervical and supraclavicular regions, which are directly accessible and some already used for vagus nerve stimulator placement. The more distal windows lie in the superior mediastinum and upper thorax, and placement of a device below the clavicle would require thoracic surgery, which is more invasive and possibly less practical for the sole goal of VNS selectivity. Nonetheless, data on these distal windows are informative in three ways: they define the anatomical limits of what cervical placement can achieve; they may be relevant when a device is placed during esophageal or cardiac surgeries for other reasons; and they provide anatomical data for computational models of neuromodulation and for future less or non-invasive approaches.

### Implications for more selective vagal neuromodulation

Despite its implications for selective neuromodulation, our mapping study does not report functional, nor neuromodulation selectivity. The extent to which a stimulation device placed at a certain level of the vagal trunk engages fibers and functions from an organ while sparing others, depends on additional factors that were not assessed here. Such factors include electrode geometry and spatial relationship to underlying nerve anatomy (Schiefer et al. 2010; McGee and Grill 2014; Fitchett et al. 2021; Strauss et al. 2023), electrode-tissue interface, including possible encapsulation and tissue impedance (Umair Ahmed et al. 2021; Mughrabi et al. 2021) and stimulus waveform (Cheng et al. 2025; Chang et al. 2022). Therefore, anatomical separation of branch emergence is only one of the conditions that could facilitate and constrain neuromodulation selectivity.

Because each branch is formed by one or more fascicles emerging together from the trunk, branch emergence zones mapped here may correspond to regions in the trunk in which fascicles re-organize internally (Settell et al. 2020; Jayaprakash N et al. 2023; Thompson N et al. 2026). For example, in swine nerves, fascicles that arise from the nodose ganglion, and contain many afferent fibers, and fascicles that bypass the nodose ganglion, and contain many efferent fibers, are initially separate close to the nodose ganglion, but progressively merge into mixed fascicles in the caudal direction (Jayaprakash N et al. 2023). Conversely, fascicles that give rise to organ-specific branches remain separate close to branch emergence, but progressively merge with other fascicles in the rostral direction (Jayaprakash N et al. 2023; Nicole Thompson et al. 2025) . On the other hand, fascicles that give rise to some cardiac branches remain separate for much longer distances, in both swine (Jayaprakash N et al. 2023; Nicole Thompson et al. 2025) and in humans (Kronsteiner B et al. 2024).

Gross anatomy of branch emergence and microscopic anatomy of fascicles in the vagal trunk represent aspects of underlying nerve fiber organization: a motor branch innervating laryngeal muscles will likely be formed by separation of laryngeal fibers from one or more motor fascicles to form new fascicles that eventually lead to the emergence of the laryngeal motor branch. In that sense, the branch emergence zones identified in this study may correspond to regions where fascicular organization of the vagal trunk is expected to change; that intraneural fiber organization is likely different for different organ innervations. Importantly, our study does not resolve the functionally and neurochemically distinct fiber types within each branch (Kronsteiner B et al. 2024), which ultimately mediate the responses resulting from activation of the respective branches and likely have a significant impact on attainable selectivity of stimulation at different levels of the vagal trunk.

The analysis of relative emergence of targeted vs. off-target branches suggests possible sites for implantation of devices for more selective vagal neuromodulation (Figure 9). First, because sympathetic branches emerge within a few cm from the jugular foramen (Figure 3, 6), implanting a cuff above that level would be consistent with recruiting vagal–sympathetic communicating branches, but would also likely recruit muscular, vascular, cardiac, pulmonary, and esophageal branches that emerge distal to this level. However, on the right, a cuff placed near the laryngeal prominence may still engage sympathetic trunk (subgroup) branches while sparing more cranial pharyngeal, superior laryngeal, carotid sinus, and carotid bifurcation branches (Figure 9B).

Second, a device placed just cranial to the superior border of the clavicle, within a range of 10– 15 cm from the JF, would recruit cardiac-related branches on both sides, and would spare most carotid sinus and carotid bifurcation branches emerging distally to it (Figure 9C). Sinoatrial and atrial automaticity is primarily controlled by the superficial cardiac plexus, arising mostly on the left, whereas atrioventricular conduction is primarily controlled by the deep cardiac plexus, arising mostly on the right (Zandstra et al. 2021; Coote 2013). Placement of a device more distally on the left (21 cm) and more proximal on the right (17 cm) would be anatomically favoring the respective functions, assuming that functional engagement is consistent with branch emergence. (Figure 9E).

Third, a device placed proximal to carotid bifurcation (Figure 7C; Figure 9C) would favor recruitment of carotid sinus and carotid bifurcation branches, some of which may be relevant to baroreflex modulation (Capilupi et al. 2020; Lohmeier and Iliescu 2011; Iliescu et al. 2014). This placement would favor broader cardiovascular branch engagement compared to the more distal placement, discussed above. Placing a cuff further proximally, at or above 5 cm from the JF, would likely benefit selective cardiovascular neuromodulation less, given the anatomical separation between targeted branches (carotid sinus and bifurcation branches) and off-target branches (muscular, laryngeal sympathetic communicating branches). (Figure 9D).

Fourth, because recurrent laryngeal branches emerge below the clavicle on both sides, a device placed at or above the clavicle could target laryngeal fibers, e.g., for laryngeal pacing in spastic dysphonia, in which recurrent laryngeal nerve is intact (Friedman et al. 1989), while sparing more proximal sympathetic branches and most cervical vascular branches (Figure 9F). However, because aortic and lower visceral pathways remain distal to this level, these targets could still be co-recruited.

## Limitations

Our study has several limitations. Our cohort (n=28 donors, median age 88) is skewed toward older adults. Embalmed postmortem tissue may differ from in vivo nerve geometry in dimensions and compliance. Because donors were elderly and embalmed, absolute distances from the jugular foramen are not directly transferable to living patients without additional calibration. However, the proximal-to-distal ordering and landmark-relative spacing of branches should be assumed to carry over to living subjects. The precision of reported distance values is provided for reasons of completeness and does not imply millimeter precision, however a sub-centimeter scale may be relevant to nerve visualization during surgery.

The branch taxonomy relies on traced endpoints. In some cases, assigning a single target label to a branch with complex distal arborization required judgment calls, which we addressed through the multiple-targets category. Direct quantitative comparison with prior thoracic literature is limited because most studies do not report distances along the vagus trunk and use different reference frames. Our dissection also did not include abdominal course of the vagus nerve.

The exclusion of one of the donors from analysis due to unusual branching complexity highlights that a minority of individuals (here, 1 in 29) may present with an unusually complex vagal branching pattern in which discrete emergence zones are less demarcated; in such cases the landmark-registered windows we describe here would not be reliable. The map reported here is an anatomical reference for surgical planning rather than a directly transferable surgical coordinate frame. In a realistic clinical scenario, the windows would inform the initial surgical approach, but patient-specific confirmation of branch anatomy before or at the time of implantation will still be needed, e.g., using high-resolution ultrasound imaging (Settell et al. 2021), with calibration for estimating actual nerve dimensions (Dörschner et al. 2023) . High-resolution ultrasound resolves individual branch takeoffs, so in principle the branch-free interval around a landmark could be measured for a specific patient preoperatively, which is precisely the information needed to position a cuff within that individual’s available window rather than a population-average one.

Anatomical dissection was performed from the jugular foramen through the thoracic cavity and did not include the abdominal course of the vagus nerve. Below the diaphragm, the vagal trunk breaks up into esophageal and gastric plexuses, that give rise to hepatic, celiac, and other subdiaphragmatic branches. The complex plexiform branching of subdiaphragmatic nerves made it impossible for us to trace and reliably map branches using gross anatomical dissection methods. This limits the implications of our study, as any inference about gastrointestinal targets is beyond the region we mapped. Characterizing those branches within a landmark-registered framework, possibly using whole-body, high resolution imaging methods validated against gross dissection (Pelot et al. 2025), could extend the atlas to abdominal targets for neuromodulation.

A possible next step is to standardize translations between common thoracic operative landmarks (thoracic inlet, aortic arch, carina, hilum, hiatus) and our registered coordinate system, so that future comparisons can use either frame. Integrating finer branch-destination annotations would allow more direct alignment with studies distinguishing upper vs. lower pulmonary branches and with descriptions of esophageal plexus formation and vagal trunk variability.

## Acknowledgments

We are grateful to the individuals who chose to donate their bodies to academic medicine through the Anatomical Gift Programs of the Zucker School of Medicine, SUNY Upstate Medical University, and Lewis Katz School of Medicine of Temple University. Their donations made this study possible. Although this manuscript necessarily uses technical anatomical language, we recognize that these materials came from once-living human beings whose final act of generosity now supports medical education, anatomical discovery, and future patient care. We also extend our appreciation to their families and to the anatomical gift program staff who stewarded these donations with care and dignity.

We thank Drs. Kevin J. Tracey, Bruce T. Volpe, Ibrahim Mughrabi, Naveen Jayaprakash, Netanel Ben-Shalom and Harold Silverman, for thoughtful discussions that helped refine our understanding of vagus nerve anatomy, its functional organization, and its relevance to bioelectronic medicine. We are also grateful to the broader scientific and clinical community whose ongoing conversations around selective vagal neuromodulation helped shape the translational framing of this work.

This study was performed with partial support from NIH-NINDS 1R01NS136685 (SZ) and NIH-SPARC 75N98022C00019-P00003-9999–1 (SZ).

## Data Availability

The interactive atlas, underlying dataset, and figure-generating code are available at github.com/drsiyarb/vagus_nerve_explorer and archived on Zenodo (Bahadır 2026) (doi:10.5281/zenodo.21268191). The primary anatomical data (microCT, immunohistochemistry, and ultrasound) for all donors are available as a SPARC data collection (T. J. Levy 2026) (doi:10.26275/2vva-qmjw). The dissection protocol(Nassrallah et al. 2024) is available on protocols.io (doi:10.17504/protocols.io.14egn683ml5d/v1).

## Notes

https://github.com/drsiyarb/vagus_nerve_explorer

### Competing Interest Statement

The authors have declared no competing interest.

### Summary of Updates

1. Statistical analysis and reporting. Analyses were revised to address pseudoreplication and account for clustering within donors. Descriptive results are now reported as medians with interquartile ranges. Linear mixed-effects models were added for each branch group, with subgroup, side, sex, and their interactions as fixed effects and donor as a random intercept. New tables report adjusted means, 95% confidence intervals, intraclass correlation coefficients, exact p-values, and corrected contrasts. 2. Visualization and organization. A simplified bilateral reference schematic was added, showing the proximal-to-distal arrangement, variability, and overlap of major branch groups and common subgroups relative to surgical landmarks. Three repetitive Results sections were consolidated into a single group-based narrative, reducing them from approximately 3,300 to 1,450 words without removing data. Figure captions were shortened, and the dissection figure was enlarged and visually grouped by branch category. 3. Landmark registration and anatomical variability. The landmark-normalization procedure was clarified, and a worked example comparing short and long nerves before and after registration was added. Full ranges, outliers, landmark variability, donor-specific branch-free intervals, and classification performance for the cervical-thoracic transition are now reported. The revised text emphasizes that the identified zones represent population-level distributions rather than sharp anatomical boundaries. 4. Interpretation of anatomical separation. Claims of functional selectivity were replaced with the more precise concept of anatomical separation. The manuscript now explicitly states that gross branch emergence is an anatomical proxy, not evidence of fiber-level or electrophysiological selectivity. The Discussion was expanded to address fascicular organization, fiber composition, electrode geometry, electrode-tissue interactions, encapsulation, impedance, and stimulation waveform. 5. Translational scope and limitations. The manuscript now explains how surgeons, device developers, and anatomists may use the atlas while distinguishing anatomical opportunity from surgical accessibility. Expanded limitations address the elderly embalmed cohort, the excluded unusually branched donor, absence of subdiaphragmatic dissection, limited transferability of absolute distances, and the need for patient-specific anatomical confirmation. 6. Reproducibility and data quality. The interactive explorer's data schema, filters, linked views, and separation-index implementation are now described. The dataset, explorer, figure-generating code, and reproducible notebook are available through GitHub and an archived Zenodo snapshot. Several duplicated or carried-over segment-length values were reconciled with source records and corrected without changing the statistical results.

