## Supplementary Files for "A standardized, surgically relevant map for emergence of organ-specific branches from the human vagus nerve"

¹ Institute for Bioelectronic Medicine, Feinstein Institutes for Medical Research, Manhasset, NY, USA
 ² Elmezzi Graduate School of Molecular Medicine, Manhasset, NY, USA
 ³ Northwell Health, New Hyde Park, NY, USA
 ⁴ Donald and Barbara Zucker School of Medicine at Hofstra/Northwell, Hempstead, NY, USA
 ⁵ Penn State College of Medicine, Hershey, PA, USA
 ⁶ Lewis Katz School of Medicine of Temple University, Philadelphia, PA, USA
 ⁷ Hackensack Meridian School of Medicine, Nutley, NJ, USA

### Supplementary Methods

*Details of Anatomical Dissection*

The vagus nerve and its cardiovascular targets were isolated, bilaterally, in the cervical region. After disarticulation of craniovertebral junction, and hyperflexion of the head, the vagus nerve was exposed within carotid sheath at the parapharyngeal space. Neck and appropriate thorax dissections were performed along with the disarticulation to expose the vagus from the jugular foramen to the thoracic cavity. We defined “cervical” rather than “cranial” to be the point at which the superior laryngeal nerve (SLN) emerged from the vagal trunk. Cranial to the emergence of the SLN, the vagus nerve shared a connective tissue sheath with the accessory and hypoglossal nerves. To avoid mistakenly attributing a branch to the vagus, we excluded branches emerging from the cranial-most segment of the vagus. Once the vagus had been exposed, sutures were placed, bilaterally, into the epineurium of the vagus nerve at the levels of the jugular foramen, superior laryngeal prominence, carotid bifurcation, and superior border of the clavicle as landmark indicators. We systematically followed each branch of the vagus to its endpoints, carefully documenting the target endpoint of each branch. If a branch innervated more than one target, those multiple endpoints were documents and termed *Multiple Targets*. The endpoint target of each branch was documented in writing, in excel data sheets, by video and photographically.

In detail, we approached the vagus nerve from the anterolateral direction. The vagus nerve was initially exposed and identified in the cervical region (defined as the point at which the superior laryngeal nerve emerged from the vagus, to the superior border of the clavicle). Two identical cervical skin flaps were created. The superior edge of the skin flap extended from the inferior edge of the mandible towards the area inferior to the ear lobe. The inferior edge of the skin flap extended from the jugular notch, along the superior edge of the clavicle, towards the acromion process. The two skin flaps were connected midline over the laryngeal prominence and reflected laterally, remaining superficial to the platysma to preserve branches of cervical spinal nerves. Once these nerves were identified, the sternocleidomastoid muscle was detached from its inferior attachments and reflected superolaterally to expose the carotid sheath. At this point, the spinal accessory nerve was identified (and marked using a green colored suture) and the submandibular gland was excised to expose the hypoglossal nerve (the latter marked with a black colored suture). The ansa cervicalis branches were identified (and marked with a blue tissue dye), as were any other branches to larygyneal muscles from the vagus nerve (as presented in the results section, there were some direct branches to the larygyneal muscles from the vagus nerve). Thereafter, the carotid sheath was opened to expose the vagus nerve. Once the vagus was exposed, sutures were placed bilaterally into the epineurium of the vagus nerve at the level of the carotid bifurcation, superior laryngeal prominence, and superior border of the clavicle, as landmark indicators. The level of the jugular foramen was also labeled with a black suture, when the jugular foramen was eventually exposed.

The thoracic region of the vagus was exposed by creating a thoracoabdominal skin flap, as described previously(Shannon Knutson et al. 2023). Once exposed, both pectoralis major and minor muscles were detached inferiorly and reflected superolaterally. The rib cage was opened by cutting the inferior ribs (beginning at the 11th rib) along the border created by the midaxillary line and skin flap. The second and first ribs, and clavicle, were cut with large bone pliers. Once the rib cage is mobile, superficial fascial layers were separated from the sternum and the rib cage reflected inferiorly. The thoracic dissection began with removal of the lungs and heart, with incisions made approximately 1 cm from the root of the lung. Posterior portions of the middle and inferior lobes were left intact for later examination. A midline incision was made through the pericardium and following the diaphragmatic surface of the heart to separate it from the inferior vena cava. The heart was tilted anteriorly to separate it from the pulmonary vessels posteriorly, before a horizontal incision was made across the great vessels near the base of the heart. Once the heart was removed from the pericardium, anterior and lateral portions of the pericardium were cut. The posterior portion of the pericardium was maintained to preserve the deeper lying esophageal plexus.

Before further manipulating the thoracic vagus nerves, sutures are placed bilaterally into the epineurium of the vagus at the level of the previously marked superior border of the clavicle (if not marked already). The right vagus was followed as it coursed deep to the brachiocephalic vein, making note of the location right recurrent laryngeal nerve as it looped behind the right subclavian artery. The right vagus was followed into the space deep to the superior vena cava to the entrance of the azygos vein. The right esophageal plexus was identified and dissected out from the surrounding fascia posterior to the lung root. The posterior pericardium was slowly dissected from the esophageal plexus at this point. The left vagus was followed deep to the brachiocephalic vein and onto the aorta, making note of the left recurrent laryngeal nerve as it looped under the aortic arch. Careful dissection on both sides along the pericardium, mediastinum and lungs was performed, sparing all branches from the vagus to target end organs.

The vagus was further dissected using a posterior pharyngeal approach. To prepare for this dissection, a laminectomy is performed of the cervical spine. The spinous processes of the upper thoracic and lower cervical spine are palpated to form the midline cut of two large flaps to expose the cervical vertebra. This midline cut extended from the superior portion of the scapula to the occipital protuberance. The flaps then extended approximately 5 cm laterally on both sides to create two flaps that were removed along with the underlying neck and upper back musculature. This flap was as deep as the transverse processes, bilaterally, to avoid damaging the cervical spinal nerve roots as they emerged from the vertebral foramina and column. Once the cervical vertebrae from C2 - C7 were exposed, a bone saw was used to cut into the laminae just lateral to the spinous processes and just deep enough to avoid damaging the underlying spinal cord, rootlets and dorsal root ganglia. After this portion of the vertebral column was removed, the laminae were slowly removed using a hammer, chisel and bone shears.

After the laminectomy was performed, the cranial, cervical, and thoracic portions of the specimen were removed via decapitation/disarticulation to create an extractable tissue block. From the anterior side, the esophageal plexus was cut between the bottom of the lung roots and the esophageal hiatus to allow for separation of the aorta and esophagus from the body. Upon leaving the thoracic cavity, the cervical block with the vagus in situ was now separated from the cadaver by cutting through the brachial plexus and surrounding musculature. Posteriorly, the cervical spine was separated at the bottom of the laminectomy, allowing for the vertebral bodies to come forward with the tissue block. Once the tissue block is formed, further reductions were made via a craniotomy and several coronal cuts. These cuts were made through the posterior cranial fossa within 1-2 cm of the jugular foramen and through the anterior cranial fossa so that the anterior portion of the pharynx is preserved. The vertebral bodies of C2 - C5 were removed, as was C1, to expose the posterior pharynx. Using a hammer, chisel and bone shears, the vagus was followed cranially to the jugular foramen into the carotid sinus. The level of the jugular foramen was marked with a suture in the epineurium of the vagus at this time).

The number of branches per vagus nerve was documented, as was the point of emergence of each branch from the vagus nerve relative to the jugular foramen. We systematically followed each branch of the vagus to its endpoint(s). Sutures of various colors and sizes, and tissue marking dyes (22-050-455, FisherScientific, Epredia™ Mark-It™ Tissue Marking Dye kit, Richard-Allan Scientific LLC (a subsidiary of Epredia), Kalamazoo, MI, USA) were used as fiduciary markers for the various subsets of vagal nerve branches. After marking, the vagus nerve and its branches were recorded manually, by video and photodocumented, both prior to and after removal and segmentation.

Rationale For group aggregation

The rationale for group aggregation came from two complementary frameworks. From an evolutionary perspective, the circumpharyngeal ridge has been proposed to define the head–trunk interface, with the venous pole (embryological precursor of the heart) marking this boundary in crown gnathostomes (Higashiyama et al. 2016; Hirasawa et al. 2016). Developmentally, this framework is supported by the human embryology literature, which describes the pharyngeal arches as contributing to the neurovascular and musculoskeletal anatomy of the head, neck, and upper thorax(Frisdal and Trainor 2014). We therefore used the term Pharyngeal Arch Target-related domain (or cervical domain), for sympathetic, muscular, and vascular targets, including recurrent laryngeal and aortic branches. Cardiac, pulmonary, and esophageal targets are grouped separately as the Primitive Mediastinum Target-related domain (or thoracic domain) reflecting their caudal visceral/thoracic anatomy.

The cervical domain is then further divided into three longitudinal zones (separate from the segments that we defined by the landmarks) because its constituent muscular and vascular targets did not distribute along the length continuously, but instead showed separable proximal, intermediate, and distal emergence regions along the vagus. These positional subdivisions are termed the superior cervical, mid/low cervical, and upper thoracic zones. This zone based partitioning is also consistent with pharyngeal arch developmental anatomy, in which different vagus-associated laryngeal territories are linked to different caudal pharyngeal arches; for example, cricothyroid/superior laryngeal territory is associated with the fourth arch, whereas intrinsic laryngeal/recurrent laryngeal territory is associated with the sixth arch(Frisdal and Trainor 2014). These domains and zones did not replace the original group/subgroup taxonomy, but used as an additional analysis for testing regional topography along the vagus nerve.

Some of the cadaveric branch sets are publically available on the University of Pennsylvania, Pennsieve Platform for Data Management at https://discover.pennsieve.io/datasets:

Stavros Zanos, Naveen Jayaprakash, Qanud Khaled, Zeinab Nasrallah, Mary Barbe, Frank Lui Chen, Larry Miller, Theodoros Zanos, Todd J Levy, M.S., Avantika Vardhan, Jinxuan Cang, Viktor Toth, Kevin Coppa, Netanel Ben-Shalom, Weiguo Song, Nicole Carpentiere, Theofilos Kanavos, Effrosyni Birbas, Siyar Bahadir, Parisa Saleknezhad. Dataset published for SPARC program, “Human vagus nerve anatomical reconstruction using microCT immunohistochemistry and ultrasound” - f004, Pennsieve Discover. https://discover.pennsieve.io/datasets/505. Published Nov 14, 2025

Stavros Zanos, Naveen Jayaprakash, Qanud Khaled, Zeinab Nasrallah, Mary Barbe, Frank Lui Chen, Larry Miller, Theodoros Zanos, Todd J Levy, M.S., Avantika Vardhan, Jinxuan Cang, Viktor Toth, Kevin Coppa, Netanel Ben-Shalom, Weiguo Song, Siyar Bahadir, Nicole Carpentiere, Parisa Saleknezhad. Dataset published for SPARC program, “Human vagus nerve anatomical reconstruction using microCT immunohistochemistry and ultrasound” - f006, Pennsieve Discover. https://discover.pennsieve.io/datasets/438. Published June 9, 2025.

Stavros Zanos, Naveen Jayaprakash, Qanud Khaled, Zeinab Nasrallah, Mary Barbe, Frank Lui Chen, Larry Miller, Theodoros Zanos, Todd J Levy, M.S., Avantika Vardhan, Jinxuan Cang, Viktor Toth, Kevin Coppa, Netanel Ben-Shalom, Weiguo Song, Nicole Carpentiere, Theofilos Kanavos, Effrosyni Birbas, Siyar Bahadir, Parisa Saleknezhad. Dataset published for SPARC program, “Human vagus nerve anatomical reconstruction using microCT immunohistochemistry and ultrasound” - f008, Pennsieve Discover. <https://discover.pennsieve.io/datasets/512> Published Nov 14, 2025

Stavros Zanos, Naveen Jayaprakash, Qanud Khaled, Zeinab Nasrallah, Mary Barbe, Frank Lui Chen, Larry Miller, Theodoros Zanos, Todd J Levy, M.S., Avantika Vardhan, Jinxuan Cang, Viktor Toth, Kevin Coppa, Netanel Ben-Shalom, Weiguo Song, Nicole Carpentiere, Theofilos Kanavos, Effrosyni Birbas, Siyar Bahadir, Parisa Saleknezhad. Dataset published for SPARC program, “Human vagus nerve anatomical reconstruction using microCT immunohistochemistry and ultrasound” - f009, Pennsieve Discover. <https://discover.pennsieve.io/datasets/513>. Published Nov 14, 2025

Stavros Zanos, Naveen Jayaprakash, Qanud Khaled, Zeinab Nasrallah, Mary Barbe, Frank Lui Chen, Larry Miller, Theodoros Zanos, Todd J Levy, M.S., Avantika Vardhan, Jinxuan Cang, Viktor Toth, Kevin Coppa, Netanel Ben-Shalom, Weiguo Song, Nicole Carpentiere, Theofilos Kanavos, Effrosyni Birbas, Siyar Bahadir, Parisa Saleknezhad. Dataset published for SPARC program, “Human vagus nerve anatomical reconstruction using microCT immunohistochemistry and ultrasound” - f011, Pennsieve Discover. <https://discover.pennsieve.io/datasets/514>. Published Nov 14, 2025

Supplementary Results

#### 1. Prevalence and pattern of emergence of vagal branches

The final cohort comprise of 28 donors (15 female, 13 male; median age 88 years, range 57–90+) and 56 vagus nerves (28 left, 28 right; Table 1 The carotid bifurcation landmark is located at 6.25 cm from the jugular foramen (range 2.70–10.10), the laryngeal prominence at 8.62 cm (range 5.41–12.10), and the superior border of the clavicle at 15.60 cm (range 9.68–22.50) (Table 1). Total nerve length is 32.65 cm (range 24.50–44.70), similar for left and right nerves. A total of 2,177 branches (997 left, 1,180 right) branches are identified and distances to jugular foramen and other anatomical landmarks are measured. Branch emergence spans 0.20–43.55 cm (native distance) from the jugular foramen (mean 17.52 cm; native distance); after landmark-based scaling and normalization, registered emergence distances spans 0.32–32.68 cm (mean 17.41 cm; Table 1).

Most vagal branches emerge in the thoracic region, below the clavicle (61.1% of branches). Branches of the cervical vagus are commonly found in the upper cervical segment, from the jugular foramen to the carotid bifurcation (18.1%), followed by the lower cervical segment, from the laryngeal prominence to the superior border of the clavicle (14.0%), and the fewest in the mid-cervical segment, between the carotid bifurcation and the laryngeal prominence (6.8%) (Table 1). Likewise, branching density is highest in the thorax (1.44 branches/cm/nerve), followed by the upper cervical (1.07) and mid-cervical (0.92) segments, and the lowest in the lower cervical segment (0.76) (Table 1). Additional details on the emergence of branches from different levels of the right and left vagus nerves are in Supplementary Tables S2 and S4, with Supplementary Table S2 including sex stratified data.

Branches are classified in one of 7 groups, according to the organ or structure of termination: sympathetic, muscular, vascular, cardiac, pulmonary, esophageal and multiple targets. The largest groups are esophageal (n=522), vascular (n=499), and cardiac (n=460). Emergence of these groups exhibits a proximal-to-distal ordering along the nerve (Figure 2; Table 2). Sympathetic branches are closest to the jugular foramen (mean distance 5.28 cm), followed by muscular (9.59 cm), vascular (10.70 cm), and multiple-target branches (11.06 cm), and more distally cardiac (19.65 cm), pulmonary (25.36 cm), and esophageal (26.57 cm) branches. Side differences in mean emergence distance are modest, yet are most pronounced for multiple-target branches (mean L–R difference = -5.15 cm; more proximal emergence on left vs. right), vascular branches (+2.74 cm), and pulmonary branches (+2.08 cm). Details of branch emergence site distances for right vs left, and female vs male subjects, are reported in Supplementary Table S2.

Most groups include several, anatomically-defined subgroups (Figure 4). For each sub-group, we registered the frequency for left side-only presence (present on left, absent on right in a given donor), right-only presence, bilateral presence (present on both sides in the same donor), and total presence (present on either side). Three subgroups were present in all donors: SLN, RLN, and cardiopulmonary. Other highly prevalent subgroups included superior cervical ganglion (96%), esophageal plexus (93%), common carotid (86%), pulmonary (86%), superficial cardiac plexus (82%), and pharyngeal (79%). Among the 25 common subgroups reported here, the least prevalent subgroups were cardiac (from esophageal plexus) (21%), cardiopulmonary (from esophageal plexus) (29%), ansa cervicalis (32%), general cardiac (36%), and laryngeal (39%) (Figure 3).

Frequent bilateral presence isnoted for the major laryngeal branches (SLN and RLN, 96% bilateral each), esophageal plexus (86%) and cardiopulmonary (82%). In contrast, left-biased pattern is seen for the aorta (left-only (L) 54%, right-only (R) 4%, bilateral (B) 0%) and the superficial cardiac plexus (L 57%, R 7%, B 18%). Examples of right-biased sub-groups includes carotid bifurcation (R 21%, L 7%), sympathetic trunk (R 25%, L 14%), and carotid sinus (R 25%, L 14%). Some subgroups are unilateral without side bias (e.g., deep cardiac plexus: L 14%, R 14%) (Figure 3). Detailed data, stratified by side and sex, are in Supplementary Figure 7 and Supplementary Table S3.

##### Sympathetic Group

Within the sympathetic group (n=199), most branching targets the Superior Cervical Ganglion (n=143; pooled mean 4.06 cm, range 0.32–11.17), with left SCG branching more proximal than right (Left 3.73 [0.58–8.05] vs Right 4.41 [0.32–11.17]) (Figure 4A, Supplementary Table S2). Sympathetic trunk branching is less frequent but more distal (n=52; pooled mean 8.85 cm, range 0.91–19.48), again with a leftward proximal shift (Left 8.11 [2.78–19.48] vs Right 9.18 [0.91–15.32]). A rare mixed label indicating branching to both trunk and SCG is observed (sympathetic trunk and superior cervical ganglion, n=4 female donor bodies; pooled mean 2.69 cm, range 2.00–3.59; see Supplementary Table S2). No statistical significant difference is noted between male and female subjects in terms of branch emergence.

##### Muscular Group

Within the Muscular group (n=214), branching spanns a broad proximodistal range (Figure 4B, Supplementary Table S4). Pharyngeal and SLN branching occursproximally (Pharyngeal: n=46, mean 3.01 cm, range 0.77–7.97; SLN: n=58, mean 3.94 cm, range 1.45–6.35), whereas Ansa Cervicalis and General Laryngeal branching occurs more distally (Ansa cervicalis: n=28, mean 8.47 cm, range 3.78–14.79; General Laryngeal (one right-sided non-recurrent laryngeal nerve branch from Subject 2305 is added to this category): n=26, mean 12.41 cm, range 2.18–21.98). RLN branching formed the most distal muscular distribution (RLN: n=55, mean 20.20 cm, range 13.64–28.23) and showed marked laterality (Left RLN: mean 22.84 cm, range 18.38–28.23; Right RLN: mean 17.47 cm, range 13.64–20.46).

Side-specific values for the remaining muscular subgroups are as follows: Pharyngeal branching showed a slight leftward proximal shift (Left n=20, mean 2.74 cm, range 1.32–5.18 vs Right n=26, mean 3.21 cm, range 0.77–7.97; Δmean (L–R) = -0.47 cm). SLN branching is similar between sides (Left n=29, mean 3.89 cm, range 1.45–6.35 vs Right n=29, mean 3.98 cm, range 2.31–5.84; Δmean (L–R) = -0.09 cm). Ansa cervicalis branching is slightly more distal on the left (Left n=16, mean 8.99 cm, range 3.78–14.79 vs Right n=12, mean 7.77 cm, range 4.00–11.13; Δmean (L–R) = +1.22 cm). General Laryngeal branching showed a modest rightward distal shift (Left n=11, mean 11.92 cm, range 9.82–12.84 vs Right n=15, mean 13.08 cm, range 5.66–21.19; Δmean (L–R) = -1.16 cm). No statisticall significant difference is noted between male and female subjects in terms of branch emergence.

##### Vascular Group

Within the Vascular group (n=499), branching spans a broad range with subgroup-specific patterns, with data shown in Figure 4C, Supplementary Tables S4. Proximal vascular branching includes Carotid Sinus (n=34; mean 5.29 cm; range 1.18–11.88) and Internal Carotid Artery (n=61; mean 5.43 cm; range 0.42–15.28). Carotid Bifurcation branching occurrs at slightly more distal positions (n=40; mean 6.36 cm; range 3.43–13.56). Multiple Branches To Carotid Artery are present (n=74; mean 8.21 cm; range 1.00–20.93) and spans a wide range. More distally, branches to the Common Carotid (n=117; mean 10.99 cm; range 3.17–23.87) and Internal Jugular Vein (n=65; mean 10.84 cm; range 2.09–20.32) are observed. General Vascular branching occurrs at more distal positions (n=65; mean 15.97 cm; range 5.06–22.45), bridging the cervicothoracic range. Branches to the Aorta represented the most distal vascular subgroup (n=42; mean 22.16 cm; range 16.68–29.24), as expected. A single External Carotid Artery branch is observed (n=1, male donor body) and is treated as a rare occurrence (Supplementary Tables S2, S4).

Laterality effects across vascular branching are generally modest, but several subgroups showed side differences in mean branching location: Carotid Sinus branching is more distal on the right (Left n=11, mean 5.03 cm, range 3.63–8.91 vs Right n=23, mean 6.73 cm, range 2.31–19.48; Δmean (L–R) = -1.70 cm), whereas Internal Carotid Artery branching is more distal on the left (Left n=24, mean 6.98 cm, range 2.16–16.04 vs Right n=37, mean 4.95 cm, range 1.58–14.22; Δmean (L–R) = +2.03 cm). Carotid Bifurcation branching is also slightly more distal on the left (Left n=17, mean 6.96 cm, range 2.31–17.58 vs Right n=23, mean 5.84 cm, range 0.58–11.65; Δmean (L–R) = +1.12 cm). In contrast, Multiple Branches To Carotid Artery, Common Carotid, and Internal Jugular Vein showed minimal side differences (Δmean (L–R) = +0.20 cm, +0.19 cm, and -0.28 cm, respectively). General Vascular branching is slightly more distal on the right (Left n=37, mean 15.05 cm, range 6.35–29.08 vs Right n=28, mean 16.86 cm, range 15.46–22.52; Δmean (L–R) = -1.81 cm), whereas Aortic branching remained strongly left-sided and more distal on the left (Left n=38, mean 21.44 cm, range 9.82–29.08 vs Right n=4, mean 19.49 cm, range 18.72–20.53; Δmean (L–R) = +1.95 cm). No statisticall significant difference is noted between male and female subjects in terms of branch emergence.

##### Cardiac Group

Within the Cardiac group (n=459 with valid registered distances), branching is predominantly below the clavicle (81.0%) (Figure 4D). Data is also provided in Supplementary Tables S4. The cranial-to-clavicle (surgically accessible) component accounted for (19.0%) and lay almost entirely between the laryngeal prominence and the clavicle (17.9%); only (1.1%) cardiac branches occur cranial to the laryngeal prominence. Importantly, the accessible cardiac component is not exclusively the superficial cardiac plexus: among cardiac branches above the clavicle (n=87), the Superficial Cardiac Plexus contributes 49.4%, Cardiopulmonary contributes 36.8%, with smaller contributions from General Cardiac (8.0%) and Deep Cardiac Plexus (5.7%). A single entry called Cardiac Plexus (1/87, 1.1%) is added to Superficial Cardiac Plexus. (Figure 4D, Supplementary Tables S4).

At the subgroup level, Cardiopulmonary branching is the most prevalent (n=292; mean 20.79 cm; range 9.92–31.34) but had a limited above-clavicle component (11.0%) - and those Cardiopulmonary branches above the clavicle occurs exclusively on the right (0/76 left vs 32/216 right; 14.8% of right Cardiopulmonary branching). Superficial Cardiac Plexus branching is more proximal (n=92; mean 16.28 cm; range 5.46–26.41) and contributes a larger accessible fraction (45.7% above the clavicle), with most of the left-sided accessible cardiac branching coming from this subgroup (32 of 41 above-clavicle left branches). Deep Cardiac Plexus (n=42; mean 19.67 cm; range 9.58–27.02) and General Cardiac (n=32; mean 19.65 cm; range 12.82–26.90) are smaller and largely thoracic, with modest above clavicle components (Deep Cardiac: 11.9%; General Cardiac: 7/32, 21.9%). Together, these results highlight a surgically relevant asymmetry: left-sided accessible cardiac branching targets overwhelmingly Superficial Cardiac Plexus, whereas the right side uniquely added a meaningful Cardiopulmonary contribution above the clavicle (and a smaller Deep Cardiac Plexus contribution) (Figure 4D, Supplementary Tables S4 ).

Side-specific subgroup values are as follows: Superficial Cardiac Plexus branching is slightly more distal on the right (Left n=65, mean 15.93 cm, range 9.82–24.35 vs Right n=27, mean 16.82 cm, range 13.06–19.32; Δmean (L–R) = -0.89 cm). Deep Cardiac Plexus branching is more distal on the left (Left n=13, mean 22.02 cm, range 19.78–29.08 vs Right n=29, mean 19.00 cm, range 12.84–23.34; Δmean (L–R) = +3.02 cm). General Cardiac branching showed a modest rightward distal shift (Left n=18, mean 18.41 cm, range 9.82–29.08 vs Right n=14, mean 19.72 cm, range 18.72–20.53; Δmean (L–R) = -1.31 cm). Cardiopulmonary branching is more distal on the left overall (Left n=76, mean 22.46 cm, range 19.78–29.08 vs Right n=216, mean 19.14 cm, range 8.89–24.35; Δmean (L–R) = +3.32 cm). A single Cardiac Plexus branch is observed on the left (n=1, mean 11.33 cm, range 11.33–11.33) and is treated as a rare occurrence. No statisticall significant difference is noted between male and female subjects in terms of branch emergence.

##### Pulmonary Group

The Pulmonary group contained 239 branches. Pulmonary branching is entirely below the clavicle (239/239, 100%), with a pooled mean registered distance of 25.36 cm (range 17.73–32.68) (Figure 4E, Supplementary Tables S4 and S5). Laterality showed a distal shift on the left (Left n=110; 26.48 cm [19.93–32.68]) compared with the right (Right n=129; 24.40 cm [17.73–31.84]), yielding Δmean (L–R) = +2.08 cm. No statisticall significant difference is noted between male and female subjects in terms of branch emergence.

##### Esophageal Group

There are 522 esophageal-related branches (Left n=202, Right n=320). All esophageal branching is below the clavicle on both sides. Overall registered distance is 26.57 cm (range 16.02–32.68). There is a distal shift on the left (Left 27.26 [20.92–32.68] vs Right 26.14 [16.02–32.62]; Δmean (L–R) = +1.12 cm) (Figure 4F, Supplementary Tables S4).

At the subgroup level, branches to Esophageal Plexus are the most common (n=259; mean 27.53 [16.02–32.68]). General Esophageal branches (direct branches to the esophagus from the vagus) are also frequent (n=117; mean 24.57 [17.53–32.25]) and are more proximal than Esophageal Plexus branching (Figure 4F, Supplementary Tables S4). Branching patterns that contribute to other systems are also observed after esophageal plexus formation and are reported as explicitly derived labels: Pulmonary (from Esophageal Plexus) (n=99; mean 26.60 [22.65–32.62]), Cardiopulmonary (from Esophageal Plexus) (n=23; mean 25.28 [19.59–31.95]), and less frequent Cardiac (from Esophageal Plexus) branching (n=24; mean 27.18 [22.92–32.23]). Side differences are most pronounced for Cardiopulmonary (from Esophageal Plexus) (Δmean +3.32 cm, left more distal) and Cardiac (from Esophageal Plexus) (Δmean −3.38 cm, right more distal; small n), while Pulmonary (from Esophageal Plexus) showed only a small side difference (Δmean −0.33 cm, right slightly more distal) (Figure 4F, Supplementary Table S4).

For all data in Figure 4, female and male stratified data are presented in (Supplementary Figure 1 and Supplementary Table S2). No statistical significant difference is noted between male and female subjects in terms of branch emergence.

**Supplementary Figure 1**

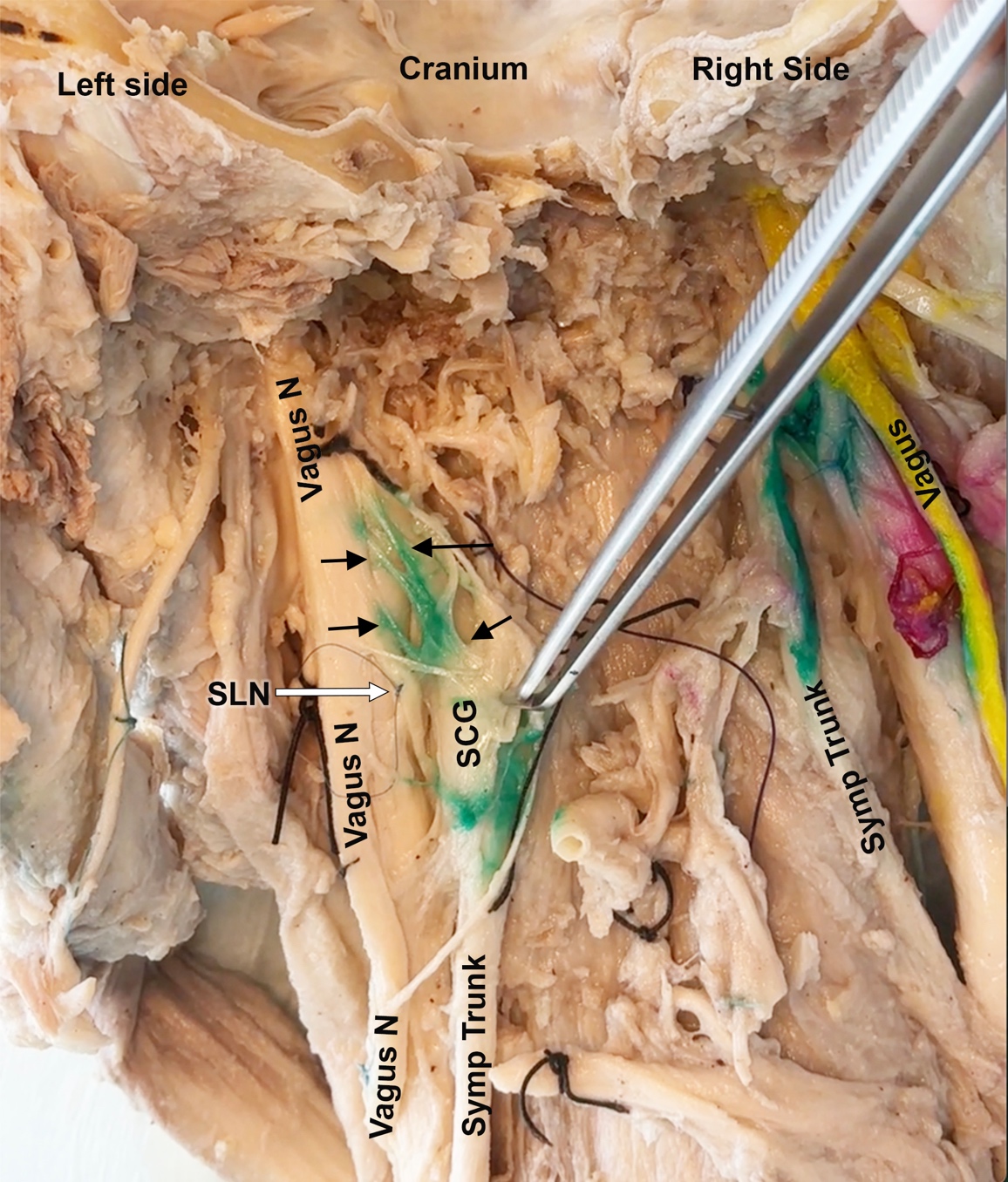

Figure 1. Photographic image of connections between the vagus nerve and sympathetic structures in cervical region. Cranium, right and left sides are indicated in this posterior view of the cervical region. The posterior side of the vagus nerve (N) is colored yellow in this image. Forceps are used to pull the superior cervical ganglion (SCG) towards the midline, showcasing branches connecting the vagus and SCG (black arrows, colored green). Abbreviations: N = nerve, SCG = superior cervical ganglion, SLN = superior laryngeal nerve (demarcated by a small pale blue suture and white arrow), Symp Trunk = sympathetic trunk. Black sutures mark cervical spinal roots.

**Supplementary Figure 2**

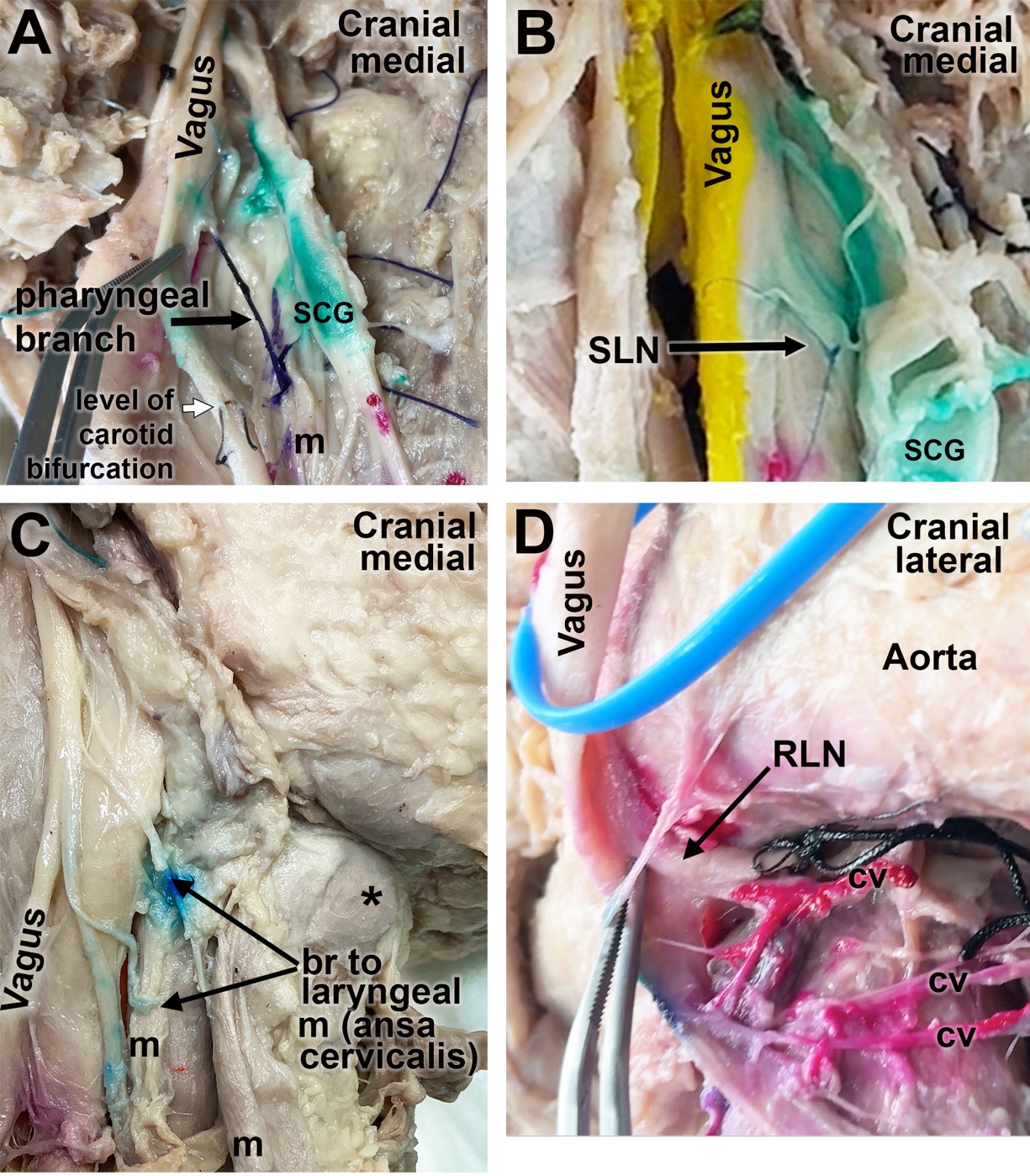

Figure 2. Photographic image of muscular branches from vagus nerve in cervical and upper thoracic regions. Cranial and medial or lateral sides of regions, and vagus nerve, are indicated. (A) Posterior view of upper cervical region showing branches connecting the vagus nerve to pharyngeal muscles (m) (branch is dyed purple, black arrow), superior cervical ganglion (SCG; dyed green). The level of the carotid bifurcation is also indicated (white arrow, small black suture). (B) Posterior side of vagus (dyed yellow) gives rise to the superior laryngeal nerve (SLN, small pale blue suture and black arrow) and connections to sympathetic structures (dyed green). (C). Anterior view of vagus showing branches (br) to laryngeal muscles (m), some presumedly carried by the ansa cervicalis branches (which are C1) that are hitchhiking on the vagus nerve. *: laryngeal prominence. (D) Anterior view of the vagus (blue vessel loop) as it wraps around the aorta. Here, the vagus nerves gives rise to the recurrent laryngeal nerve (RLN, black arrow and larger knotted black suture) and cardiovascular branches (cv) passing to the deep cervical plexus.

**Supplementary Figure 3**

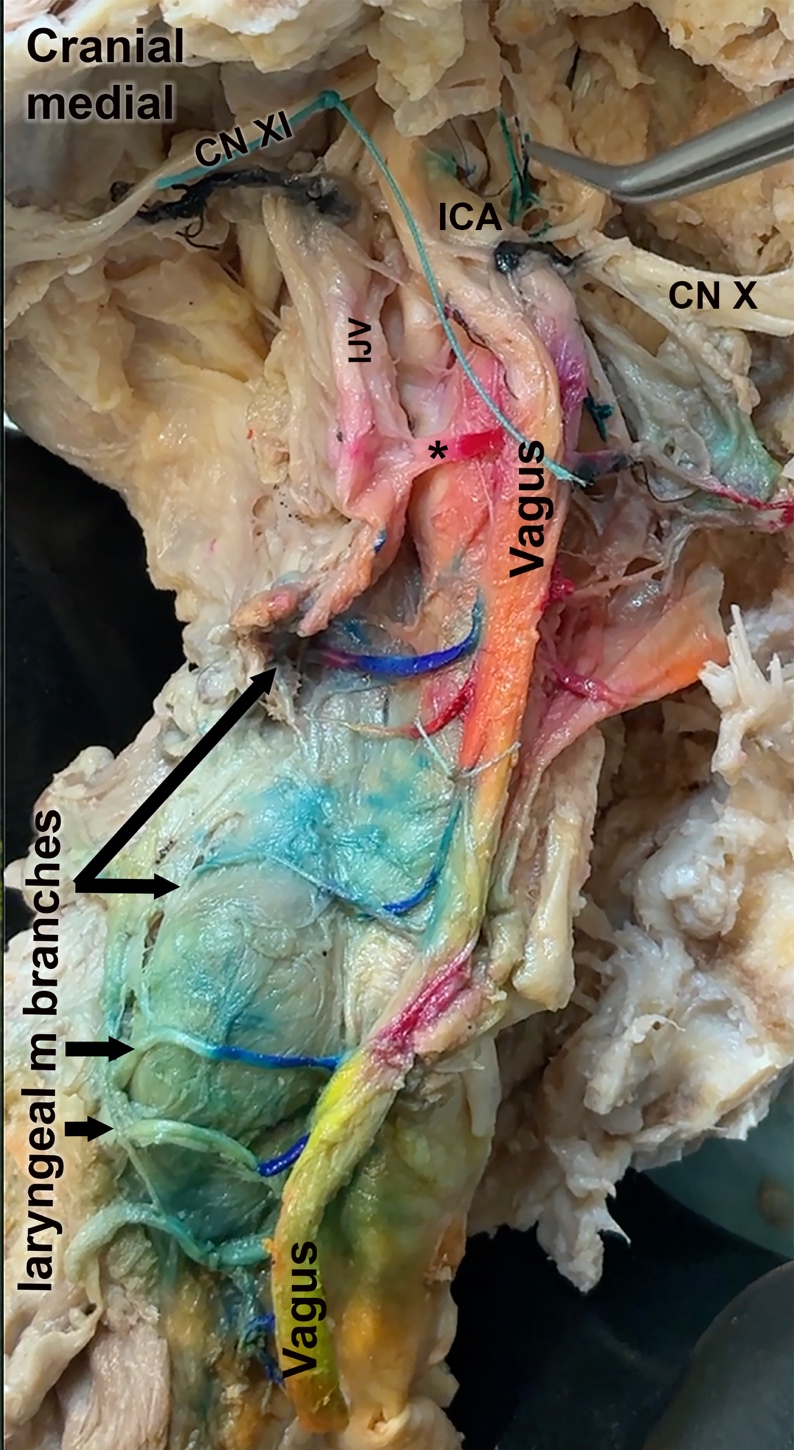

Figure 3. Photographic image of muscular branches from vagus nerve in cranial and cervical regions. Cranial and medial sides are indicated, as is vagus nerve (orange dye indicates anterior side in this image; yellow dye indicates the posterior side, both sides visible due to a twisting of the vagus as it descends towards the thorax). Spinal accessory nerve (CN XI, larger green suture), hypoglossal nerve (CN X), internal carotid artery (ICA), internal jugular vein (IJV), and laryngeal muscle (m) branches (blue and faded blue color) are indicated. * and red dyed branch delineates a cardiovascular branch from the vagus to the internal jugular vein.

**Supplementary Figure 4**

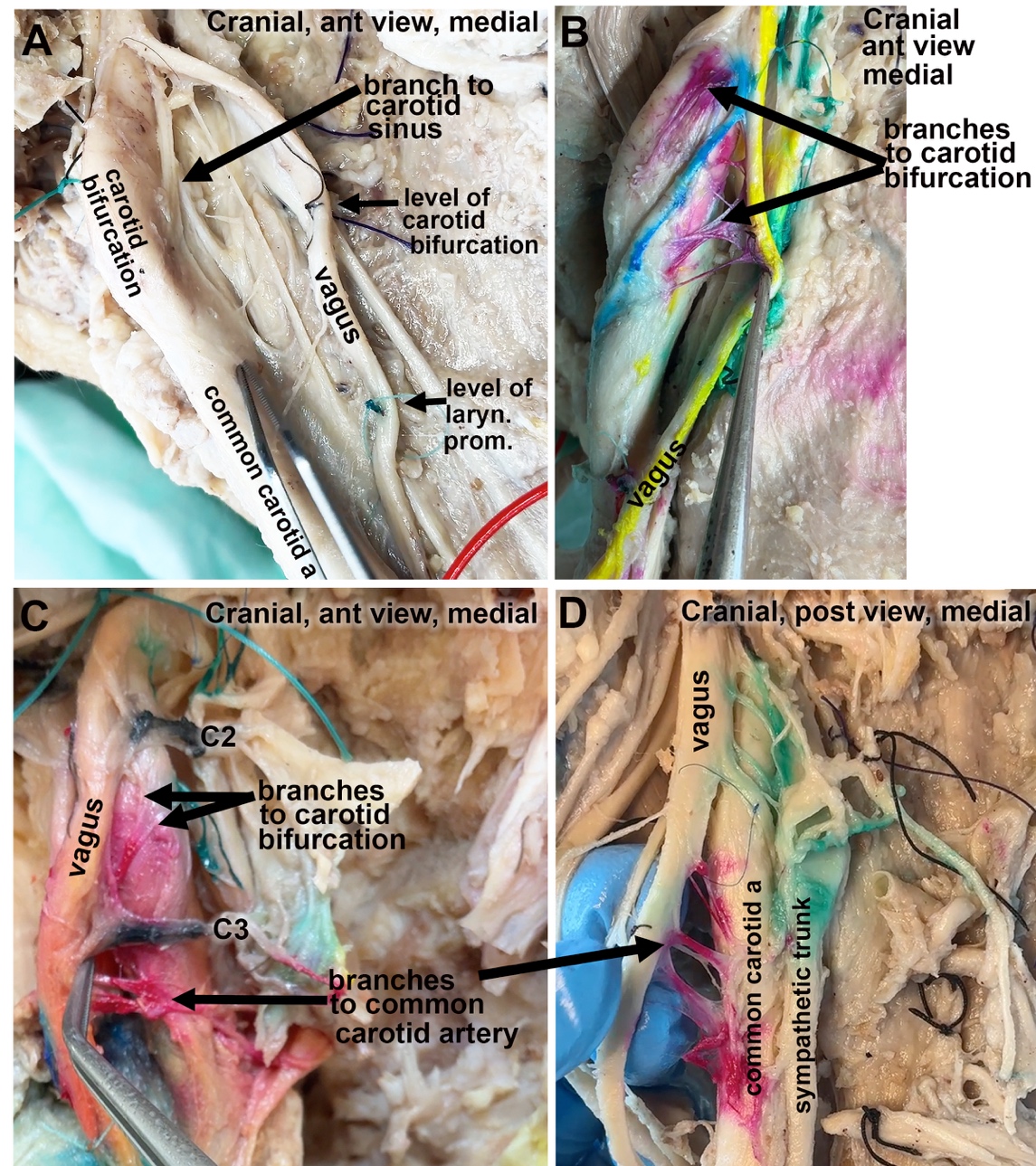

Figure 4. Photographic image of vascular branches from vagus in cranial and cervical regions. Cranial, medial side, anterior vs. posterior (post) views are indicated. (A) Image depicting carotid bifurcation, branch from vagus to carotid sinus, a small black suture marking the level of carotid bifurcation on the vagus nerve, and a blue suture marking the level of laryngeal prominence on the vagus nerve. (B) Anterior side of vagus marked with yellow dye. Branches connecting the vagus nerve to the carotid bifurcation of carotid artery (i.e., cardiovascular branches; dyed red), and laryngeal muscles (dyed blue). (C) Branches connecting the vagus with cranial (C) roots of C2 and C3 (dyed black), carotid bifurcation (dyed red, thick black arrow), and common carotid artery (dye red and thinner black arrow). (D) Branches between the vagus and the sympathetic ganglion and trunk (dyed green), and common carotid artery (dyed red and long black arrow).

**Supplementary Figure 5**

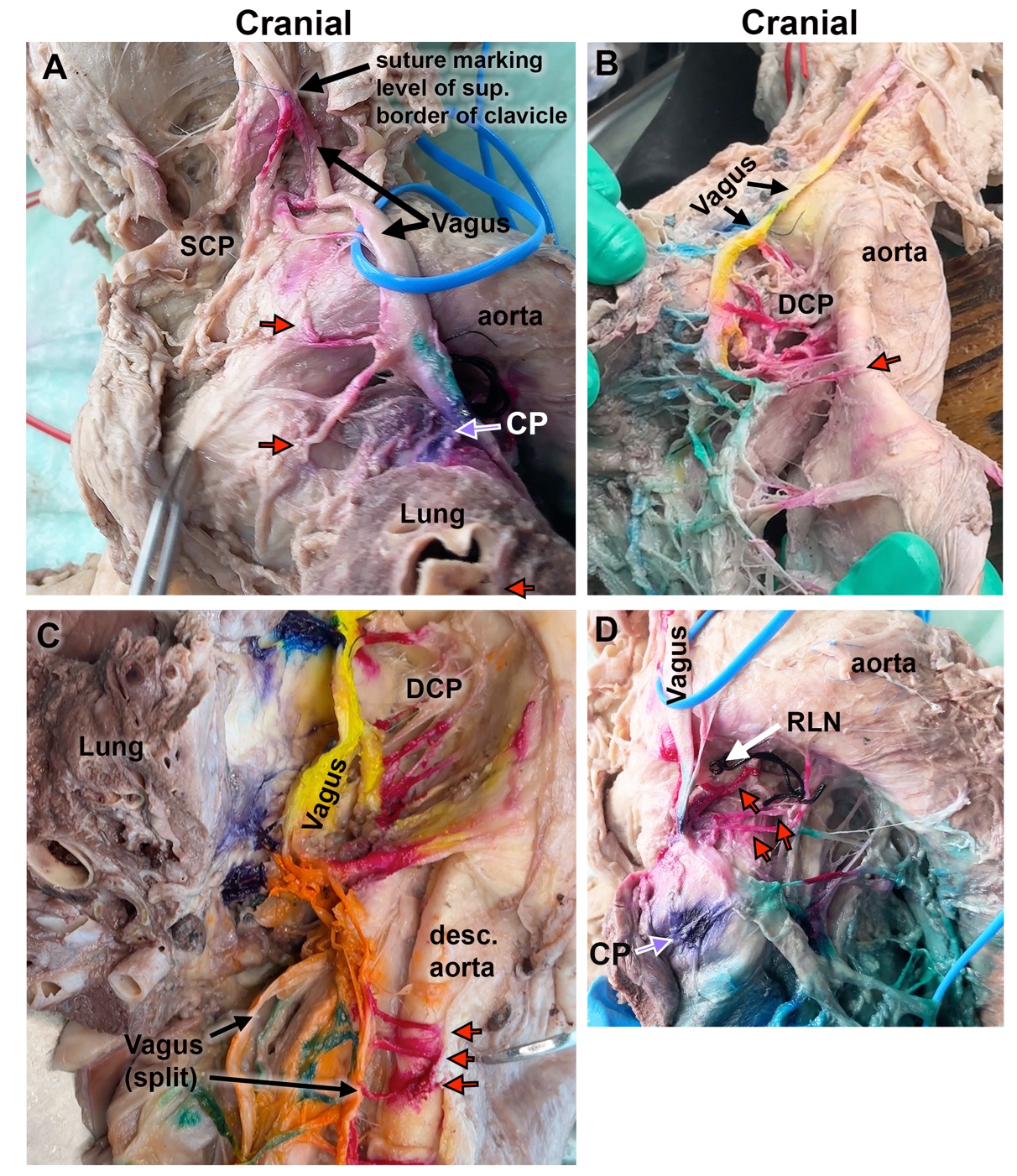

Figure 5. Photographic image of cardiovascular and cardiac branches from the vagus in the upper thoracic region. Top side of each image is cranial end of the image. Aorta and lung (when present in image) are indicated. (A) A blue suture on the vagus nerve demarcates the level of the superior border of the clavicle (vagus is delineated with a blue vessel loop). “SCP” marks the site of cardiac branches to the superior cardiac plexus (cranial to the aorta). Branches from the vagus to the arch of the aorta are delineated with red arrows, while cardiopulmonary (CP) branches are delineated by a purple arrow. (B) Anterior view of vagus (yellow dye) descending across the aorta before giving rise to deep cervical plexus (DCP) branches (dyed red, one indicated with red arrow). Blue-green dyed branches are tracheoesophageal; greened dyed branches are eophageal branches. (C) Another anterior view of the vagus (yellow dye) in a different cadaver giving rise to the deep cervical plexus (DCP) branches (dyed red) as well as the branches to the descending aorta (red arrows). The vagus nerve in this cadaver then split into trunks before descending further into the thorax. (D) Vagus is delineated with a blue vessel loop, before it gives off a recurrent larynageal nerve (RLN, large knotted black suture), several cardiac branches (red arrows and dye), and a cardiopulmonary branch (purple arrows and dye).

**Supplementary Figure 6**

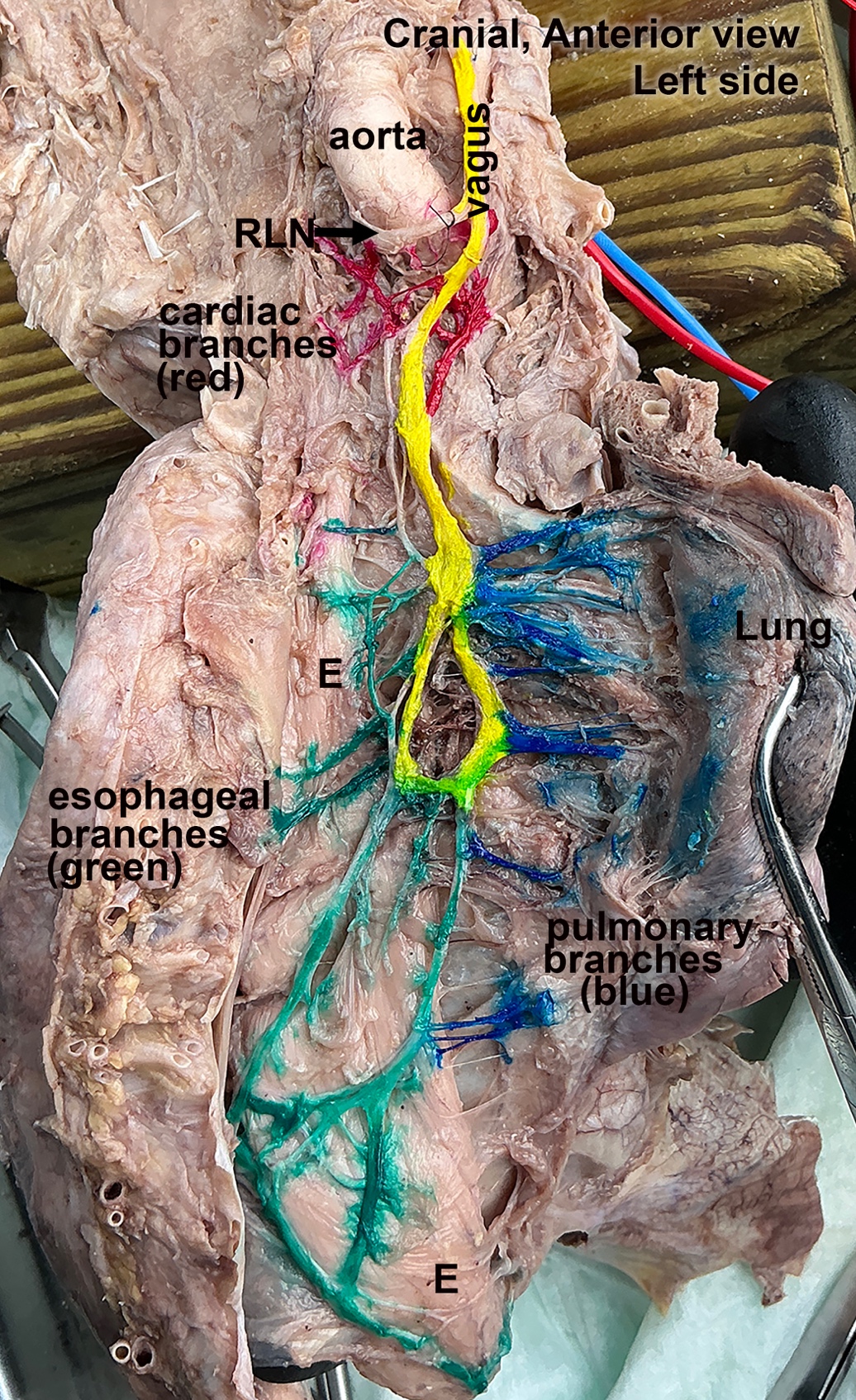

Figure 6. Photographic image of the thoracic vagus. Anterior side of vagus is dyed yellow. It gives off a recurrent laryngeal nerve branch (RLN, small black suture), cardiac branches caudal to the aorta (dyed red), pulmonary branches to the lungs (dyed blue) and many esophageal branches (dyed green) that innervate the esophagus (E) directly (more cranial branches) or indirectly via the eophageal plexus (more caudal branches).

**Supplementary Figure 7.**
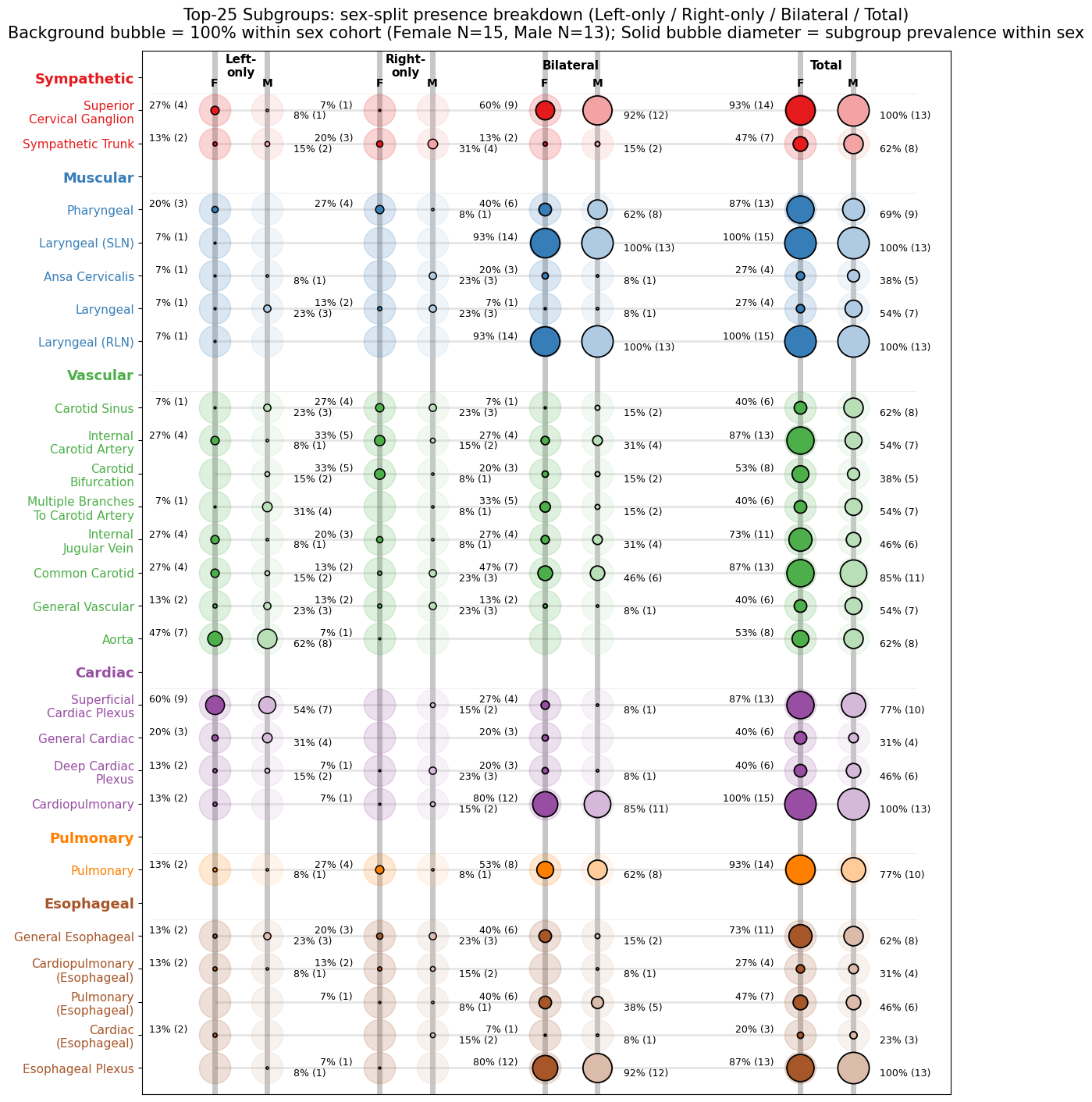

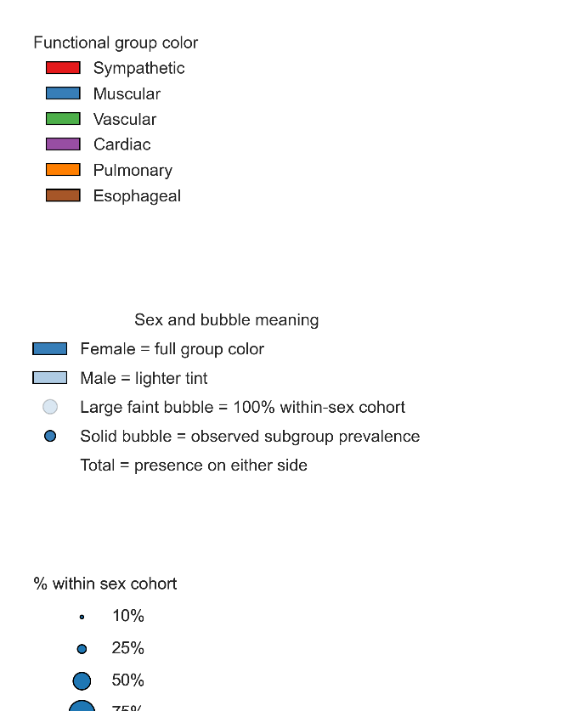

**Supplementary Figure 7. Sex- and side-stratified “subway map” of donor-level branch presence across the vagus nerve (Top 25 subgroups).** This layout summarizes how often common branch subgroups occur unilaterally, bilaterally, or in either side overall, while simultaneously showing sex-specific differences in donor-level presence patterns. The 25 most frequently observed branch subgroups are shown as sex-resolved donor-level presence phenotypes across the registered vagus dataset. Rows are organized beneath functional-group headers and ordered within each group from more proximal to more distal mean emergence position. For each subgroup, four presence classes are displayed: Left-only, Right-only, Bilateral, and Total (present on either side). Within each class, Female is shown first in the full functional-group color and Male second in a lighter tint of the same color. Large faint background circles indicate 100% of the corresponding sex-specific cohort (Female N=15, Male N=13), whereas solid circle diameter scales with subgroup prevalence within that sex. Labels report prevalence as percent of the sex-specific cohort, with donor counts in parentheses.

**Supplementary Figure 8.**
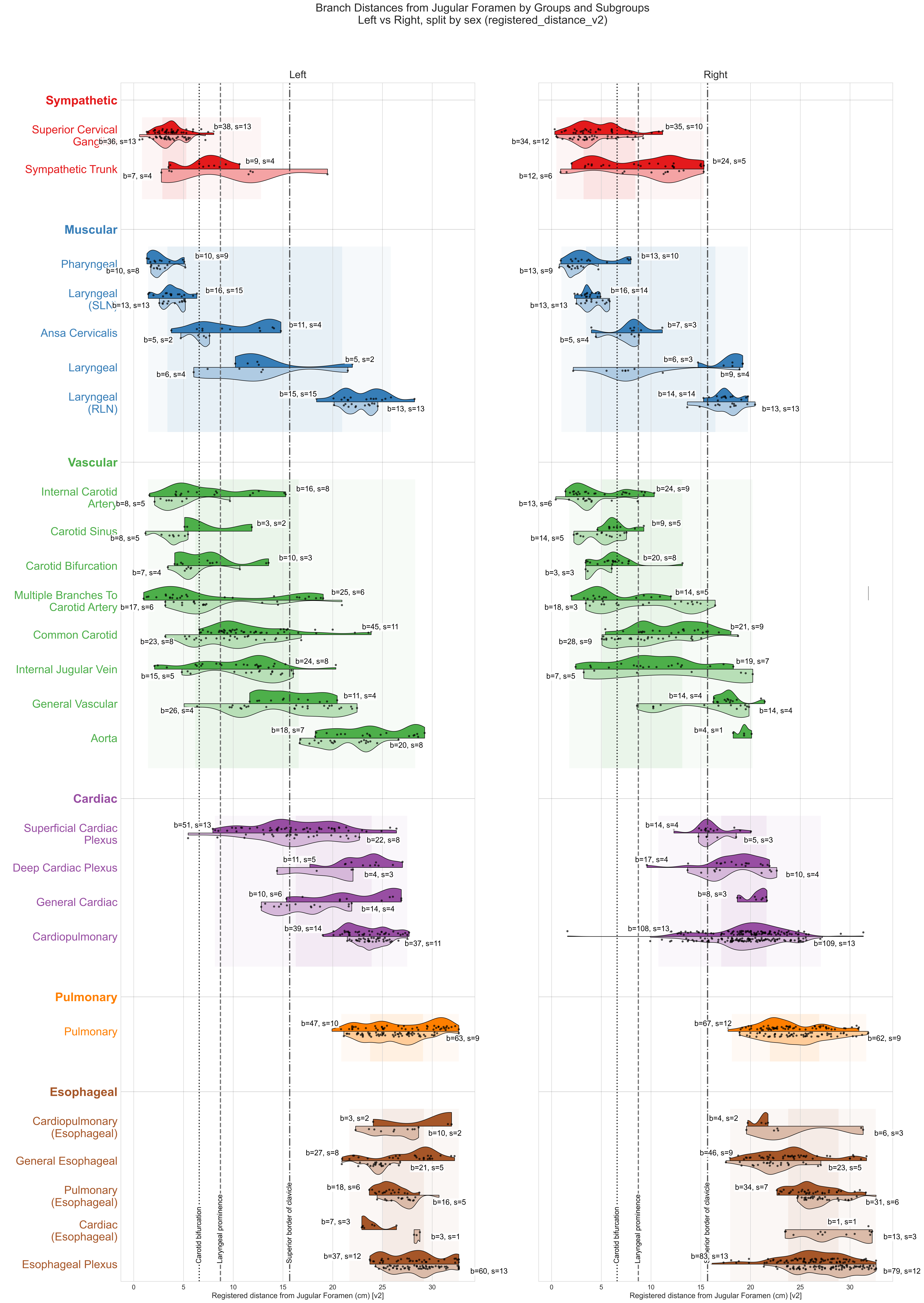

**Supplementary Figure 8. Branch distances on the template vagus nerve by functional group and subgroup, split by sex and side (Top 25).** Subgroups are arranged beneath functional-group header rows and ordered within each group from more proximal to more distal mean registered distance. The 25 most frequently observed branch subgroups are shown as sex-resolved distributions along the registered vagus template. Left and right panels display Left and Right vagus nerves, respectively. Within each subgroup row, split violins show Female branches in the full functional-group color and Male branches in a lighter tint of the same color; black points indicate individual branches. Text annotations report b = number of branches and s = number of contributing subjects for each sex within each side. Vertical reference lines mark the cohort template positions of the carotid bifurcation, laryngeal prominence, and superior border of the clavicle. Light background bands indicate the pooled distribution range and interquartile span of each functional group within each panel.

**Supplementary Figure 9.**
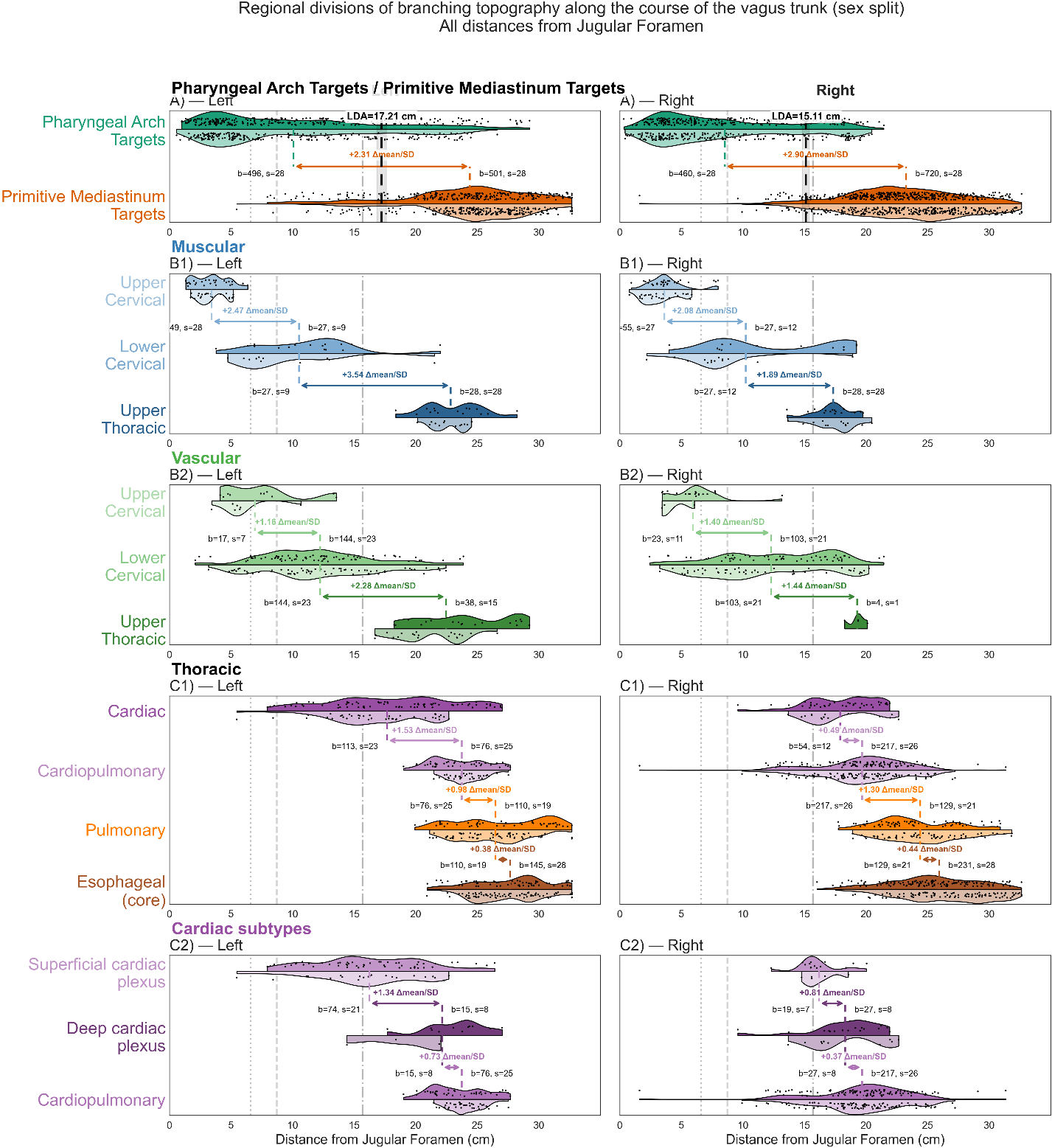

**A1, A2, B1, B2, etc not delineated.**

**Supplementary Figure 9. Sex-resolved regional divisions of branching topography along the human vagus trunk.** Branch emergence locations are plotted as distance (in cm) from the jugular foramen, increasing caudally. Rows show five regional analyses and columns show Left versus Right data. Within each category, split violins display Female distributions in a darker color; Male distributions in a lighter tint of the same color; black jittered points indicate individual branch events. Vertical reference lines mark the carotid bifurcation, laryngeal prominence, and superior border of the clavicle on the registered template. Colored dashed stubs denote pooled within-side category means, and double-headed arrows indicate pooled shifts between adjacent categories, annotated as standardized effect size (Δmean/SD). Text labels report b = number of branch events and s = number of donors contributing to that category within that side. Black dashed vertical line marks the pooled within-side linear discriminant analysis boundary between pharyngeal arch target branches and primitive mediastinum target branches, with the shaded band indicating the bootstrap confidence interval. A) Pharyngeal arch targets versus primitive mediastinum targets. B1) Muscular subdivision into upper cervical, lower cervical, and upper thoracic groups. B2) Vascular subdivision into upper cervical, lower cervical, and upper thoracic groups. C1) Thoracic subdivision into cardiac, cardiopulmonary, pulmonary, and esophageal-core groups. C2) Cardiac subtypes comprising superficial cardiac plexus, deep cardiac plexus, and cardiopulmonary branches. Sex splitting is shown for visualization of subgroup distributions, whereas the overlaid mean, effect-size, and LDA annotations are computed from pooled within-side data.

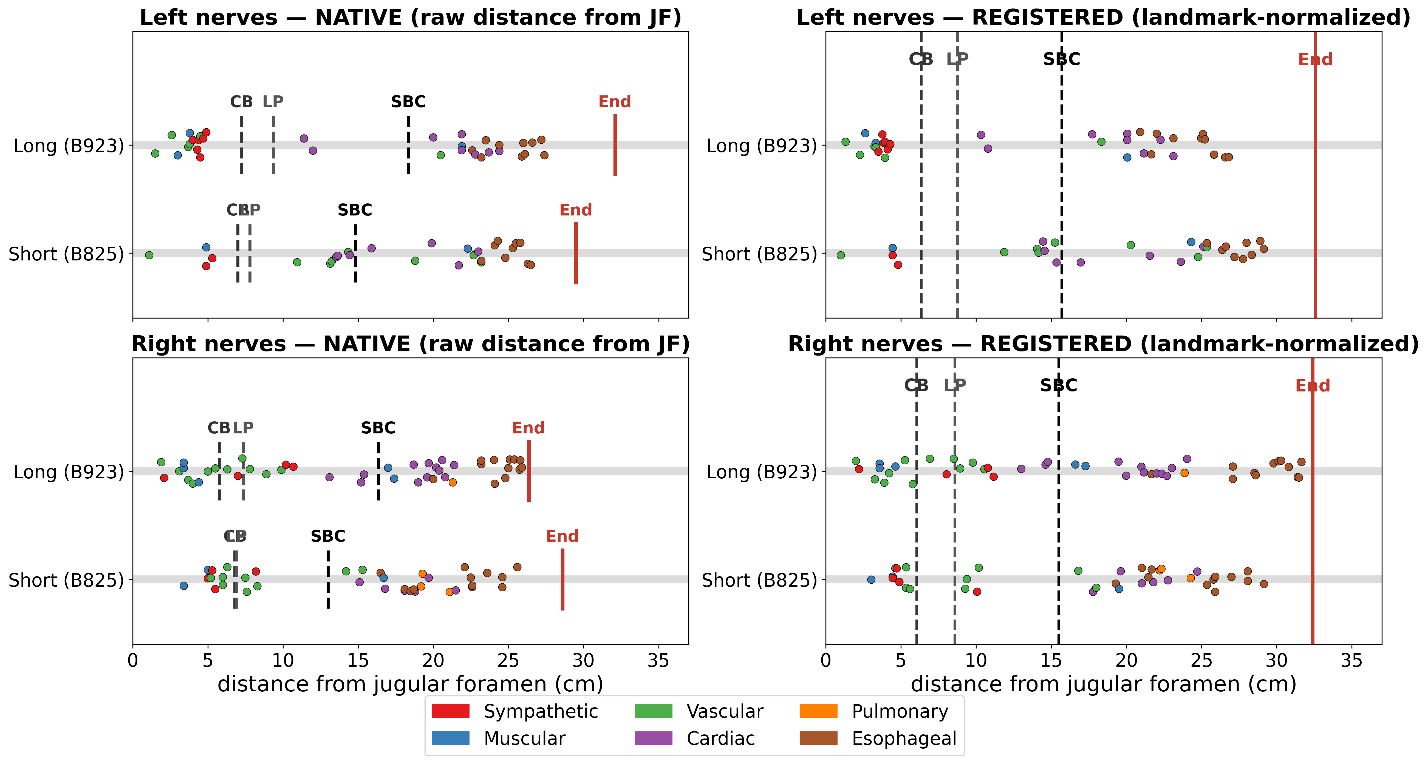

Supplementary Figure 10. Landmark registration aligns branches from nerves of different lengths on a common distance axis. Emergence positions of branches from two short (donor B825, left and right) and two long (donor B923, left and right) vagus nerves are shown in native (actual distance from the jugular foramen) and registered (landmark-normalized) coordinates, for the left (top) and right (bottom) sides. Branches are colored by functional group; dashed lines mark the carotid bifurcation (CB), laryngeal prominence (LP), and superior border of the clavicle (SBC), and the red line marks the nerve end. In native coordinates the landmarks and nerve end of the two donors lie at different absolute positions; after registration they coincide at the cohort-mean template positions, while each branch retains its order and its position relative to the landmarks.

**Supplementary Table S1.** Nerve sides with non-canonical landmark ordering (cm from jugular foramen).

| Donor | Side | Carotid bifurcation (cm) | Laryngeal prominence (cm) | Superior border of clavicle (cm) | Handling |
| --- | --- | --- | --- | --- | --- |
| B788 | R | 6.5 | 5.5 | 14.9 | Inversion |
| B924 | L | 10.1 | 9.0 | 17.6 | Inversion |
| B924 | R | 9.1 | 9.1 | 17.1 | Tie |
| B927 | L | 8.0 | 8.0 | 15.5 | Tie |
| B927 | R | 6.4 | 5.9 | 13.9 | Inversion |

“Inversion” = laryngeal prominence recorded proximal to carotid bifurcation; “Tie” = identical carotid and laryngeal coordinates, requiring ε handling).

| Supplementary Table S2. Sex-stratified summary table of all data of vagal branch emergence by group and subgroup.  This table compiles the full sex-resolved summary of registered branch-emergence distances along the standardized vagus nerve axis. At the start of each group block, a group-level summary row reports the pooled distribution for that entire group. The rows beneath each header then report subgroup-level values separately for Female and Male, with Female listed first. For each row, values are shown for the pooled dataset and separately for the left (L) and right (R) vagus as branch count (n), mean registered distance from the jugular foramen (cm), interquartile range (IQR), and side difference in mean emergence location (Δmean [L–R], cm). Positive Δmean values indicate more distal emergence on the left, whereas negative values indicate more distal emergence on the right. Rows are organized by group and, within each group, ordered proximodistally by subgroup-level mean emergence location. | | | | | | | | | | |
| --- | --- | --- | --- | --- | --- | --- | --- | --- | --- | --- |
| Group and Subgroup (indented) | Total n | Total mean (cm) | Total IQR (cm) | n (L) | Mean (L) (cm) | IQR (L) (cm) | n (R) | Mean (R) (cm) | IQR (R) (cm) | Δmean (L–R) (cm) |
| Sympathetic (Female) | 110 | 5.51 | 3.22–7.33 | 49 | 4.43 | 2.91–5.29 | 61 | 6.37 | 3.39–8.96 | -1.94 |
| Sympathetic (Male) | 89 | 5.00 | 2.77–5.65 | 43 | 4.50 | 2.77–5.12 | 46 | 5.48 | 2.79–7.56 | -0.98 |
| Superior Cervical Ganglion (Female) | 73 | 4.30 | 2.91–5.94 | 38 | 3.87 | 2.88–4.42 | 35 | 4.77 | 3.00–6.05 | -0.90 |
| Superior Cervical Ganglion (Male) | 70 | 3.80 | 2.70–4.82 | 36 | 3.57 | 2.69–4.56 | 34 | 4.05 | 2.71–5.36 | -0.48 |
| Sympathetic Trunk (Female) | 33 | 8.52 | 5.00–12.17 | 9 | 7.21 | 6.59–8.63 | 24 | 9.00 | 4.76–12.28 | -1.79 |
| Sympathetic Trunk (Male) | 19 | 9.44 | 5.48–11.85 | 7 | 9.27 | 3.69–11.85 | 12 | 9.54 | 7.28–11.97 | -0.27 |
| Sympathetic Trunk & Superior Cervical Ganglion (Female) | 4 | 2.69 | 2.40–2.86 | 2 | 2.58 | 2.56–2.60 | 2 | 2.80 | 2.40–3.19 | -0.22 |
| Sympathetic Trunk & Superior Cervical Ganglion (Male) | 0 |  |  | 0 |  |  | 0 |  |  |  |
| Muscular (Female) | 114 | 10.08 | 3.58–17.30 | 57 | 10.76 | 3.69–20.08 | 57 | 9.40 | 3.58–17.10 | 1.37 |
| Muscular (Male) | 100 | 9.02 | 3.11–15.92 | 47 | 10.16 | 3.26–21.20 | 53 | 8.01 | 2.97–13.64 | 2.15 |
| Pharyngeal (Female) | 23 | 3.52 | 2.08–4.41 | 10 | 2.78 | 1.78–3.40 | 13 | 4.10 | 2.42–5.58 | -1.32 |
| Pharyngeal (Male) | 23 | 2.49 | 1.79–2.81 | 10 | 2.71 | 2.11–3.01 | 13 | 2.33 | 1.77–2.64 | 0.38 |
| Laryngeal (SLN) (Female) | 32 | 3.75 | 3.30–4.49 | 16 | 3.80 | 3.15–4.66 | 16 | 3.71 | 3.38–4.14 | 0.09 |
| Laryngeal (SLN) (Male) | 26 | 4.16 | 3.39–5.08 | 13 | 4.01 | 3.02–4.87 | 13 | 4.31 | 3.52–5.38 | -0.31 |
| Ansa Cervicalis (Female) | 18 | 9.30 | 7.16–12.26 | 11 | 10.07 | 6.91–13.34 | 7 | 8.10 | 8.00–8.65 | 1.97 |
| Ansa Cervicalis (Male) | 10 | 6.97 | 6.59–7.94 | 5 | 6.63 | 6.49–7.27 | 5 | 7.31 | 6.90–8.34 | -0.67 |
| General Laryngeal, and any non-recurrent laryngeal branches (Female) | 12 | 15.75 | 12.54–18.44 | 5 | 13.73 | 11.46–12.57 | 7 | 17.20 | 16.01–18.70 | -3.47 |
| General Laryngeal and any non-recurrent laryngeal branches (Male) | 15 | 9.82 | 7.28–12.01 | 6 | 11.90 | 8.23–12.94 | 9 | 8.43 | 7.14–8.34 | 3.48 |
| Laryngeal (RLN) (Female) | 29 | 20.39 | 17.47–23.40 | 15 | 23.03 | 21.34–24.67 | 14 | 17.56 | 16.81–18.14 | 5.47 |
| Laryngeal (RLN) (Male) | 26 | 19.99 | 18.03–22.33 | 13 | 22.61 | 21.60–23.71 | 13 | 17.38 | 16.48–18.36 | 5.23 |
| Vascular (Female) | 277 | 10.83 | 5.79–15.13 | 152 | 12.06 | 6.88–15.71 | 125 | 9.34 | 5.34–13.16 | 2.72 |
| Vascular (Male) | 222 | 10.53 | 5.28–14.97 | 125 | 11.75 | 5.71–16.84 | 97 | 8.96 | 5.03–13.03 | 2.78 |
| Internal Carotid Artery (Female) | 40 | 6.18 | 3.23–8.29 | 16 | 7.94 | 4.59–11.77 | 24 | 5.01 | 2.55–7.43 | 2.93 |
| Internal Carotid Artery (Male) | 21 | 3.99 | 3.19–4.04 | 8 | 4.41 | 3.14–4.98 | 13 | 3.73 | 3.19–4.01 | 0.68 |
| Carotid Sinus (Female) | 12 | 6.82 | 5.69–7.29 | 3 | 7.46 | 5.25–8.63 | 9 | 6.61 | 5.81–6.93 | 0.85 |
| Carotid Sinus (Male) | 22 | 4.45 | 2.99–5.51 | 8 | 3.83 | 3.40–4.67 | 14 | 4.81 | 2.99–6.00 | -0.98 |
| External Carotid Artery (Female) | 0 |  |  | 0 |  |  | 0 |  |  |  |
| External Carotid Artery (Male) | 1 | 6.03 | 6.03–6.03 | 1 | 6.03 | 6.03–6.03 | 0 |  |  |  |
| Carotid Bifurcation (Female) | 30 | 6.66 | 4.80–7.71 | 10 | 7.64 | 4.80–8.23 | 20 | 6.17 | 5.16–6.84 | 1.47 |
| Carotid Bifurcation (Male) | 10 | 5.46 | 4.01–5.76 | 7 | 5.93 | 5.22–5.74 | 3 | 4.37 | 3.53–4.84 | 1.56 |
| Multiple Branches to Carotid Artery (Female) | 39 | 7.65 | 3.65–10.47 | 25 | 8.38 | 3.65–16.42 | 14 | 6.34 | 4.25–9.09 | 2.04 |
| Multiple Branches to Carotid Artery (Male) | 35 | 8.84 | 5.10–13.36 | 17 | 8.04 | 4.87–7.31 | 18 | 9.60 | 5.20–14.01 | -1.56 |
| Internal Jugular Vein (Female) | 43 | 10.18 | 6.96–12.73 | 24 | 10.21 | 6.88–12.65 | 19 | 10.14 | 7.64–12.85 | 0.07 |
| Internal Jugular Vein (Male) | 22 | 12.13 | 8.97–14.89 | 15 | 11.85 | 10.71–14.36 | 7 | 12.71 | 6.44–19.28 | -0.86 |
| Common Carotid (Female) | 66 | 12.13 | 9.09–14.76 | 45 | 12.14 | 9.07–14.77 | 21 | 12.10 | 9.16–14.73 | 0.04 |
| Common Carotid (Male) | 51 | 9.52 | 6.54–11.67 | 23 | 9.71 | 6.53–11.91 | 28 | 9.36 | 6.63–11.36 | 0.35 |
| General Vascular (Female) | 25 | 17.00 | 16.27–18.22 | 11 | 15.75 | 13.21–18.11 | 14 | 17.98 | 16.89–18.17 | -2.23 |
| General Vascular (Male) | 40 | 15.32 | 10.81–19.53 | 26 | 15.25 | 10.91–19.63 | 14 | 15.47 | 10.41–19.35 | -0.22 |
| Aorta (Female) | 22 | 23.19 | 19.57–27.11 | 18 | 24.06 | 20.93–28.02 | 4 | 19.27 | 19.08–19.54 | 4.79 |
| Aorta (Male) | 20 | 21.03 | 18.78–23.35 | 20 | 21.03 | 18.78–23.35 | 0 |  |  |  |
| Cardiac (Female) | 259 | 19.62 | 17.04–22.45 | 112 | 20.02 | 16.17–24.04 | 147 | 19.32 | 17.11–21.54 | 0.70 |
| Cardiac (Male) | 201 | 19.68 | 16.67–22.68 | 77 | 20.26 | 16.27–23.70 | 124 | 19.32 | 16.77–21.77 | 0.95 |
| Superficial Cardiac Plexus (Female) | 65 | 16.33 | 14.36–18.96 | 51 | 16.37 | 13.71–19.91 | 14 | 16.16 | 15.44–16.57 | 0.21 |
| Superficial Cardiac Plexus (Male) | 27 | 16.17 | 14.40–18.50 | 22 | 16.16 | 13.95–19.51 | 5 | 16.23 | 15.58–16.67 | -0.08 |
| Deep Cardiac Plexus (Female) | 28 | 20.24 | 18.00–22.19 | 11 | 23.20 | 21.59–24.48 | 17 | 18.33 | 17.06–19.83 | 4.88 |
| Deep Cardiac Plexus (Male) | 14 | 18.54 | 16.38–21.94 | 4 | 19.18 | 17.40–21.93 | 10 | 18.28 | 16.38–21.74 | 0.90 |
| General Cardiac (Female) | 18 | 21.66 | 20.47–24.11 | 10 | 22.57 | 20.72–25.98 | 8 | 20.53 | 20.07–21.31 | 2.04 |
| General Cardiac (Male) | 14 | 17.07 | 14.66–18.59 | 14 | 17.07 | 14.66–18.59 | 0 |  |  |  |
| Cardiopulmonary (Female) | 147 | 20.77 | 19.05–23.12 | 39 | 23.46 | 21.62–25.22 | 108 | 19.79 | 18.20–22.02 | 3.67 |
| Cardiopulmonary (Male) | 146 | 20.69 | 18.19–23.67 | 37 | 24.03 | 23.10–25.09 | 109 | 19.55 | 17.68–21.88 | 4.48 |
| Cardiac Plexus (Female) | 1 | 11.33 | 11.33–11.33 | 1 | 11.33 | 11.33–11.33 | 0 |  |  |  |
| Cardiac Plexus (Male) | 0 |  |  | 0 |  |  | 0 |  |  |  |
| Pulmonary (Female) – lateral and medial pulmonary branches combined | 114 | 25.19 | 22.14–28.11 | 47 | 27.03 | 24.31–30.64 | 67 | 23.90 | 21.79–26.54 | 3.13 |
| Pulmonary (Male) – lateral and medial pulmonary branches combined | 125 | 25.51 | 23.03–27.72 | 63 | 26.07 | 23.73–27.95 | 62 | 24.94 | 22.45–27.19 | 1.14 |
| Esophageal (Female) | 260 | 26.02 | 23.86–28.69 | 92 | 27.14 | 24.86–29.37 | 168 | 25.41 | 23.24–27.92 | 1.73 |
| Esophageal (Male) | 262 | 27.12 | 24.93–29.40 | 110 | 27.37 | 25.32–29.08 | 152 | 26.94 | 24.70–29.66 | 0.42 |
| Cardiopulmonary (from Esophageal Plexus) (Female) | 7 | 24.53 | 21.32–27.83 | 3 | 29.20 | 27.83–31.77 | 4 | 21.03 | 20.76–21.59 | 8.17 |
| Cardiopulmonary (from Esophageal Plexus) (Male) | 16 | 25.60 | 22.76–28.08 | 10 | 25.98 | 24.36–27.90 | 6 | 24.98 | 22.45–29.04 | 0.99 |
| General Esophageal (Female) | 73 | 25.30 | 22.15–28.73 | 27 | 27.38 | 25.88–29.66 | 46 | 24.08 | 21.71–25.51 | 3.30 |
| General Esophageal (Male) | 44 | 23.35 | 21.94–24.87 | 21 | 24.42 | 24.08–24.93 | 23 | 22.38 | 21.04–23.78 | 2.04 |
| Pulmonary (from Esophageal Plexus) (Female) | 52 | 26.04 | 24.59–27.23 | 18 | 25.69 | 24.76–26.53 | 34 | 26.23 | 24.40–27.78 | -0.54 |
| Pulmonary (from Esophageal Plexus) (Male) | 47 | 27.21 | 25.86–28.00 | 16 | 27.15 | 26.44–27.87 | 31 | 27.24 | 25.72–28.34 | -0.09 |
| Cardiac (from Esophageal Plexus) (Female) | 8 | 24.82 | 23.03–24.76 | 7 | 23.81 | 23.01–24.09 | 1 | 31.85 | 31.85–31.85 | -8.04 |
| Cardiac (from Esophageal Plexus) (Male) | 16 | 28.36 | 26.98–30.85 | 3 | 28.46 | 28.31–28.60 | 13 | 28.34 | 26.79–31.97 | 0.12 |
| Esophageal Plexus (Female) | 120 | 26.62 | 24.72–28.99 | 37 | 28.13 | 25.07–29.73 | 83 | 25.95 | 23.94–28.53 | 2.18 |
| Esophageal Plexus (Male) | 139 | 28.31 | 26.39–30.37 | 60 | 28.63 | 26.53–30.14 | 79 | 28.07 | 25.55–30.43 | 0.55 |
| Multiple Targets (Female) | 20 | 9.12 | 3.05–14.84 | 11 | 8.85 | 2.55–16.19 | 9 | 9.45 | 4.44–14.83 | -0.60 |
| Multiple Targets (Male) | 24 | 12.68 | 7.50–18.44 | 12 | 8.37 | 4.65–12.19 | 12 | 16.98 | 17.58–18.55 | -8.61 |
| Vascular And Pharyngeal (Female) | 1 | 2.10 | 2.10–2.10 | 0 |  |  | 1 | 2.10 | 2.10–2.10 |  |
| Vascular And Pharyngeal (Male) | 0 |  |  | 0 |  |  | 0 |  |  |  |
| Vascular And Sympathetic Chain (Female) | 4 | 4.24 | 3.92–4.68 | 2 | 4.35 | 4.26–4.44 | 2 | 4.13 | 3.64–4.63 | 0.22 |
| Vascular And Sympathetic Chain (Male) | 5 | 5.08 | 2.80–5.81 | 5 | 5.08 | 2.80–5.81 | 0 |  |  |  |
| Sympathetic Chain, Cardiovascular, And Laryngeal Muscles (Female) | 0 |  |  | 0 |  |  | 0 |  |  |  |
| Sympathetic Chain, Cardiovascular, & Laryngeal Muscles (Male) | 2 | 7.72 | 7.69–7.76 | 2 | 7.72 | 7.69–7.76 | 0 |  |  |  |
| Sympathetic Chain, Cardiovascular, & Pulmonary (Female) | 11 | 7.78 | 2.55–14.15 | 5 | 2.19 | 1.75–2.94 | 6 | 12.44 | 12.03–14.87 | -10.25 |
| Sympathetic Chain, Cardiovascular, & Pulmonary (Male) | 6 | 11.85 | 8.26–16.34 | 3 | 10.09 | 6.78–14.33 | 3 | 13.61 | 11.14–16.90 | -3.52 |
| Vascular & Laryngeal (Female) | 1 | 11.66 | 11.66–11.66 | 1 | 11.66 | 11.66–11.66 | 0 |  |  |  |
| Vascular & Laryngeal (Male) | 1 | 16.51 | 16.51–16.51 | 1 | 16.51 | 16.51–16.51 | 0 |  |  |  |
| Vascular & Muscular - Laryngeal (Female) | 0 |  |  | 0 |  |  | 0 |  |  |  |
| Vascular & Muscular - Laryngeal (Male) | 1 | 14.98 | 14.98–14.98 | 0 |  |  | 1 | 14.98 | 14.98–14.98 |  |
| Vascular & Superficial Cardiac Plexus (Female) | 3 | 22.00 | 20.84–22.63 | 3 | 22.00 | 20.84–22.63 | 0 |  |  |  |
| Vascular &Superficial Cardiac Plexus (Male) | 9 | 17.87 | 18.36–18.55 | 1 | 12.84 | 12.84–12.84 | 8 | 18.50 | 18.41–18.57 | -5.65 |

**Supplementary Table S3.** Donor-level laterality and bilaterality of the 25 most frequently observed vagal branch subgroups [% of donors (donor count)]

Rows list the Top-25 subgroups (selected by branch frequency) and are ordered proximodistally within each group by the total (pooled by side) mean registered distance on the v2 axis (cm). Presence is defined at the donor level (N=28): a subgroup is counted as present on a side if observed at least once on that donor-side; Left-only and Right-only indicate unilateral presence. Bilateral indicates presence on both sides within the same donor. Total indicates presence on either side. Values are reported as percent of donors with donor counts in parentheses.

| Group | Subgroup | Mean distance (cm) | Left-only presence  % (donor ct) | Right-only presence  % (donor ct) | Bilateral presence  % (donor ct) | Total presence  % (donor ct) |
| --- | --- | --- | --- | --- | --- | --- |
| Sympathetic | Superior Cervical Ganglion | 4.06 | 18% (5) | 4% (1) | 75% (21) | 96% (27) |
| Sympathetic | Sympathetic trunk | 8.85 | 14% (4) | 25% (7) | 14% (4) | 54% (15) |
| Muscular | Pharyngeal | 3.01 | 11% (3) | 18% (5) | 50% (14) | 79% (22) |
| Muscular | Laryngeal (SLN) | 3.94 | 4% (1) | 0% (0) | 96% (27) | 100% (28) |
| Muscular | Ansa Cervicalis | 8.47 | 7% (2) | 11% (3) | 14% (4) | 32% (9) |
| Muscular | General Laryngeal | 12.41 | 14% (4) | 18% (5) | 7% (2) | 39% (11) |
| Muscular | Laryngeal (RLN) | 20.20 | 4% (1) | 0% (0) | 96% (27) | 100% (28) |
| Vascular | Internal Carotid Artery | 5.43 | 18% (5) | 25% (7) | 29% (8) | 71% (20) |
| Vascular | Carotid sinus | 5.29 | 14% (4) | 25% (7) | 11% (3) | 50% (14) |
| Vascular | Carotid Bifurcation | 6.36 | 7% (2) | 21% (6) | 18% (5) | 46% (13) |
| Vascular | Multiple branches to carotid artery | 8.21 | 18% (5) | 4% (1) | 25% (7) | 46% (13) |
| Vascular | Common Carotid | 10.99 | 21% (6) | 18% (5) | 46% (13) | 86% (24) |
| Vascular | Internal Jugular Vein | 10.84 | 18% (5) | 14% (4) | 29% (8) | 61% (17) |
| Vascular | General Vascular | 15.97 | 18% (5) | 18% (5) | 11% (3) | 46% (13) |
| Vascular | Aorta | 22.16 | 54% (15) | 4% (1) | 0% (0) | 57% (16) |
| Cardiac | Superficial Cardiac Plexus | 16.28 | 57% (16) | 7% (2) | 18% (5) | 82% (23) |
| Cardiac | Deep Cardiac Plexus | 19.67 | 14% (4) | 14% (4) | 14% (4) | 43% (12) |
| Cardiac | General Cardiac | 19.65 | 25% (7) | 0% (0) | 11% (3) | 36% (10) |
| Cardiac | Cardiopulmonary | 20.79 | 7% (2) | 11% (3) | 82% (23) | 100% (28) |
| Pulmonary | Pulmonary (medial and lateral combined) | 25.36 | 11% (3) | 18% (5) | 57% (16) | 86% (24) |
| Esophageal | Cardiopulmonary (Esophageal) | 25.28 | 11% (3) | 14% (4) | 4% (1) | 29% (8) |
| Esophageal | General Esophageal | 24.57 | 18% (5) | 21% (6) | 29% (8) | 68% (19) |
| Esophageal | Pulmonary (Esophageal) | 26.60 | 0% (0) | 7% (2) | 39% (11) | 46% (13) |
| Esophageal | Cardiac (Esophageal) | 27.18 | 7% (2) | 7% (2) | 7% (2) | 21% (6) |
| Esophageal | Esophageal Plexus | 27.53 | 4% (1) | 4% (1) | 86% (24) | 93% (26) |

**Supplemental Table S4.** Branch location by side (total (T), left (L) and right (R)) organized by group and subgroup.

Values are reported as n (number of branches), mean branching location (cm), and range (min-max). Δmean (L-R) is Mean(L) - Mean(R) (positive values indicate more distal branching on the left; negative values indicate more distal branching on the right). Distances are defined on the registered longitudinal template (Methods).

| Group | Subgroup | n (T) | Mean (T) | Range (T) | n (L) | Mean (L) | Range (L) | n (R) | Mean (R) | Range (R) | Δmean (L–R) |
| --- | --- | --- | --- | --- | --- | --- | --- | --- | --- | --- | --- |
| Sympathetic | Superior Cervical Ganglion | 143 | 4.06 | 0.32–11.17 | 74 | 3.73 | 0.58–8.05 | 69 | 4.41 | 0.32–11.17 | -0.69 |
| Sympathetic | Sympathetic Trunk | 52 | 8.85 | 0.91–19.48 | 16 | 8.11 | 2.78–19.48 | 36 | 9.18 | 0.91–15.32 | -1.07 |
| Sympathetic | Sympathetic Trunk & Superior Cervical Ganglion | 4 | 2.69 | 2.00–3.59 | 2 | 2.58 | 2.54–2.62 | 2 | 2.8 | 2.00–3.59 | -0.22 |
| Muscular | Pharyngeal | 46 | 3.01 | 0.77–7.97 | 20 | 2.74 | 1.32–5.18 | 26 | 3.21 | 0.77–7.97 | -0.47 |
| Muscular | Laryngeal (SLN) | 58 | 3.94 | 1.45–6.35 | 29 | 3.89 | 1.45–6.35 | 29 | 3.98 | 2.31–5.84 | -0.09 |
| Muscular | Ansa Cervicalis | 28 | 8.47 | 3.78–14.79 | 16 | 8.99 | 3.78–14.79 | 12 | 7.77 | 4.00–11.13 | +1.22 |
| Muscular | General Laryngeal | 26 | 12.59 | 5.66–21.19 | 11 | 11.92 | 9.82–12.84 | 15 | 13.08 | 5.66–21.19 | -1.16 |
| Muscular | Laryngeal (RLN) | 55 | 19.02 | 8.89–29.08 | 28 | 22.05 | 19.78–29.08 | 27 | 15.87 | 8.89–19.32 | +6.18 |
| Vascular | Internal Carotid Artery | 61 | 5.75 | 1.58–16.04 | 24 | 6.98 | 2.16–16.04 | 37 | 4.95 | 1.58–14.22 | +2.03 |
| Vascular | External Carotid Artery | 1 | 6.03 | 6.03–6.03 | 1 | 6.03 | 6.03–6.03 | 0 |  |  |  |
| Vascular | Carotid Sinus | 34 | 6.18 | 2.31–19.48 | 11 | 5.03 | 3.63–8.91 | 23 | 6.73 | 2.31–19.48 | -1.70 |
| Vascular | Carotid Bifurcation | 40 | 6.32 | 0.58–17.58 | 17 | 6.96 | 2.31–17.58 | 23 | 5.84 | 0.58–11.65 | +1.12 |
| Vascular | Multiple Branches To Carotid Artery | 74 | 8.45 | 0.91–19.48 | 42 | 8.54 | 0.91–19.48 | 32 | 8.34 | 2.47–15.11 | +0.20 |
| Vascular | Common Carotid | 117 | 11.58 | 3.78–23.34 | 68 | 11.66 | 3.78–23.34 | 49 | 11.47 | 4.00–22.52 | +0.19 |
| Vascular | Internal Jugular Vein | 65 | 11.75 | 3.78–23.34 | 39 | 11.64 | 3.78–23.34 | 26 | 11.92 | 4.00–22.52 | -0.28 |
| Vascular | General Vascular | 65 | 15.82 | 6.35–29.08 | 37 | 15.05 | 6.35–29.08 | 28 | 16.86 | 15.46–22.52 | -1.81 |
| Vascular | Aorta | 42 | 21.25 | 9.82–29.08 | 38 | 21.44 | 9.82–29.08 | 4 | 19.49 | 18.72–20.53 | +1.95 |
| Cardiac | Cardiac Plexus | 1 | 11.33 | 11.33–11.33 | 1 | 11.33 | 11.33–11.33 | 0 |  |  |  |
| Cardiac | Superficial Cardiac Plexus | 92 | 16.19 | 9.82–24.35 | 65 | 15.93 | 9.82–24.35 | 27 | 16.82 | 13.06–19.32 | -0.89 |
| Cardiac | Deep Cardiac Plexus | 42 | 19.94 | 12.84–29.08 | 13 | 22.02 | 19.78–29.08 | 29 | 19.00 | 12.84–23.34 | +3.02 |
| Cardiac | General Cardiac | 32 | 18.98 | 9.82–29.08 | 18 | 18.41 | 9.82–29.08 | 14 | 19.72 | 18.72–20.53 | -1.31 |
| Cardiac | Cardiopulmonary | 292 | 20.00 | 8.89–29.08 | 76 | 22.46 | 19.78–29.08 | 216 | 19.14 | 8.89–24.35 | +3.32 |
| Pulmonary | Pulmonary | 239 | 24.47 | 16.04–32.68 | 110 | 26.48 | 19.93–32.68 | 129 | 22.76 | 16.04–31.84 | +3.72 |
| Esophageal | Cardiopulmonary (Esophageal) | 23 | 23.58 | 17.73–32.68 | 13 | 25.33 | 23.24–32.68 | 10 | 21.07 | 17.73–23.34 | +4.26 |
| Esophageal | General Esophageal | 117 | 24.02 | 16.04–32.68 | 56 | 23.90 | 20.53–32.68 | 61 | 24.12 | 16.04–32.62 | -0.22 |
| Esophageal | Pulmonary (Esophageal) | 99 | 25.18 | 16.04–32.68 | 20 | 26.73 | 25.78–32.68 | 79 | 24.79 | 16.04–31.84 | +1.94 |
| Esophageal | Cardiac (Esophageal) | 24 | 25.55 | 16.04–32.68 | 5 | 24.81 | 24.57–25.28 | 19 | 25.74 | 16.04–32.62 | -0.93 |
| Esophageal | Esophageal Plexus | 259 | 25.84 | 16.04–32.68 | 108 | 26.15 | 19.93–32.68 | 151 | 25.62 | 16.04–32.62 | +0.53 |

**Supplementary Table S5.** Pulmonary and pulmonary-derived branching location by source (total (T), left (L) and right (R)). Extracted from Supplemental Table 4.

| Subgroup | n (T) | Mean (T), cm | Range (T), cm | n (L) | Mean (L), cm | Range (L), cm | n (R) | Mean (R), cm | Range (R), cm | Δmean (L–R), cm |
| --- | --- | --- | --- | --- | --- | --- | --- | --- | --- | --- |
| Pooled Pulmonary-derived | 654 | 23.47 | 1.66–32.68 | 233 | 25.58 | 19.01–32.68 | 421 | 22.3 | 1.66–32.62 | 3.28 |
| Pulmonary (Pulmonary group, lateral and medial) | 239 | 25.36 | 17.73–32.68 | 110 | 26.48 | 19.93–32.68 | 129 | 24.4 | 17.73–31.84 | 2.08 |
| Cardiopulmonary (Cardiac group) | 293 | 20.73 | 1.66–31.34 | 76 | 23.74 | 19.01–27.68 | 217 | 19.67 | 1.66–31.34 | 4.07 |
| Pulmonary (from Esophageal Plexus) | 99 | 26.6 | 22.65–32.62 | 34 | 26.38 | 23.68–30.69 | 65 | 26.71 | 22.65–32.62 | -0.33 |
| Cardiopulmonary (from Esophageal Plexus) | 23 | 25.28 | 19.59–31.95 | 13 | 26.72 | 22.30–31.95 | 10 | 23.4 | 19.59–31.32 | 3.32 |

**Supplementary Table S6.** Branchless intervals around landmarks, with “carotid” indicating the carotid bifurcation, “larynx” indicating the laryngeal prominence, and “clavicle” indicating the superior border of the clavicle.

Linear mixed-effects models of registered branch-emergence distance from the jugular foramen (cm), per branch group. Fixed effects: sex, side, subgroup, with two-way interactions. Random effect: donor (random intercept). The Pulmonary group is included as a distinct subgroup within the Cardiac group. Reference levels shown as beta=0.00. Significant p (<0.05) in bold.

| Donor | Side | carotid_up_cm | carotid_down_cm | larynx_up_cm | larynx_down_cm | clavicle_up_cm | clavicle_down_cm |
| --- | --- | --- | --- | --- | --- | --- | --- |
| *2305* | Left | 3.68 | 0.39 | 0.64 | 1.57 | 0.60 | 2.56 |
| *2305* | Right | 1.39 | 0.16 | 1.97 | 0.01 | 0.45 | 0.59 |
| *2316* | Left | 3.65 | 1.36 | 0.22 | 0.67 | 0.83 | 0.08 |
| *2316* | Right | 0.49 | 1.19 | 0.94 | 4.44 | 0.96 | 0.37 |
| *B624* | Left | 0.10 | 0.18 | 1.46 | 1.39 | 0.33 | 0.09 |
| *B624* | Right | 0.22 | 0.14 | 1.99 | 5.36 | 1.60 | 0.99 |
| *B733* | Left | 1.28 | 1.82 | 0.31 | 2.82 | 0.22 | 0.47 |
| *B733* | Right | 0.54 | 2.06 | 0.07 | 1.32 | 4.01 | 0.68 |
| *B763* | Left | 0.88 | 6.33 | 3.01 | 4.19 | 1.46 | 2.17 |
| *B763* | Right | 0.54 | 0.54 | 0.76 | 2.46 | 0.09 | 1.21 |
| *B788* | Left | 0.88 | 12.89 | 3.01 | 10.75 | 9.97 | 3.79 |
| *B788* | Right | 3.07 | 2.08 | 0.05 | 5.38 | 0.10 | 0.05 |
| *B791* | Left | 0.78 | 0.84 | 1.11 | 1.95 | 5.01 | 4.22 |
| *B791* | Right | 1.06 | 0.56 | 0.30 | 2.37 | 3.63 | 2.83 |
| *B796* | Left | 0.63 | 0.06 | 1.96 | 2.11 | 4.85 | 2.56 |
| *B796* | Right | 2.72 | 0.60 | 1.08 | 3.32 | 2.48 | 0.58 |
| *B815* | Left | 0.46 | 5.14 | 2.59 | 3.01 | 3.95 | 0.72 |
| *B815* | Right | 1.45 | 1.45 | 0.68 | 0.07 | 0.43 | 0.46 |
| *B822* | Left | 0.21 | 1.07 | 0.94 | 0.92 | 0.28 | 0.47 |
| *B822* | Right | 0.10 | 1.42 | 0.71 | 0.48 | 0.02 | 0.13 |
| *B824* | Left | 0.24 | 1.68 | 0.46 | 0.43 | 6.28 | 2.69 |
| *B824* | Right | 1.47 | 2.99 | 3.60 | 0.86 | 0.18 | 0.14 |
| *B825* | Left | 1.79 | 5.28 | 3.92 | 3.15 | 0.33 | 1.29 |
| *B825* | Right | 0.98 | 2.68 | 3.12 | 0.55 | 5.50 | 1.14 |
| *B848* | Left | 0.38 | 0.32 | 1.82 | 0.16 | 1.24 | 8.16 |
| *B848* | Right | 0.18 | 1.13 | 1.00 | 0.36 | 0.18 | 1.85 |
| *B922* | Left | 0.24 | 2.56 | 2.38 | 0.42 | 1.32 | 0.73 |
| *B922* | Right | 0.33 | 0.94 | 1.19 | 1.31 | 0.85 | 3.40 |
| *B923* | Left | 2.30 | 3.74 | 4.43 | 1.61 | 4.89 | 2.05 |
| *B923* | Right | 0.80 | 0.33 | 0.21 | 0.21 | 0.91 | 0.93 |
| *B924* | Left | 1.30 | 9.91 | 3.44 | 7.78 | 10.39 | 0.82 |
| *B924* | Right | 0.74 | 7.08 | 2.87 | 4.95 | 0.70 | 0.77 |
| *B925* | Left | 1.21 | 4.73 | 3.35 | 2.60 | 0.61 | 0.13 |
| *B925* | Right | 0.83 | 0.41 | 0.13 | 0.55 | 0.41 | 0.33 |
| *B927* | Left | 1.20 | 3.68 | 3.33 | 1.54 | 1.58 | 3.02 |
| *B927* | Right | 2.28 | 2.37 | 4.42 | 0.24 | 0.36 | 1.04 |
| *B930* | Left | 0.56 | 0.39 | 1.75 | 0.02 | 0.29 | 4.08 |
| *B930* | Right | 0.54 | 0.10 | 0.13 | 0.61 | 0.33 | 0.41 |
| *B935* | Left | 0.53 | 0.72 | 1.42 | 0.35 | 1.75 | 0.51 |
| *B935* | Right | 0.54 | 0.13 | 2.01 | 0.28 | 0.64 | 2.66 |
| *B956* | Left | 0.31 | 0.61 | 1.52 | 0.91 | 1.37 | 0.19 |
| *B956* | Right | 1.08 | 0.31 | 0.13 | 2.96 | 0.38 | 1.46 |
| *C70* | Left | 1.13 | 5.38 | 3.26 | 3.25 | 2.71 | 1.01 |
| *C70* | Right | 4.09 | 0.34 | 1.71 | 1.20 | 0.58 | 0.09 |
| *C78* | Left | 0.56 | 0.46 | 0.40 | 1.00 | 1.18 | 0.65 |
| *C78* | Right | 0.06 | 0.15 | 1.02 | 0.47 | 0.70 | 0.75 |
| *C94* | Left | 0.00 | 0.36 | 0.10 | 0.28 | 0.49 | 0.89 |
| *C94* | Right | 0.78 | 1.25 | 0.43 | 0.49 | 0.67 | 1.59 |
| *H160* | Left | 0.65 | 0.00 | 0.16 | 3.57 | 0.55 | 0.10 |
| *H160* | Right | 0.40 | 0.52 | 0.47 | 0.34 | 0.18 | 2.50 |
| *H2024* | Left | 1.51 | 0.89 | 0.04 | 0.13 | 0.20 | 0.86 |
| *H2024* | Right | 0.01 | 5.71 | 2.15 | 3.57 | 0.47 | 0.04 |
| *H516* | Left | 0.24 | 0.20 | 1.05 | 0.09 | 0.33 | 1.83 |
| *H516* | Right | 0.19 | 0.50 | 0.42 | 0.28 | 0.12 | 0.13 |
| *H824* | Left | 0.24 | 1.33 | 0.39 | 2.97 | 2.13 | 2.02 |
| *H824* | Right | 1.02 | 1.16 | 0.98 | 0.06 | 0.18 | 0.27 |
| Mean (Left) | | 0.96 | 2.58 | 1.73 | 2.13 | 2.33 | 1.72 |
| SD (Left) | | 0.94 | 3.17 | 1.33 | 2.39 | 2.80 | 1.78 |
| Mean (Right) | | 1.00 | 1.37 | 1.23 | 1.59 | 0.97 | 0.98 |
| SD (Right) | | 0.98 | 1.65 | 1.16 | 1.76 | 1.33 | 0.92 |
| Mean (All) | | 0.98 | 1.98 | 1.48 | 1.86 | 1.65 | 1.35 |
| SD (All) | | 0.95 | 2.58 | 1.26 | 2.09 | 2.28 | 1.45 |

**Supplementary Tables S7–S11.** Fixed-effect (beta) estimates from the per-group linear mixed-effects models of vagal branch-emergence distance. A separate model was fit for each of the five multi-subgroup branch groups — Sympathetic (Table S7, ICC = 0.105), Vascular (S8, ICC = 0.117), Muscular (S9, ICC = 0.097), Cardiac (S10, ICC = 0.280), and Esophageal (S11, ICC = 0.229) — each of the form: registered distance from the jugular foramen ~ sex + side + subgroup + sex×side + sex×subgroup + side×subgroup, with a random intercept for donor to account for repeated branches within the same donor. In each table, every row is one model term: "Effect" gives the term type (intercept, a main effect, or a two-way interaction), and "Variable" and "Interaction variable" give the factor level(s) being estimated relative to the reference cell — a female donor, the left side, and the most-proximal subgroup of that group — which is fixed at 0.00. "Beta Estimate" is the coefficient in centimetres (the shift in emergence distance relative to the reference), reported with its standard error, 95% confidence limits (Lower CL, Upper CL), and p-value (Pr > |t|). The intraclass correlation coefficient (ICC), reported per group, is the fraction of total variance attributable to between-donor differences.

**Supplementary Table S7. LMM beta estimates — Sympathetic group (ICC = 0.105)**

| **Effect** | **Variable** | **Interaction variable** | **Beta Estimate** | **Standard Error** | **Lower CL** | **Upper CL** | **Pr > \|t\|** |
| --- | --- | --- | --- | --- | --- | --- | --- |
| Intercept |  |  | 3.71 | 0.51 | 2.71 | 4.71 | **<0.0001** |
| Sex | F |  | 0.00 | . | . | . | . |
| Sex | M |  | -0.07 | 0.71 | -1.45 | 1.32 | 0.9237 |
| Side | L |  | 0.00 | . | . | . | . |
| Side | R |  | 0.96 | 0.63 | -0.28 | 2.20 | 0.1310 |
| Subgroup | Superior Cervical Ganglion |  | 0.00 | . | . | . | . |
| Subgroup | Sympathetic Trunk |  | 3.94 | 0.93 | 2.12 | 5.77 | **<0.0001** |
| Sex * Side | M | R | -0.70 | 0.84 | -2.35 | 0.96 | 0.4097 |
| Sex * Subgroup | M | Sympathetic Trunk | 1.10 | 1.02 | -0.89 | 3.10 | 0.2780 |
| Side * Subgroup | R | Sympathetic Trunk | 0.10 | 0.98 | -1.82 | 2.03 | 0.9183 |

**Supplementary Table S9. LMM beta estimates — Muscular group (ICC = 0.097)**

| **Effect** | **Variable** | **Interaction variable** | **Beta Estimate** | **Standard Error** | **Lower CL** | **Upper CL** | **Pr > \|t\|** |
| --- | --- | --- | --- | --- | --- | --- | --- |
| Intercept |  |  | 3.12 | 0.76 | 1.64 | 4.61 | **<0.0001** |
| Sex | F |  | 0.00 | . | . | . | . |
| Sex | M |  | -0.73 | 0.93 | -2.56 | 1.10 | 0.4347 |
| Side | L |  | 0.00 | . | . | . | . |
| Side | R |  | 0.73 | 0.86 | -0.96 | 2.41 | 0.3960 |
| Subgroup | Pharyngeal |  | 0.00 | . | . | . | . |
| Subgroup | Laryngeal (SLN) |  | 0.35 | 0.90 | -1.42 | 2.11 | 0.6999 |
| Subgroup | Ansa/Laryngeal |  | 8.51 | 0.98 | 6.59 | 10.43 | **<0.0001** |
| Subgroup | Laryngeal (RLN) |  | 19.83 | 0.93 | 18.01 | 21.66 | **<0.0001** |
| Sex * Side | M | R | -0.77 | 0.73 | -2.20 | 0.66 | 0.2931 |
| Sex * Subgroup | M | Laryngeal (SLN) | 1.60 | 1.03 | -0.42 | 3.63 | 0.1205 |
| Sex * Subgroup | M | Ansa/Laryngeal | -1.59 | 1.14 | -3.82 | 0.64 | 0.1619 |
| Sex * Subgroup | M | Laryngeal (RLN) | 0.62 | 1.06 | -1.46 | 2.69 | 0.5600 |
| Side * Subgroup | R | Laryngeal (SLN) | -0.34 | 1.03 | -2.36 | 1.69 | 0.7461 |
| Side * Subgroup | R | Ansa/Laryngeal | -0.83 | 1.08 | -2.94 | 1.28 | 0.4411 |
| Side * Subgroup | R | Laryngeal (RLN) | -5.73 | 1.06 | -7.81 | -3.65 | **<0.0001** |

**Supplementary Table S8. LMM beta estimates — Vascular group (ICC = 0.117)**

| **Effect** | **Variable** | **Interaction variable** | **Beta Estimate** | **Standard Error** | **Lower CL** | **Upper CL** | **Pr > \|t\|** |
| --- | --- | --- | --- | --- | --- | --- | --- |
| Intercept |  |  | 7.93 | 0.89 | 6.19 | 9.67 | **<0.0001** |
| Sex | F |  | 0.00 | . | . | . | . |
| Sex | M |  | -2.56 | 1.21 | -4.93 | -0.19 | **0.0344** |
| Side | L |  | 0.00 | . | . | . | . |
| Side | R |  | -2.84 | 1.00 | -4.79 | -0.89 | **0.0044** |
| Subgroup | Internal Carotid Artery |  | 0.00 | . | . | . | . |
| Subgroup | Carotid Sinus/Bifurcation |  | -0.86 | 1.17 | -3.15 | 1.44 | 0.4649 |
| Subgroup | Internal Jugular Vein |  | 2.27 | 1.07 | 0.18 | 4.37 | **0.0335** |
| Subgroup | Common Carotid |  | 4.35 | 0.94 | 2.50 | 6.20 | **<0.0001** |
| Subgroup | General Vascular |  | 8.67 | 1.25 | 6.22 | 11.12 | **<0.0001** |
| Subgroup | Aorta |  | 15.19 | 1.22 | 12.79 | 17.58 | **<0.0001** |
| Sex * Side | M | R | 0.63 | 0.80 | -0.95 | 2.20 | 0.4356 |
| Sex * Subgroup | M | Carotid Sinus/Bifurcation | 0.37 | 1.33 | -2.24 | 2.99 | 0.7803 |
| Sex * Subgroup | M | Internal Jugular Vein | 4.24 | 1.40 | 1.50 | 6.99 | **0.0025** |
| Sex * Subgroup | M | Common Carotid | -0.27 | 1.22 | -2.65 | 2.12 | 0.8249 |
| Sex * Subgroup | M | General Vascular | 0.98 | 1.42 | -1.80 | 3.77 | 0.4889 |
| Sex * Subgroup | M | Aorta | 0.81 | 1.65 | -2.42 | 4.04 | 0.6241 |
| Side * Subgroup | R | Carotid Sinus/Bifurcation | 2.09 | 1.30 | -0.45 | 4.64 | 0.1074 |
| Side * Subgroup | R | Internal Jugular Vein | 3.07 | 1.35 | 0.42 | 5.73 | **0.0232** |
| Side * Subgroup | R | Common Carotid | 2.53 | 1.19 | 0.20 | 4.85 | **0.0330** |
| Side * Subgroup | R | General Vascular | 2.66 | 1.42 | -0.13 | 5.45 | 0.0615 |
| Side * Subgroup | R | Aorta | -1.16 | 2.30 | -5.67 | 3.35 | 0.6137 |

**Supplementary Table S10. LMM beta estimates — Cardiac group (ICC = 0.280)**

| **Effect** | **Variable** | **Interaction variable** | **Beta Estimate** | **Standard Error** | **Lower CL** | **Upper CL** | **Pr > \|t\|** |
| --- | --- | --- | --- | --- | --- | --- | --- |
| Intercept |  |  | 15.89 | 0.63 | 14.65 | 17.12 | **<0.0001** |
| Sex | F |  | 0.00 | . | . | . | . |
| Sex | M |  | -0.11 | 1.00 | -2.06 | 1.84 | 0.9130 |
| Side | L |  | 0.00 | . | . | . | . |
| Side | R |  | -0.32 | 0.85 | -1.98 | 1.35 | 0.7082 |
| Subgroup | Superficial Cardiac Plexus |  | 0.00 | . | . | . | . |
| Subgroup | General Cardiac |  | 6.83 | 1.05 | 4.77 | 8.89 | **<0.0001** |
| Subgroup | Deep Cardiac Plexus |  | 6.41 | 0.95 | 4.54 | 8.27 | **<0.0001** |
| Subgroup | Cardiopulmonary |  | 8.33 | 0.60 | 7.16 | 9.50 | **<0.0001** |
| Subgroup | Pulmonary |  | 10.85 | 0.60 | 9.68 | 12.02 | **<0.0001** |
| Sex * Side | M | R | 0.60 | 0.54 | -0.45 | 1.65 | 0.2600 |
| Sex * Subgroup | M | General Cardiac | -6.45 | 1.48 | -9.35 | -3.54 | **<0.0001** |
| Sex * Subgroup | M | Deep Cardiac Plexus | -2.84 | 1.29 | -5.37 | -0.30 | **0.0285** |
| Sex * Subgroup | M | Cardiopulmonary | -0.71 | 0.86 | -2.41 | 0.98 | 0.4103 |
| Sex * Subgroup | M | Pulmonary | -0.42 | 0.86 | -2.10 | 1.26 | 0.6245 |
| Side * Subgroup | R | General Cardiac | -1.09 | 1.64 | -4.30 | 2.12 | 0.5055 |
| Side * Subgroup | R | Deep Cardiac Plexus | -3.24 | 1.28 | -5.75 | -0.74 | **0.0112** |
| Side * Subgroup | R | Cardiopulmonary | -4.35 | 0.93 | -6.19 | -2.52 | **<0.0001** |
| Side * Subgroup | R | Pulmonary | -1.84 | 0.94 | -3.69 | 0.01 | 0.0506 |

**Supplementary Table S11. LMM beta estimates — Esophageal group (ICC = 0.229)**

| **Effect** | **Variable** | **Interaction variable** | **Beta Estimate** | **Standard Error** | **Lower CL** | **Upper CL** | **Pr > \|t\|** |
| --- | --- | --- | --- | --- | --- | --- | --- |
| Intercept |  |  | 26.65 | 0.63 | 25.42 | 27.88 | **<0.0001** |
| Sex | F |  | 0.00 | . | . | . | . |
| Sex | M |  | -2.50 | 0.84 | -4.15 | -0.85 | **0.0030** |
| Side | L |  | 0.00 | . | . | . | . |
| Side | R |  | -3.00 | 0.54 | -4.07 | -1.94 | **<0.0001** |
| Subgroup | General Esophageal |  | 0.00 | . | . | . | . |
| Subgroup | Pulmonary (Esophageal) |  | 0.21 | 0.78 | -1.32 | 1.75 | 0.7834 |
| Subgroup | Esophageal Plexus |  | 1.72 | 0.69 | 0.37 | 3.07 | **0.0123** |
| Sex * Side | M | R | 0.34 | 0.51 | -0.67 | 1.35 | 0.5073 |
| Sex * Subgroup | M | Pulmonary (Esophageal) | 1.93 | 0.91 | 0.16 | 3.71 | **0.0326** |
| Sex * Subgroup | M | Esophageal Plexus | 2.98 | 0.74 | 1.53 | 4.44 | **<0.0001** |
| Side * Subgroup | R | Pulmonary (Esophageal) | 3.35 | 0.75 | 1.88 | 4.82 | **<0.0001** |
| Side * Subgroup | R | Esophageal Plexus | 1.50 | 0.63 | 0.25 | 2.74 | **0.0184** |

**Supplementary Table S12. Classification performance of the cervical–thoracic LDA boundary**

One-dimensional linear discriminant analysis of registered distance from the jugular foramen, classifying each branch as cervical (pharyngeal-arch: sympathetic, muscular, vascular, multiple-target) versus thoracic (primitive-mediastinum: cardiac, pulmonary, esophageal), fit separately per side on the full published dataset .Thoracic is the positive class: sensitivity = proportion of thoracic branches correctly classified, specificity = proportion of cervical branches correctly classified. Boundary = LDA decision point (−intercept/coefficient). Values are resubstitution (in-sample) estimates. AUC, area under the ROC curve.

| Side | n cervical | n thoracic | Boundary (cm) | Accuracy | Sensitivity | Specificity | AUC |
| --- | --- | --- | --- | --- | --- | --- | --- |
| Left | 496 | 501 | 17.2 | 0.85 | 0.89 | 0.80 | 0.93 |
| Right | 460 | 720 | 15.1 | 0.90 | 0.95 | 0.82 | 0.97 |
| *Pooled* | *956* | *1221* | *16.0* | *0.87* | *0.92* | *0.81* | *0.95* |

**Supplementary Table S13. Side-specific zone/domain emergence contrasts (least-squares means)**

For each contrast shown in Figure 7, the effect size is the difference between adjacent zone means (each zone mean is the average of the least-squares means of its constituent subgroups) and the p-value is the least-squares-means contrast at the zone boundary, both evaluated separately for the left and right vagus nerve from the per-group linear mixed-effects models (fixed: sex, side, subgroup + two-way interactions; random intercept for donor), averaging over sex. Δ, distal minus proximal (cm). Developmental domains carry no p-value; the pulmonary→esophageal step crosses two group models and has no single-model p. n.s., not significant (p ≥ 0.05).

| Group | Contrast | Left Δ (cm) | Left p | Right Δ (cm) | Right p |
| --- | --- | --- | --- | --- | --- |
| Developmental domains | Pharyngeal Arch → Primitive Mediastinum | +13.0 | — | +12.1 | — |
| Muscular zones | Upper Cervical → Lower Cervical | +7.1 | < 0.001 | +6.5 | < 0.001 |
|  | Lower Cervical → Upper Thoracic | +12.4 | < 0.001 | +7.5 | < 0.001 |
| Vascular zones | Upper Cervical → Lower Cervical | +4.6 | < 0.001 | +6.4 | < 0.001 |
|  | Lower Cervical → Upper Thoracic | +8.1 | < 0.001 | +6.0 | < 0.001 |
| Thoracic groups | Cardiac → Cardiopulmonary | +5.1 | < 0.001 | +2.2 | < 0.01 |
|  | Cardiopulmonary → Pulmonary | +2.7 | < 0.001 | +5.2 | < 0.001 |
|  | Pulmonary → Esophageal (core) | +0.4 | — | +1.0 | — |
| Cardiac subgroups | Superficial cardiac plexus → Deep cardiac plexus | +5.0 | < 0.001 | +1.8 | n.s. (0.07) |
|  | Deep cardiac plexus → Cardiopulmonary | +3.0 | < 0.001 | +1.9 | < 0.01 |

### Data Availability

The branch distance data and the scripts to reproduce the figures, as well as a data explorer dashboard that lets users define their own comparison groups and perform their own selectivity analyses, and visualize the distributions on a 3D post mortem dissection model is available at https://github.com/drsiyarb/vagus_nerve_explorer
